# Proteomics of the astrocyte secretome reveals changes in their response to soluble oligomeric Aß

**DOI:** 10.1101/2023.01.18.523916

**Authors:** Vittoria Matafora, Alena Gorb, Wendy Noble, Angela Bachi, Beatriz Gomez Perez-Nievas, Maria Jimenez-Sanchez

## Abstract

Astrocytes associate with amyloid plaques in Alzheimer’s disease (AD). Astrocytes react to changes in the brain environment, including to increasing concentrations of amyloid-ß (Aß). However, the precise response of astrocytes to soluble small Aß oligomers at concentrations similar to those present in the human brain has not been addressed. In this study, we exposed astrocytes to neuronal media containing soluble human Aß oligomers and used proteomics to investigate changes in the astrocyte secretome. Our data shows dysregulated secretion of astrocytic proteins involved in the extracellular matrix and cytoskeletal organization and increase secretion of proteins involved in oxidative stress responses and those with chaperone activity. Several of these proteins have been identified in previous transcriptomic and proteomic studies using brain tissue from human AD and cerebrospinal fluid (CSF). Our work highlights the relevance of studying astrocyte secretion to understand the brain response to AD pathology and the potential use of these proteins as biomarkers for the disease.

## INTRODUCTION

Alzheimer’s disease (AD) pathology is classically defined by the extracellular deposition of amyloid-ß (Aß) and the intracellular accumulation of hyperphosphorylated tau, which is accompanied by neuronal and synapse loss, leading to cognitive impairment (Knopman et al. 2021). Astrocyte reactivity, involving transcriptional, morphological and functional changes in astrocytes in response to Aß and tau, is also a key feature of AD. Glial fibrillary acidic protein (GFAP)-positive astrocytes usually cluster around amyloid plaques (Itagaki et al. 1989; Serrano-Pozo et al. 2013). Astrocyte reactivity indicated by increased GFAP expression is one of the characteristics that allows AD demented cases to be distinguished from those who are clinically non-demented while having AD-like pathology (Barroeta-Espar et al. 2019; Perez-Nievas et al. 2013), and evidence suggests that this astrocyte reaction precedes Aß and tau deposition in human brain (Carter et al. 2012; Rodriguez-Vieitez et al. 2016). Recently, plasma GFAP levels have been associated with increased Aß pathology (Benedet et al. 2021; Pereira et al. 2021) and are elevated in older individuals at risk of AD (Chatterjee et al. 2021) further highlighting the relevance of astrocytes in the disease.

Astrocytes react to Aß plaques in AD, however, the specific functional changes that this reactivity causes and the implications in the onset and progression of AD remains uncertain. Exposure to extracellular Aß induces an astrocytic response in culture, as evidenced by an increase in astrocyte-mediated neurotoxicity through release of N-SMase (Jana and Pahan 2010) and soluble inflammatory factors (Garwood et al. 2011), or increased synaptotoxicity mediated by the secretion of factors such as glutamate (Talantova et al. 2013), complement C3 (Lian et al. 2015; 2016), or CXCL1 (Perez-Nievas et al. 2021). Additionally, Aß exposure may interfere with astrocytic protective functions as it results in decrease secretion of synaptogenic factors such as TSP-1 (Rama Rao et al. 2013) or TGF-ß1 (Diniz et al. 2017) and impairs phagocytosis and degradation of dystrophic neurites (Sanchez-Mico 2021).

Previous proteomics studies have characterized the astrocyte secretome in resting conditions (Dowell, Johnson, and Li 2009; Greco et al. 2010; Han et al. 2014; Lafon-Cazal et al. 2003) and in response to proinflammatory stimuli such as lipopolysaccharide (LPS) or cytokines (Lafon-Cazal et al. 2003; Delcourt et al. 2005; Keene et al. 2009), as well as specific insults including cholinergic stimulation (Moore et al. 2009), angiogenin (Skorupa et al. 2013), mechanic injury (Thorsell et al. 2008; Lai et al. 2013) or in response to endoplasmic reticulum (ER) stressors (Smith et al. 2020). The astrocyte secretome has also been characterized in response to synthetic Aß42 (Lai et al. 2013). However, much of the existing research characterizing the astrocyte response in AD is limited due to the lack of appropriate tools to mimic in culture the species and concentrations of Aß found in AD brain. Synthetic Aß peptides have been typically used at doses that exceed the physiological concentrations of Aß by 100-1000 times. In addition, the nature of the peptides is crucial to determine their underlying effects. Rather than monomers or highly aggregated forms, soluble Aß oligomers are the species typically associated with synaptic dysfunction and loss of dendritic spines (Lacor et al. 2004; Shankar et al. 2007), increased tau phosphorylation and missorting into dendrites (Zempel et al. 2010), impairment of axonal transport (Vossel et al. 2010; Sherman et al. 2016) or increased reactive oxygen speces (ROS) production (Behl et al. 1994). Investigating how astrocyte secretion is modulated in response to physiological forms of Aß will better help to understand the contribution of astrocytes to AD pathology.

In this study, we characterized changes in the astrocyte secretome in response to soluble Aβ oligomers, at concentrations and species similar to those present in the human brain, secreted into culture medium from neurons that express a human *APP* transgene with the double Swedish mutation (APPSwe, Tg2576 mice (Hsiao et al. 1996)). Using an optimized methodology to analyse the extracellular media with mass spectrometry while discriminating serum components, we generated a list of proteins whose levels are significantly changed in the astrocyte secretome upon treatment with neuron-derived Aß oligomers. We used functional and pathway annotation to explore the processes that are altered in astrocytes in these disease-mimicking conditions. Our data identified that Aβ exposed astrocytes show altered secretion of proteins involved in reorganization of the extracellular matrix and the cytoskeleton, as well as in protective antioxidant and chaperone function responses. Some of the identified proteins are altered in CSF and tissues from AD patients, supporting the idea that astrocyte-secreted proteins can be explored as potential biomarkers for disease and can aid understanding of functional astrocyte changes in AD.

## RESULTS

### Analysis of the astrocyte secretome using an in-house library shows changes in response to Aß oligomers

To investigate changes in the astrocyte secretome in response to soluble Aß oligomers similar to those found in human disease brain, we exposed primary mouse astrocytes to conditioned media of neurons derived from Tg2576 mouse embryos that express a human *APP* transgene harbouring the double Swedish mutation K670N, M671L (APPSwe, Hsiao et al. 1996) or from their wild type littermates (WT) (Figure 1A). The presence of low molecular weight Aß oligomers in Tg2576 media has been widely characterized (Wu et al. 2010; DaRocha-Souto et al. 2012; Perez-Nievas et al. 2021), with Aß42:Aß40 ratios of 1:10 as in human AD brain, and concentrations ranging between 2 and 8 nM, similar to what it has been reported in human CSF (Snider et al. 2009). Conditioned media was diluted to treat primary astrocytes with 2 nM and 0.2 nM concentration of Aß40 and Aß42 oligomers, respectively. Astrocytes were exposed to APPSwe neuron conditioned media (APPSwe_NCM) or wild type neuron conditioned media (WT_NCM) and the astrocyte conditioned media was collected after 24h. The conditioned media from stimulated astrocytes (APPSwe_ACM) or control (WT_ACM) were subjected to mass spectrometry, together with APPSwe_NCM or WT_NCM (Figure 1A), to control for the components that were already present in neuron media. Importantly, exposure to Aß containing media at the concentration and time used in this study did not result in astrocyte cell death, as we previously reported (Perez-Nievas et al. 2021), and therefore altered cellular viability is not a confounding factor in the proteomic analysis.

**Figure 1.**
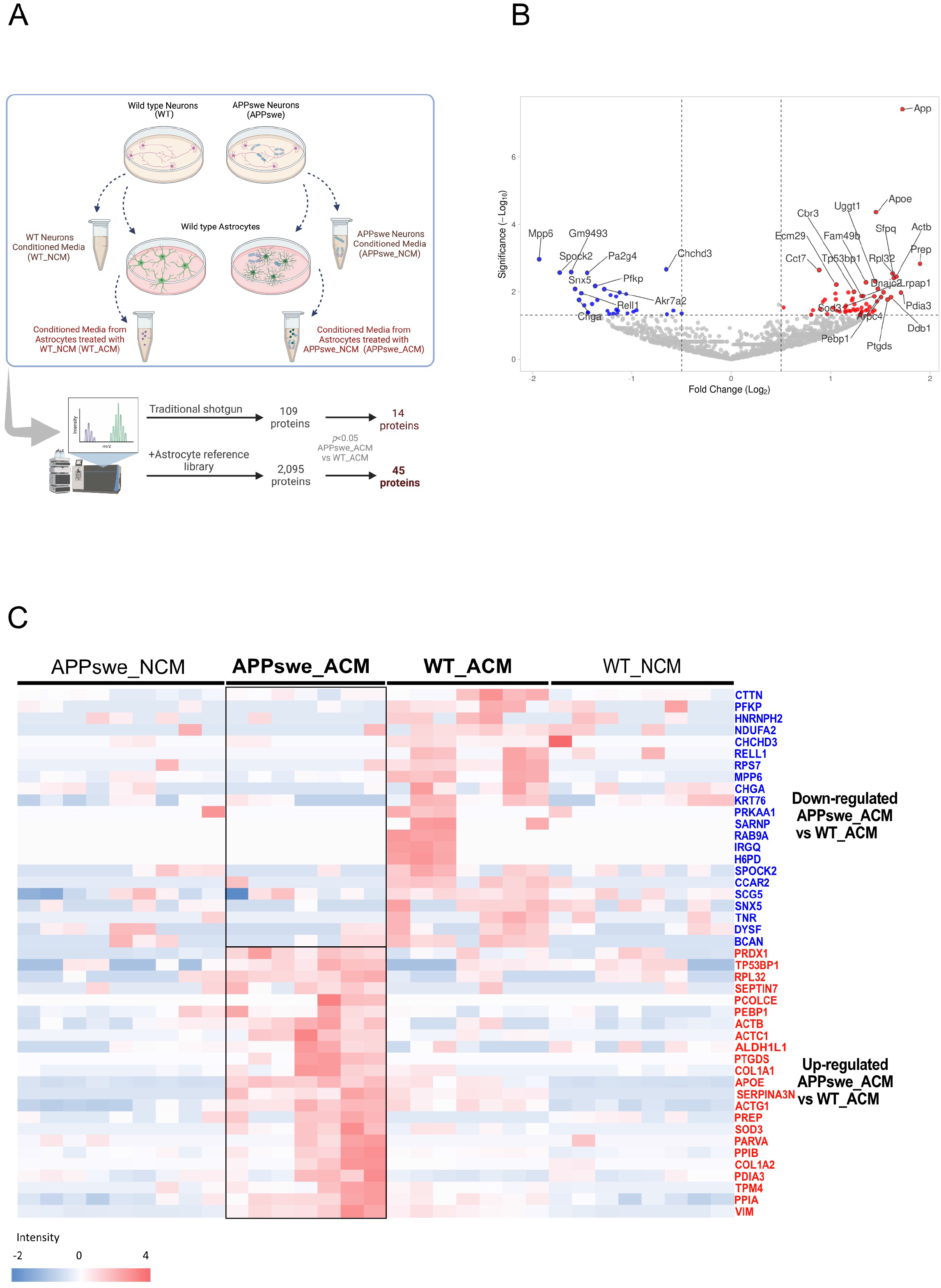
Proteomics analysis of the astrocyte conditioned media in response to Aß-containing media. **(A)** Schematic representation of the experimental approach. Conditioned media (NCM) from primary neurons prepared from Tg2576 mice (APPswe) or wild-type littermates (WT) mice were collected at DIV14. Wild-type astrocytes were treated with either APPswe conditioned media (APPswe_NCM) or WT conditioned media (WT_NCM), and resultant astrocyte-conditioned media (ACM) was analysed by mass spectrometry. Traditional shotgun proteomics identified 109 proteins differentially secreted by astrocytes treated with Aβ-containing media, while referencing to an astrocyte library increased the yield to 2,095 proteins. Created with BioRender.com. **(B)** Volcano plot showing proteins significantly up (red) and downregulated (blue) in the secretome of Aβ-stimulated astrocytes, APPswe_ACM, compared to control, WT_ACM, before proteins contained in neuron media were excluded. The significance threshold was set at ≥ 1.3 and the absolute fold change threshold was set at > 0.5. **(C)** Heatmap of proteins differentially secreted from Aβ-stimulated astrocytes. The blue-to-red scale indicates low-to-high protein levels. Differentially expressed proteins were defined based on a *p*<0.05.

An initial analysis using traditional shotgun proteomics identified 109 proteins, 14 of which were significantly altered in the secretome of astrocytes exposed to APPswe neuron conditioned media (APPSwe_NCM) compared to wild type (WT_NCM) (data not shown). Proteomics of the secretome is usually limited by the high abundance of serum proteins, which creates an excessive dynamic range between albumin and other proteins contained in the media. Our experimental design required the use of serum-like supplement to ensure neuron survival, and this contains albumin and other proteins at high abundance (Brewer et al. 1993). Albumin immunodepletion strategies were not considered to avoid the nonspecific loss of proteins of interest (Bellei et al. 2011). To overcome these limitations, we prepared an inhouse library of proteins contained in the conditioned media of astrocytes grown in media without serum for 24h, and used it as a matching library following a previously published protocol (Geyer et al. 2016).

We identified 2,095 proteins in the secretome of astrocytes treated with media from 5 independent neuronal cultures (Supplementary table 1). Figure 1B shows a volcano plot comparing the secretome from astrocytes treated with media from APPswe neurons (APPswe_ACM) with that of astrocytes treated WT neurons (WT_ACM), which rendered a list of 94 significantly altered proteins (Supplementary table 1). Because astrocytes were treated with media that contains other neuron secreted proteins in addition to Aß oligomers we excluded those proteins that were found both in APPSwe_NCM or WT_NCM respectively (Supplementary table 1). This resulted in a list of 45 astrocyte secreted proteins that were affected by exposure to oligomeric Aβ, being differentially expressed in APPSwe_ACM compared to WT_ACM (Figure 1C). Of these identified proteins, 23 and 22 proteins were significantly increased or decreased, respectively, when astrocytes were treated with media from APPswe neurons containing Aß oligomers (Table 1). Among these proteins increased, we identified known markers of reactive astrocytes such as APOE, vimentin and SERPINA3N (Escartin et al. 2021; Zamanian et al. 2012), whose expression increases in astrocytes in human AD brains (Viejo et al. 2022), demonstrating the disease relevance of the proteins identified.

**Table 1.** List of proteins proteins showing significant differences in the secretome of astrocytes treated with WT_NCM vs APPSWE_NCM (p value <0.05).

| Up-Regulated proteins |  | Log<sub>10</sub>(P-value) |
| --- | --- | --- |
| ACTB | Actin, cytoplasmic 1; Beta-actin | 1.6266 |
| ACTC1 | Actin, alpha cardiac muscle 1 | 1.8713 |
| ACTG1 | Actin, cytoplasmic 2 | 2.5402 |
| ALDH1L1 | Aldehyde dehydrogenase 1 family, member L1 | 1.6360 |
| APOE | Apolipoprotein E | 1.3224 |
| COL1A1 | Collagen alpha-1(I) chain | 1.9852 |
| COL1A2 | Collagen alpha-2(I) chain | 2.4477 |
| PARVA | Alpha-parvin | 1.7085 |
| PCOLCE | Procollagen C-endopeptidase enhancer 1 | 1.4432 |
| PDIA3 | Protein disulfide-isomerase A3 | 1.7630 |
| PEBP1 | Phosphatidylethanolamine-binding protein 1 | 1.3937 |
| PPIA | Peptidyl-prolyl cis-trans isomerase A | 4.3614 |
| PPIB | Peptidyl-prolyl cis-trans isomerase B | 1.3301 |
| PRDX1 | Peroxiredoxin-1 | 1.9455 |
| PREP | Prolyl endopeptidase | 2.8245 |
| PTGDS | Prostaglandin-H2 D-isomerase | 1.8604 |
| RPL32 | Ribosomal protein L32 | 1.3087 |
| SEPTIN7 | Septin-7 | 1.4654 |
| SERPINA3N | Serine protease inhibitor A3N | 1.4466 |
| SOD3 | Extracellular superoxide dismutase [Cu-Zn] | 1.9752 |
| TPM4 | Tropomyosin alpha-4 chain | 1.4215 |
| TP53BP1 | Tumour suppressor p53-binding protein 1 | 1.5400 |
| VIM | Vimentin | 1.4315 |
| Down-Regulated proteins |  |  |
| BCAN | Brevican core protein | 1.8948 |
| CCAR2 | Cell cycle and apoptosis regulator protein 2 | 1.3478 |
| CHCHD3 | MICOS complex subunit Mic19 | 2.6624 |
| CHGA | Chromogranin-A | 1.7527 |
| CTTN | Src substrate cortactin | 1.5946 |
| DYSF | Dysferlin | 1.4324 |
| H6PD | GDH/6PGL endoplasmic bifunctional protein | 1.3870 |
| HNRNPH2 | Heterogeneous nuclear ribonucleoprotein H2 | 1.4309 |
| IRGQ | Immunity-related GTPase family, Q | 1.3891 |
| KRT76 | Keratin, type II cytoskeletal 2 oral | 1.7513 |
| MPP6 | Membrane protein, palmitoylated 6 | 2.9662 |
| NDUFA2 | NADH:Ubiquinone Oxidoreductase Subunit A2 | 1.4054 |
| PFKP | ATP-dependent 6-phosphofructokinase, platelet type | 2.1682 |
| PRKAA1 | 5'-AMP-activated protein kinase catalytic subunit alpha-1 | 1.3570 |
| RAB9A | Ras-related protein Rab-9A | 1.3923 |
| RELL1 | RELT-like protein 1 | 1.9533 |
| RPS7 | 40S ribosomal protein S7 | 2.5822 |
| SARNP | SAP domain-containing ribonucleoprotein | 1.3805 |
| SCG5 | Neuroendocrine protein 7B2 | 1.4704 |
| SNX5 | Sorting nexin-5 | 2.0812 |
| SPOCK1 | Testican-2 | 2.5676 |
| TNR | Tenascin-R | 1.3355 |

\* - Log<sub>10</sub>(p-value) > 1.3 is equivalent to p-value < 0.05.

### Over two-thirds of Aß-modulated astrocyte-secreted proteins follow non-conventional secretory pathways

Prediction tools for subcellular localization based on experimental evidence such as DeepLoc, LAGO terms or ProtComp, predicted that only 26-35% of the differentially secreted proteins are typically found in the extracellular space (Figure 2A-C). In line with this, 35% of the differentially regulated proteins contained a secretory signal for conventional secretion, while 26% of the proteins were predicted by SecretomeP to be secreted through alternative pathways (Figure 2D-F). According to the Vesiclepedia database (Pathan et al. 2019), 75% of these proteins have been previously identified as being released in extracellular vesicles (EV) (Figure 2G). DeepTMHMM online software (Hallgren et al. 2022) predicted that 3 proteins: dysferlin (DYSF), hexose-6-phosphate dehydrogenase (H6PD) and RELT-like protein 1 (RELL1), contain transmembrane (TM) helices and therefore could be secreted through ectodomain shedding (Figure 2H). Collectively, this combination of eight complementary signal peptide and localisation prediction tools, revealed that all proteins, except three, CCAR2, RAB9a and SARNP, are either predicted to be localized in the extracellular space or to be secreted (Figure 2I and Supplementary Table 2 and 3). For those secreted proteins, only one-third are expected to be released via classical secretory pathways, while most of them will require non-classical routes, such as release in EVs.

**Figure 2.**
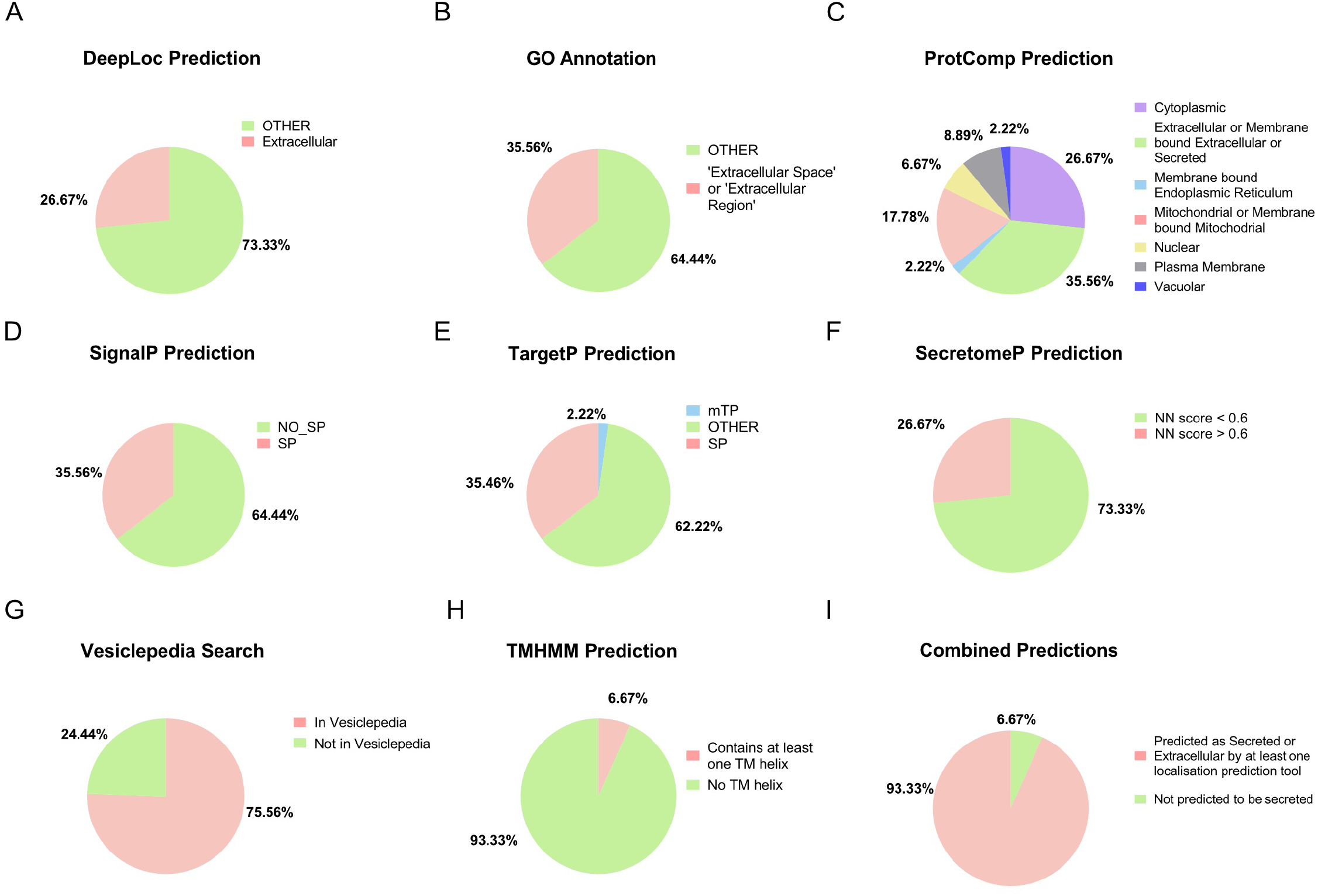
Protein localisation and secretion prediction. Different bioinformatics tools **(A-H)** and combined results **(I)** were used to predict the localisation and route of secretion of proteins identified in ACM. For SecretomeP prediction **(F)** the NN-score > 0.6 indicates that the mammalian protein is likely to be non-classically secreted. Abbreviations: SP, signal peptide; mTP, mitochondrial transit peptide; NN, non-classic secretion score; TM, transmembrane.

### Proteins related to extracellular matrix organization, cytoskeleton, response to oxidative stress and chaperone functions are overrepresented in Tthe Aß-induced astrocyte secretome

To identify astrocytic pathways and processes that could be affected by alterations in the astrocyte secretome upon exposure to Aß oligomer containing media, we performed functional annotation clustering using a combination of Gene Ontology terms (Biological Process, Cellular Component and Molecular Function), Pathways (KEGG, Reactome, and WikiPathways) and manual literature searches. Analysis of the proteins showing increased secretion upon Aß treatment (Figure 3A, 3B, Supplementary Figure 1 and Supplementary Tables 4-6) revealed an overrepresentation of proteins involved in the organization of the extracellular matrix (ECM), mostly related to the synthesis of collagen I, the main component of the ECM. Secreted proteins included collagen α-1 and −2, which together make the type I procollagen (Ricard-Blum 2011), and enzymes that catalyse the posttranslational modifications necessary to form the collagen triple helix, including the peptidyl-prolyl isomerase PPIB (Cabral et al. 2014) and the procollagen C proteinase enhancer, PCEP1 or PCOLCE (Adar et al. 1986). PREP, an prolyl endopeptidase involved in the degradation of collagen was also secreted upon Aß exposure (Gaggar et al. 2008).

**Figure 3.**
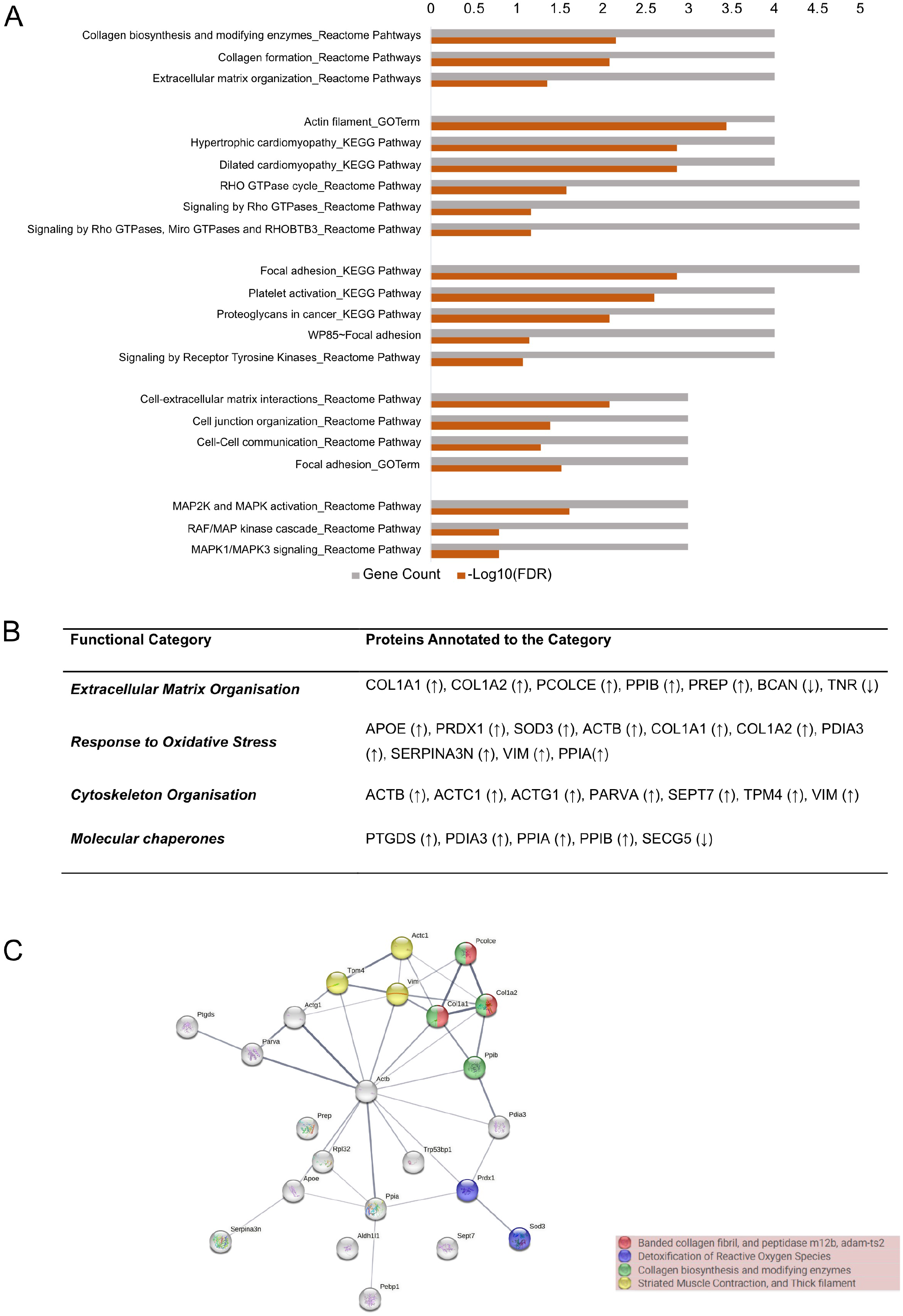
Bioinformatics analysis of proteins differentially secreted in astrocytes treated with Aß-oligomers containing media. **(A)** Functional annotation clustering analysis of differentially upregulated proteins using Database for Annotation, Visualization and Integrated Discovery (DAVID). The analysis included gene ontology (GO) and pathway (KEGG pathway, Reactome, and WikiPathways) terms. Only clusters with an enrichment score of ? 1.3 were considered significant. **(B)** Table summarising the functional categories of proteins differentially secreted by Aß-treated astrocytes. The categories were chosen and assigned based on the bioinformatics analysis and manual protein annotation. Arrows in the brackets next to the gene names indicate whether the protein was found differentially up-(↑) or down-regulated (↓). **(C)** Protein-protein interaction network for differentially upregulated proteins generated using STRING. Lines represent protein-protein associations (including but not limited to physical binding) and different thicknesses represent confidence in the interaction. Thicker lines represent higher confidence in the interaction. Circles represent individual proteins, and circles of the same colour represent proteins that were identified to be involved in the same pathway using STRING local network cluster analysis.

Although generally functioning as intracellular proteins, a high proportion of cytoskeletal components were among the proteins found secreted in response to Aß containing media. These included three out of the six known actin isoforms, ß- and γ-cytoplasmic actins (ACTB and ACTG1) and cardiac actin (ACTC1), as well as the actin-binding proteins tropomyosin α-4 chain and α-parvin, septin 7, and the astrocyte-specific intermediate filament vimentin (Figure 3A, 3B, Supplementary Figure 1 and Supplementary Table 4-6). Some of these are important structural components of focal adhesions that mediate contacts with the ECM.

Gene ontology analysis and literature searches also revealed an increase in proteins related to antioxidant activity and in the response to oxygen-containing compounds among the proteins with increased secretion (Figure 3B, Supplementary Figure 1 and Supplementary Table 5-6). Some of these proteins are known to be secreted from astrocytes including SOD3 (Iitsuka et al. 2012; Ioannou et al. 2019), APOE (Miyata and Smith 1996; Kockx et al. 2018) and SERPINA3N (Wang et al. 2019; Sánchez-Navarro et al. 2021). While not previously associated with astrocytes, peroxiredoxin-1 (PRDX1) is an intracellular enzyme with antioxidant activity reported to be secreted by tumour cells (Chang et al. 2005; 2006); and prolyl isomerase (PPIA), although cytoplasmic, is secreted as a defence mechanism in response to oxidative stress (Nigro et al. 2013). The observed changes in collagen type I may also result from a transcriptional response to oxidative stress (Martins et al. 2021).

Similar findings were obtained when using STRING to identify known and predicted physical and functional protein-protein interactions (PPIs) (Szklarczyk et al. 2021), which resulted in a significant PPI enrichment in differentially upregulated proteins (*p*-value=3.08 × 10^-11^, n=23). STRING local network clustering analysis revealed proteins involved in collagen biosynthesis and detoxification of reactive oxygen species (Figure 3C and Supplementary Table 7).

A comparison with the 332 proteins comprising the human chaperome (Brehme et al. 2014), identified proteins with reported chaperone activity, representing more than 20% of all proteins showing increased secretion upon treatment with Aß oligomer containing media (Figure 3B). These included prostaglandin synthase (PTGDS), peptidyl-prolyl isomerases A and B (PPIA and PPIB) and protein disulfide isomerase (PDIA3). In addition, further literature searches identified one protein with chaperone activity, secretogranin 5 (SECG5), in the proteins with lowered secretion. Aß-chaperone function has been reported for 3 out of 5 of these proteins: PTGDS, PDIA3 and SECG5 (Kanekiyo et al. 2007; Helwig et al. 2013; Kannaian et al. 2019; Eninger et al. 2022), suggesting that astrocytes are a source of extracellular chaperones, whose secretion is likely modulated by exposure to Aß to protect from its toxicity.

Amongst the other proteins showing decreased secretion in response to Aβ, Gene Ontology analysis revealed a significant overrepresentation of proteins, such as brevican core protein (BCAN) and tenascin-R (TNR), that are major components of perineuronal nets, specialized forms of condensed extracellular matrix that surround groups of neurons and synapses to modulate synaptic plasticity and protect neurons (Wen et al. 2018; Reichelt et al. 2019) (Supplementary Figure 2A, 2B and Supplementary Table 8). Down-regulated proteins constitute a more heterogeneous group, with no significant clusters found in DAVID and no significant PPI enrichment in STRING (*p*-value=0.172, n=22) (Supplementary Figure 2C).

Altogether, gene ontology and pathway analyses suggests that exposure of astrocytes to Aß containing medium alters the secretion of proteins involved in extracellular matrix organization and cell-extracellular matrix interactions, with enhanced secretion of proteins involved in the synthesis of collagen fibres and decreased secretion of components of specialized perineuronal structures in the extracellular matrix. In addition, our study shows an increased secretion of proteins related to the cytoskeleton and response to oxidative stress and proteins with chaperone function.

### Comparison of the Aß-stimulated astrocyte secretome with astrocyte transcriptomics and CSF proteomics studies in AD

In recent years, single-nucleus transcriptomics has identified transcriptional changes that occur specifically in astrocytes in the human AD brain. To put our results in context with human data, we compared the list of proteins that we generated with the astrocyte-specific genes that were found to be differentially expressed in early-vs late-stage human AD brain (Mathys et al. 2019), in AD vs control brain (Grubman et al. 2019; Lau et al. 2020; Sadick et al. 2022) and showing a positive correlation with Aß or phospho-tau pathology (Smith et al. 2021). We also included a comparison with a study where astrocyte-specific gene clusters were applied to 766 whole brain transcriptomes, including control, mild cognitive impairment (MCI) and demented (AD) cases (Galea et al. 2022) (Supplementary Table 9). After selecting those genes that encode proteins predicted to be secreted, we searched for those that were differentially regulated by Aβ in our study (Figure 4A). Despite the great degree of variability among published RNAseq datasets, some astrocyte secreted proteins are consistently found in AD transcriptomic studies. For example, ALDHL1 or PEBP1, whose secretion is increased upon Aß treatment, were found upregulated in AD in 3 out of 7 studies (Figure 4A). APOE, ACTB or PFKP are consistently found in nearly all transcriptomic studies, but these were reported as either up- or down-regulated (Figure 4A). BCAN is found in 4 out of 7 comparisons as upregulated in AD, while its secretion from astrocytes is decreased in our study, suggesting a complex role in the regulation of this protein in AD brain.

**Figure 4.**
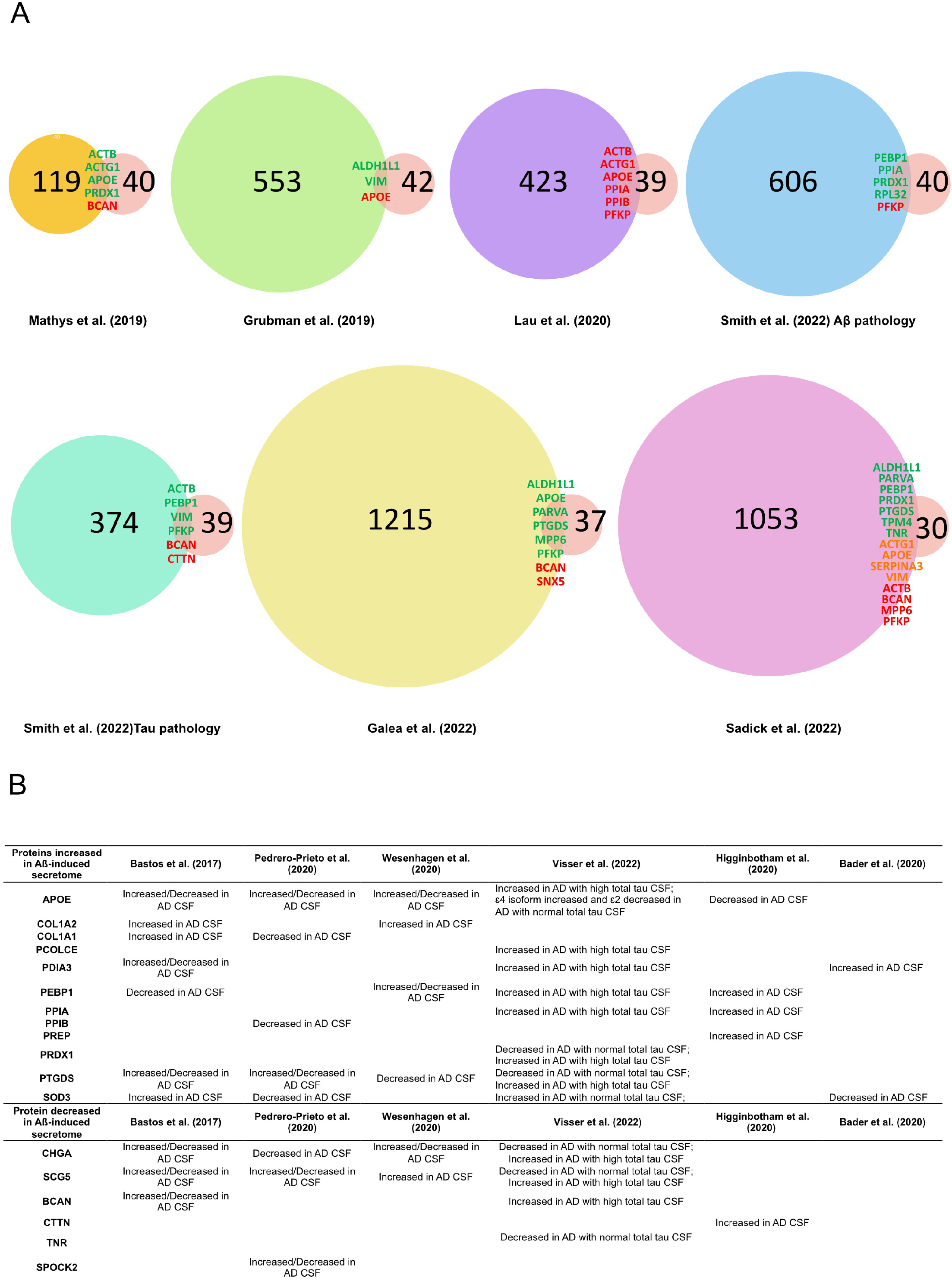
Comparison with transcriptomic and cerebrospinal fluid (CSF) Alzheimer’s disease studies. **(A)** Venn diagrams showing the numbers of proteins that overlap between this study and other transcriptomic studies of human AD samples. The number of proteins from transcriptomic studies only includes the proteins that were predicted to be extracellular or secreted using the combination of localisation prediction tools used in this study. When changes in gene expression in AD brain compared to controls were in the same direction as the secretion of proteins in response to Aß oligomers relative to control, names are shown in green; when changes were in the opposite directions, names are shown in red; when genes were reported to be both up- and downregulated, the names are shown in orange. **(B)** Table displaying the proteins that were identified in this study and that were also differentially expressed in CSF from AD compared to controls.

To further explore whether changes in protein secretion from astrocytes may reflect changes in CSF in AD, we compared our data with three systematic reviews as well as with three recent proteomic studies, which characterized the proteomic composition of CSF in AD patients compared to controls (Bastos et al. 2017; Pedrero-Prieto et al. 2020; Wesenhagen et al. 2020; Higginbotham et al. 2020; Bader et al. 2020; Visser et al. 2022) (Figure 4B). Just over half of the proteins showing increased secretion in response to media containing Aß oligomers (APOE, COL1A1, COL1A2, PCOLCE, PDIA3, PEBP1, PPIA, PPIB, PREP, PREDX1, PTGDS, SOD3) were identified as altered in AD CSF. A smaller proportion of downregulated proteins, 6 out of 22 proteins (CHGA, SCG5, BCAN, CTTN, TNR, SPOCK2), appeared as modified in AD CSF in previous studies (Bastos et al. 2017; Pedrero-Prieto et al. 2020; Wesenhagen et al. 2020; Higginbotham et al. 2020; Visser et al. 2022). Many of the identified proteins, changes are not always consistent across CSF analyses, and increased or decreased levels have been simultaneously reported in different studies. APOE is consistently found in nearly all CSF proteomic studies but, similar to the transcriptomic analysis, its levels are found up or downregulated depending on the study PEBP1, PTGDS, SOD3, CHGA and SCG5 have been repeatedly identified (4 out of 10 studies) when monitoring for changes in CSF in AD relative to controls (Table 4B). In fact, chromogranins and secretogranins have been proposed as AD biomarkers with decreased levels in CSF in AD (Abdi et al. 2006; Hölttä et al. 2015; Brinkmalm et al. 2018). Our data suggest that a proportion of the proteins that are commonly found in CSF in AD might have an astrocytic origin.

We used DisGeNET, an online database that compiles information from curated resources, GWAS catalogues, animal models and published papers (Piñero et al. 2021) to estimate the gene-disease association (GDA) score for genes encoding differentially secreted proteins in our study. 21 out of 44 (47.7%) genes were found to be associated with AD (Supplementary Figure 3 and Supplementary Table 10). Only *APOE* showed a strong association, with a GDA score of 0.7, while the rest of the genes had GDAs ≤ 0.1, consistent with the notion that, with the exception of *APOE* and *TREM*, most AD-linked genes individually have only very modest effect sizes (Escott-Price et al. 2017). While the genes encoding the other proteins have not been formally associated with AD, our data and further comparison with transcriptomic and CSF proteomics studies have identified a new set of proteins that provide insights into the likely functional consequences of astrocyte secretions in AD.

## DISCUSSION

Astrocytes are secretory cells that, in response to brain injury, disease or signals derived from other cell types, release a myriad of factors that modulate neuronal and non-neuronal cells and lead to multiple functional responses (Sofroniew 2020; Verkhratsky et al. 2016). In Alzheimer’s disease, astrocytes become reactive, partly due to their exposure to different Aß species present in the extracellular environment (reviewed in Perez-Nievas and Serrano-Pozo 2018). To our knowledge, only one study has examined the secretory response of astrocytes to Aß and they used synthetic Aß 42 in the μM range (Lai et al. 2013). Our work sought to understand how the secretory profile of astrocytes is altered in response to naturally occurring soluble oligomeric Aß, in similar nanomolar concentrations and oligomeric species to those found in AD brain (DaRocha-Souto et al. 2012; Hudry et al. 2012; Wu et al. 2012). While we cannot exclude additional effects due to other human APP-derived fragments or to proteins secreted from APPswe expressing neurons, we believe that our experimental setting closely mimics the brain environment to which astrocytes are exposed in AD. We found changes in 45 proteins, of which 23 showed higher secretion and 22 lower secretion upon exposure of astrocytes to a mixture of 1:10 Aß42: Aß40. Our study shows some similarities with Lai et al. (2013), with two proteins, APOE and prostaglandin H2 D-isomerase, identified in both datasets. However, our secretory profile differed significantly from that of astrocytes treated with synthetic Aß42 (Lai et al. 2013). These differences are not unexpected. Unlike synthetic Aß, which generates oligomers and fibrils easily within minutes to hours, especially when used at high concentrations, naturally secreted Aß peptides assemble into low molecular weight species (dimers to tetramers). These small soluble oligomeric Aß species are the most bioactive and have synaptotoxic activity (Shankar et al. 2008; T. Yang et al. 2017), and astrocytes treated with these are more likely to reflect their responses in AD brain.

A high proportion of proteins identified in this study are important for the organization of the extracellular matrix (ECM). Alterations in ECM composition have been reported in several neurodegenerative diseases, including AD (Ma et al. 2020; Freitas et al. 2021). In relation to astrocytes, a transcriptomic analysis comparing AD brains with controls identified specific clusters of astrocytes with upregulation of genes involved in ECM organization (Smith et al. 2021). Astrocytes participate in the formation of the extracellular matrix by secreting a number of factors, including proteoglycans and tenascins (Anwar et al. 2021) and, when reactive or upon Aß treatment, astrocytes have been implicated in ECM degradation through the release of matrix metalloproteinases (Deb et al. 2003; Muir et al. 2002). Our data also support this active role of astrocytes as modifiers of the ECM composition and function in AD. Of particular interest is the decreased secretion of two of the main components of the perineuronal net (PNN), brevican and tenascin R. PNNs are lattice-like assemblies of ECM proteins that surround neurons to protect them and modulate neuronal activity (Reichelt et al. 2019; Wen et al. 2018). Evidence from post-mortem brain studies suggests loss of PNNs in AD (Baig, Wilcock, and Love 2005) although other studies found preservation of these structures in disease (Brückner et al. 1999; Lendvai et al. 2013; Morawski et al. 2010). Our data suggest that astrocytes exposed to pathological Aß may decrease the secretion of PNN components, which can lead to a reduced stabilization of synapses and enhanced exposure to oxidative stress and other toxic molecules, including Aß itself (Miyata et al. 2007; Suttkus et al. 2012).

Many of the up-regulated proteins found in this study have antioxidant activity and/or are involved in the response to oxygen-containing compounds. Sings of oxidative damage are observed in aged and AD brains (Wang et al. 2014), starting in early phases of the disease (Keller et al. 2005; Butterfield et al. 2006). Some of the proteins we found dysregulated in this study have been previously linked to oxidative stress and AD. Peroxiredoxin-1, which catalyses the reduction of hydrogen peroxide and similar compounds, was found up-regulated in the cortex of AD patients (Szeliga 2020), and a recent large-scale proteomic study discovered that peroxiredoxin-1 levels were elevated in both AD brain tissues and CSF (Johnson et al. 2020). Moreover, peroxiredoxin-1 conferred resistance to Aß toxicity in PC12 cells, SH-SY5Y cells and rat primary hippocampal neurons (Cimini et al. 2013; Cumming et al. 2007). Extracellular superoxide dismutase 3 (SOD3) is another antioxidant enzyme (Wang et al. 2018) that ameliorates the oxidative damage exerted by Aß25-35 in SH-SY5Y (Yang et al. 2017), and its levels in CSF are frequently changed in AD (Bader et al. 2020; Perrin et al. 2011; Ringman et al. 2012; Visser et al. 2022). In cultured astrocytes, Aß triggers the production of reactive oxygen species (ROS) (Abramov et al. 2004; Askarova et al. 2011). The astrocyte response we report here may represent a self-defence mechanism to protect themselves against the oxidative stress caused by Aß or a mechanism to provide antioxidant support to neurons (Jiwaji and Hardingham 2022).

We also report an increased secretion of molecular chaperones. While typically considered intracellular proteins, chaperones can be secreted through classical and non-classical secretion mechanisms with important roles in extracellular proteostasis (Chaplot et al. 2020). Prostaglandin synthase is found in plaques in AD brain and its Aß chaperone activity has been reported, through both inhibiting Aß aggregation and promoting fibril disaggregation (Kanekiyo et al. 2007; Kannaian et al. 2019). PDIA3 is a disulfide isomerase with a thiol oxidoreductase activity that promotes protein folding in the ER (Chichiarelli et al. 2022). It was detected in human CSF bound to Aß (Erickson et al. 2005) and, more recently, Di Risola et al. have reported that treating SHSY5Y neuroblastoma cells with synthetic Aß led to increased extracellular PDIA3, which can bind to Aß 25-35 *in vitro* to reduce aggregation and counteract its cellular toxicity (Di Risola et al. 2022). While not previously included within the chaperone network (Brehme et al. 2014), secretogranin 5, also known as 7B2, whose levels are reduced in our Aß exposed astrocyte secretome, has also been suggested to function as a chaperone, since it colocalizes with Aß plaques in human AD brain and prevents fibrillation of Aß 40 and Aß 42 *in vitro* and blocks their cytotoxic effect in culture (Helwig et al. 2013). Single cell RNAseq and immunohistochemical studies have frequently reported that genes involved in proteostasis and chaperone-mediated responses are up-regulated specifically in astrocytes in AD (Grubman et al. 2019; Lau et al. 2020; Smith et al. 2021; Viejo et al. 2022) which, together with our study, may imply that astrocytes are a source of extracellular chaperones to protect the disease brain environment.

Three ubiquitously expressed actin isoforms are amongst the secreted proteins in response to Aß. Different roles of actin as an extracellular protein have been reported (Sudakov et al. 2017), including its function as a “danger-associated molecular pattern” (DAMP) (Ahrens et al. 2012; Srinivasan et al. 2016). DAMPs trigger an inflammatory response by binding to receptors such as Toll-like receptors, which are expressed mostly in microglia and astrocytes in the brain (Sofroniew 2020). Other proteins found in our study have been suggested as DAMPs, including peroxiredoxin 1 (PRDX1), which binds to TLR-4 after secretion from cancer cells (Liu et al. 2016; Riddell et al. 2010), or prolyl endopeptidase (PREP) that cleaves collagen to produce proline-glycine-proline (PGP), which acts as a DAMP (Patel and Snelgrove 2018) and participates in the recruitment of neurotrophils (Weathington et al. 2006). In addition, other proteins upregulated in the Aß-induced astrocyte secretome have been directly implicated in inflammation. Prostaglandin D synthase (PTGDS) catalyzes the conversion of prostaglandin H2 to prostaglandin D2, an arachidonic acid metabolite which functions as a modulator of the inflammatory response (Joo and Sadikot 2012); peptidylprolyl isomerase A (PPIA) is mainly cytoplasmic but it can also be secreted in response to different inflammatory stimuli such as LPS (Hoffmann and Schiene-Fischer 2014), and inhibition of extracellular PPIA reduces neurotoxicity and neuroinflammation markers such as NF-kB activation in a mouse model of amyotrophic lateral sclerosis (ALS) (Pasetto et al. 2017). Altogether, a large proportion of the secreted proteins identified in response to Aß oligomers may be involved in further amplifying or diminishing the inflammatory response initiated by Aß. Surprisingly, we did not identify any cytokines or chemokines. Cytokines and chemokines are typically secreted from astrocytes, and their secretion profile is altered in AD brain and more specifically upon treatment with Aß (González-Reyes et al. 2017), including our own studies using a similar experimental setting (Perez-Nievas 2021). Cytokine and chemokine secretion is typically detected by immunoassay, while its detection through mass spectrometry is challenging due to their low molecular weight and low abundance relative to other proteins in the media (Kupcova Skalnikova et al. 2017), which may explain their absence in this dataset.

Proteins showing reduced secretion in this study constitute a more heterogeneous group. Amongst these proteins, we identified proteins related to mitochondrial integrity and function (NDUFAD, CHCHD3), glucose metabolism (H6PD, PFKP), energy sensors (PRKAA1, SARNP), RNA binding proteins (HNRNPH2), intracellular trafficking (RAB9A, SNX5), as well as secreted neuroendocrine peptides (CHGA, SCG5).

Astrocyte proteins with increased secretion in response to Aß largely overlapped with proteomics studies conducted in AD CSF. Increased levels of astrocytic proteins in CSF, such as GFAP, have been associated with Aß deposition, levels of phosphor/total tau and with markers of synaptic dysfunction (Milà-Alomà et al. 2020; Salvadó et al. 2021). In fact, a CSF proteomic study in the APPPS1 mouse model of amyloidosis highlighted increased levels of proteins related to glial activation and identified APOE and SERPINA3 as CSF biomarkers (Eninger et al. 2022). Our data suggest that some of the proteins that are commonly found in CSF in AD, might be of astrocytic origin, and that changes in astrocytes can be reflected in the CSF. This may be the case for some proteins already proposed as AD biomarkers such as chromogranins (Abdi et al. 2006; Hölttä et al. 2015; Brinkmalm et al. 2018).

To conclude, our study shows that in response to human Aß oligomers, in similar concentrations and species to those found in AD brain, astrocytes present a distinct profile encompassing ECM molecules, antioxidant proteins, components of the cytoskeleton, chaperones, and regulators of inflammatory responses. These changes mimic some of those identified in AD transcriptomic and CSF studies and contribute to a better understanding of how astrocytes play a pivotal role in the molecular response to Aß in AD and can be relevant for the identification of novel biomarkers.

## METHODS

### Animals

All animal work was conducted in accordance with the UK Animals (Scientific Procedures) Act 1986 and the European Directive 2010/63/EU under UK Home Office Personal and Project Licenses and with agreement from the King’s College London (Denmark Hill) Animal Welfare and Ethical Review Board. Pregnant female CD1 mice purchased from Charles River were used to prepare mouse primary astrocyte cultures. Tg2576 mice were used to prepare primary neurons that secrete human A Aß. Tg2576 mice overexpress human APP (isoform 695) containing the double mutation K670N, M671L (Swedish mutation) under the control of the hamster prion protein promoter (Hsiao et al. 1996). Tg2576 mice were originally obtained from Taconic farms (Germantown, NY, USA), and were maintained in-house by breeding males with C57Bl/6/SJL F1 females as recommended by the suppliers. The genotype of the animals was determined by polymerase chain reaction on DNA obtained from the embryos, as previously described (Mitchell et al. 2009). Water and food were available (Picolab rodent diet 20; # 5053; Lab Diet, St Louis, MO, USA) ad libitum. Animals were housed at 19–22 °C, humidity 55%, 12-h:12-h light:dark cycle with lights on at 07:30.

### Culture of primary cortical neurons and collection of neuronal conditioned media

To obtain oligomeric Aß enriched conditioned media, we cultured neurons from cerebral cortex of Tg2576 mice bearing the human APPSwe mutation at embryonic day 15 (E15) as previously described (Wu et al. 2010; DaRocha-Souto et al. 2012; Perez-Nievas et al. 2021). Primary cortical neurons were isolated from E16.5 embryos resulting from the mating between a Tg2576 male that heterozygously overexpresses a human mutated *APP* gene and a wild-type female, thus giving rise to both transgenic (APPswe) and littermate (WT) cultures. The expression of the APPswe gene was determined for each embryo. Briefly, brains were harvested and placed in ice-cold HBSS with HEPES, where the meninges were removed, and the cerebral cortices were dissected and digested in TryplE. Cells were cultured in polyD-lysine coated plates and grown in Neurobasal medium supplemented with 2% B27, GlutaMAX™, sodium pyruvate, and 100 units/ml penicillin and 100 μg/ml streptomycin at 37 °C in a humidified incubator with 5% CO2. After 14 DIV, media from transgenic mice (APPSwe_NCM) and wild type control (WT_NCM) was collected. The concentration of Aß in the media was determined with an ELISA kit (LifeTechnologies). The concentration of Aß40 and Aß42 in the neuron media was adjusted to 2000 and 200 pM respectively in Neurobasal media with 0.5% B27, to limit the amount of high abundance proteins such as albumin (APPswe_NCM). Equivalent amounts were used to prepare media from control neurons (WT_NCM).

### Culture of primary mouse astrocytes and collection of astrocyte conditioned media

Primary astrocyte cultures were prepared from the cortex of wild type CD1 mice on postnatal days 1–3 as previously described (Schildge et al. 2013). Briefly, brains were harvested and placed in ice-cold HBSS with HEPES, where the meninges were removed, and the cerebral cortices were dissected and mechanically dissociated. Mixed glial cells were cultured in poly-D-lysine coated flasks and grown in high glucose DMEM media with 10% fetal bovine serum, GlutaMAX™ supplement and 100 units/ml penicillin and 100 μg/ml streptomycin at 37 °C in a humidified incubator with 5% CO2. Media was changed after 2 days and then every 5 days. On day 11-14, microglia cells and oligodendrocytes were removed by overnight shaking at 200 rpm and astrocytes were seeded on poly-D-lysine coated 6-multiwell plates and were used after 3 days at 80-90% confluency (>95% of cells were GFAP-positive).

Astrocyte growth medium was washed and changed to Neurobasal media supplemented with 0.5% B27 medium 24h prior to treatment. Astrocytes were cultured in Neurobasal media without B27 for 24h, to collect astrocyte conditioned media and prepare an in-house library. Astrocytes were cultured in APPswe_NCM and WT_NCM for 24h, to collect astrocyte conditioned media to analyse by mass spectrometry (APPse_ACM and WT_ACM). Media were filtered through a 0.22 μm spin filter to eliminate any detached cells or debris and concentrated through a 3 kDa molecular weight cut-off concentrating column at 13,000 g for 30 min, prior to analysis.

### Secretome analysis

Conditioned media from 5 independent neuronal cultures, with each biological replicate analysed in duplicate, were analyzed as previously described (Matafora et al. 2020) by Secret3D workflow (Matafora and Bachi 2020). Collected media was filtered on microcon filters with 10 KDa cutoff (Millipore) and buffer was exchanged with 8 M Urea 100 mM Tris pH8. 50 μg of secreted protein was sonicated with BIORUPTOR (3 cycles: 30 s on/30 s off). Cysteine reduction and alkylation were performed adding 10 mM TCEP (Thermo Scientific) and 40 mM 2-Chloroacetamide (Sigma-Aldrich) in 8 M Urea 100 mM Tris pH 8 for 30 min at room temperature. By using microcon filters with 10 kDa cutoff (Millipore), buffer was exchanged by centrifugation at 9,300 g for 10 min, and PNGase F (New England Biolabs) (1:100=enzyme: secreted proteins) was added for 1 h at room temperature following manufacturer’s instruction. Buffer was again exchanged by centrifugation at 9,300 g for 10 min with 50 mM ammonium bicarbonate, and proteins in solution were digested by Lys-C and trypsin (Kulak et al. 2014). Peptides were recovered on the bottom of the microcon filters by centrifugation at 9,300 g for 10 min, adding two consecutive washes of 50 μl of 0.5 M NaCl. Eluted peptides were purified on a homemade C18 StageTip. 1 μg of digested sample was injected onto a quadrupole Orbitrap Q-exactive HF mass spectrometer (Thermo Scientific). Peptide separation was achieved on a linear gradient from 95% solvent A (2% ACN, 0.1% formic acid) to 55% solvent B (80% acetonitrile, 0.1% formic acid) over 75 min and from 55% to 100% solvent B in 3 min at a constant flow rate of 0.25 μl/min on UHPLC Easy-nLC 1000 (Thermo Scientific) where the LC system was connected to a 23-cm fused-silica emitter of 75 μm inner diameter (New Objective, Inc. Woburn, MA, USA), packed in-house with ReproSil-Pur C18-AQ 1.9 μm beads (Dr Maisch Gmbh, Ammerbuch, Germany) using a high-pressure bomb loader (Proxeon, Odense, Denmark).

The mass spectrometer was operated in DDA (Data Dependent Acquisition) mode: dynamic exclusion enabled (exclusion duration = 15 s), MS1 resolution = 70,000, MS1 automatic gain control target = 3 × 106, MS1 maximum fill time = 60 ms, MS2 resolution = 17,500, MS2 automatic gain control target = 1 × 105, MS2 maximum fill time = 60 ms, and MS2 normalized collision energy = 25. For each cycle, one full MS1 scan range = 300–1,650 m/z was followed by 12 MS2 scans using an isolation window of 2.0 m/z.

The mass spectrometry proteomics data have been deposited to the ProteomeXchange Consortium via the PRIDE (Perez-Riverol et al. 2022) partner repository with the dataset identifier PXD036343.

### MS analysis and database search

MS analysis was performed as reported previously (Geyer et al. 2016). Raw MS files were processed with MaxQuant software (1.6.0.16), making use of the Andromeda search engine (Cox et al. 2011). MS/MS peak lists were searched against the UniProtKB Mouse complete proteome database (uniprot_cp_mouse_2019) in which trypsin specificity was used with up to two missed cleavages allowed. Searches were performed selecting alkylation of cysteine by carbamidomethylation as fixed modification, and oxidation of methionine, N-terminal acetylation and N-Deamination as variable modifications. Mass tolerance was set to 5 ppm and 10 ppm for parent and fragment ions, respectively. A reverse decoy database was generated within Andromeda, and the false discovery rate (FDR) was set to < 0.01 for peptide spectrum matches (PSMs). For identification, at least two peptide identifications per protein were required, of which at least one peptide had to be unique to the protein group. Matching between runs was performed across all samples that are neuron conditioned media and astrocyte conditioned media, plus conditioned media from astrocyte without serum that was used as reference library.

### Quantification and statistical analysis

DDA.raw files were analyzed by MaxQuant software for protein quantitation, and LFQ intensities were used. Statistical analysis was performed by using Perseus software (version 1.5.6.0) included in the MaxQuant package. Outliers were excluded based on the Principal Component Analysis (PCA) of all the samples. T-test statistical analysis was performed applying FDR < 0.05 or *p* < 0.05 as reported.

### Protein localisation prediction

SignalP-6.0 (https://services.healthtech.dtu.dk/service.php?SignalP) (Teufel et al. 2022), TargetP-2.0 (https://services.healthtech.dtu.dk/service.php?TargetP-2.0) (Armenteros et al. 2019), SecretomeP-2.0 (https://services.healthtech.dtu.dk/service.php?SecretomeP-2.0) (Bendtsen et al. 2004), DeepLoc-2.0 (https://services.healthtech.dtu.dk/service.php?DeepLoc-2.0) (Thumuluri et al. 2022), ProtComp 9.0 (http://www.softberry.com/berry.phtml?topic=protcompan&group=programs&subgroup=proloc) (Softberry, Inc., Mount Kisco, NY, USA), and GO annotations from LAGO analysis were used to identify which proteins are predicted to be secreted or found in the extracellular space.

DeepTMHMM (https://dtu.biolib.com/DeepTMHMM) (Hallgren et al. 2022) was used to predict the presence of transmembrane helices, and the Vesiclepedia database (http://microvesicles.org/) (Pathan et al. 2019) was used for the manual search of proteins reported to be secreted via extracellular vesicles.

### Bioinformatics analysis

Gene ontology (GO) term over-representation for biological process, molecular function, and cell component were identified using LAGO (Boyle et al. 2004) (https://go.princeton.edu/). *P*-values were calculated based on the hypergeometric distribution with the Bonferroni correction for multiple comparisons, with *p* < 0.05 after the correction considered significant. STRING 11.5 (https://string-db.org/) (Szklarczyk et al. 2021) was used to predict protein-protein interactions (PPIs) within the differentially expressed proteins, and local STRING network clusters were used to identify if any of the interacting proteins take part in the same biological process. Pathway enrichment analysis was performed using the Reactome pathway database using the online PANTHER v16.0 software (http://pantherdb.org/) (Mi et al. 2021) using Fisher’s Exact test and FDR to correct for multiple comparisons. Statistical significance was defined as FDR < 0.05. Functional Annotation Clustering of differentially upregulated proteins was performed using the Database for Annotation, Visualisation and Integration Discovery (DAVID, (Sherman et al. 2022) (https://david.ncifcrf.gov/home.jsp). Terms from Gene Ontology (GOTERM_BP_DIRECT, GOTERM_CC_DIRECT, and GOTERM_MF_DIRECT) and Pathways (KEGG_PATHWAY, REACTOME_PATHWAY, and WIKIPATHWAYS) were included in clustering analysis. The Enrichment Threshold EASE score was set at 1.3 (*p*-value ≤0.05). Classification stringency was set at high. The whole genome from *Mus musculus* was used as a background. Gene-disease Association (GDA) scores of genes for the differentially expressed proteins were calculated using DisGeNET v7.0 (https://www.disgenet.org/) (Piñero et al. 2021). The genes were filtered for association with the following terms: ‘Disease: Alzheimer’s Disease; CUI: C0002395’ and ‘Disease: Familial Alzheimer Disease (FAD); CUI: C0276496’.

## AUTHOR CONTRIBUTION

VM performed the mass spectrometry analysis and database search; AG did the bioinformatic analysis and comparison with transcriptomic and CSF studies; BGP-N and MJ-S undertook the animal and cell culture work and prepared samples for analysis; VM and AB conceived and designed the proteomics analysis; BGP-N and MJ-S conceived and designed the animal and cell work and the overall project; VM, AG, BGP-N and MJ-S wrote the manuscript which was commented by WN and AB.

## ACKNOWLEDGEMENTS

We would like to thank Cogentech Proteomics/MS Facility (IFOM) for critical comments and suggestions and Elena Galea (Universidad Autonoma de Barcelona) for her critical reading of the manuscript. This work is funded by an ARUK KCL Network Pump Priming grant (BGP-N, MJ-S), the Van Geest Foundation (BGP-N) and Medical Research Council Career Development Award (MR/N022696/1) and Transition Support Award (MR/V036947/1) (MJ-S).

## CONFLICT OF INTEREST

The authors declare no conflict of interest to declare that are relevant to the content of this article.

## DATA AVAILABILITY

The data that support the findings of this study are openly available ProteomeXchange Consortium via the PRIDE (Perez-Riverol et al. 2022) partner repository with the dataset identifier PXD036343.

For the purpose of open access, the author has applied a Creative Commons Attribution (CC BY) licence to any Author Accepted Manuscript version arising.

## Notes

### Competing Interest Statement

The authors have declared no competing interest.

