## Supplemental Figures for "Proteomics of the astrocyte secretome reveals changes in their response to soluble oligomeric Aß"

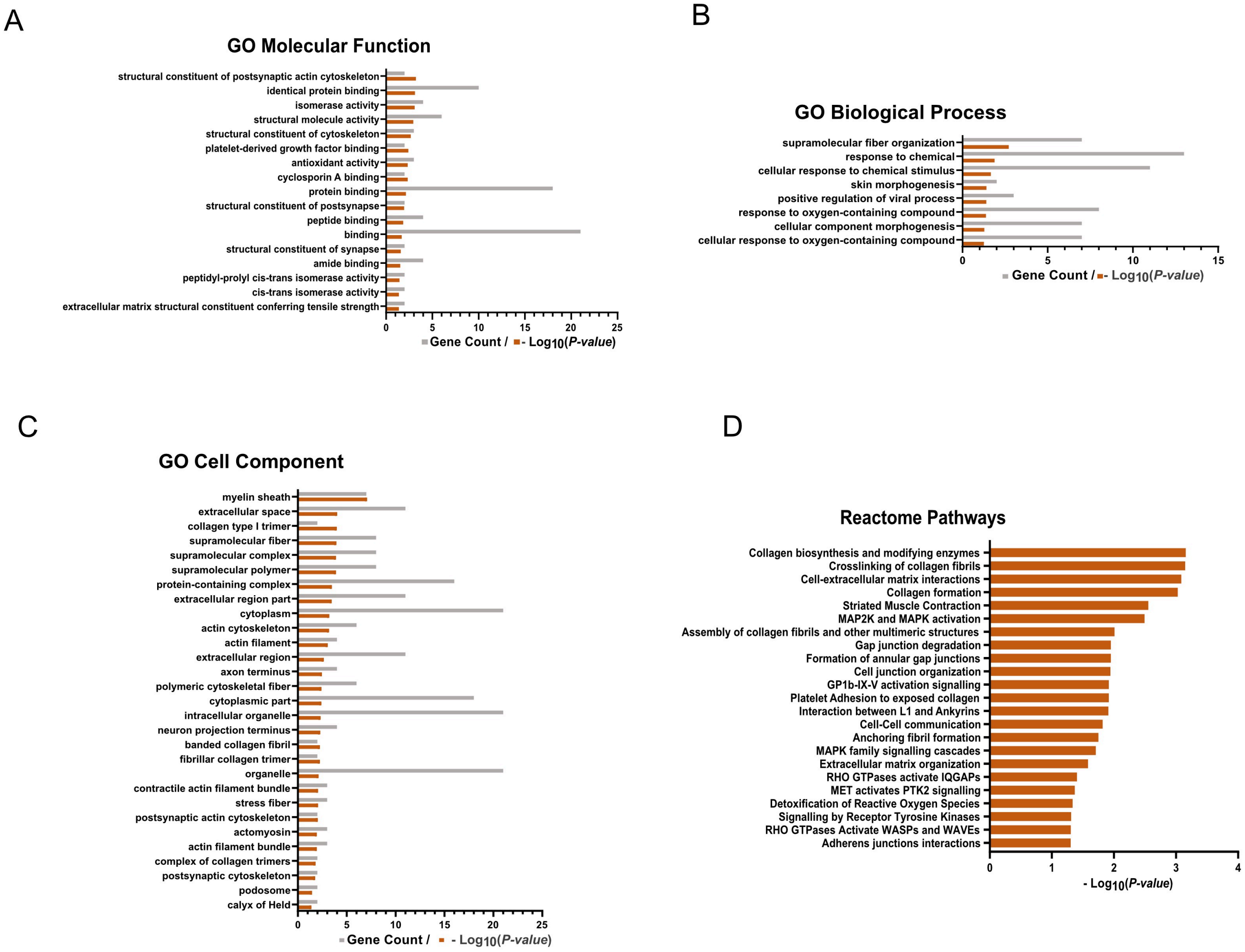


**Supplementary Figure 1**. Gene Ontology (GO) analysis for Molecular Function **(A)**, Biological Process **(B)**, Cell Component **(C)**, or Reactome pathway analysis **(D)** of proteins differentially upregulated in the media of astrocytes treated with media containing Aβ oligomers. The overrepresentation significance threshold was set at the p-value < 0.05.

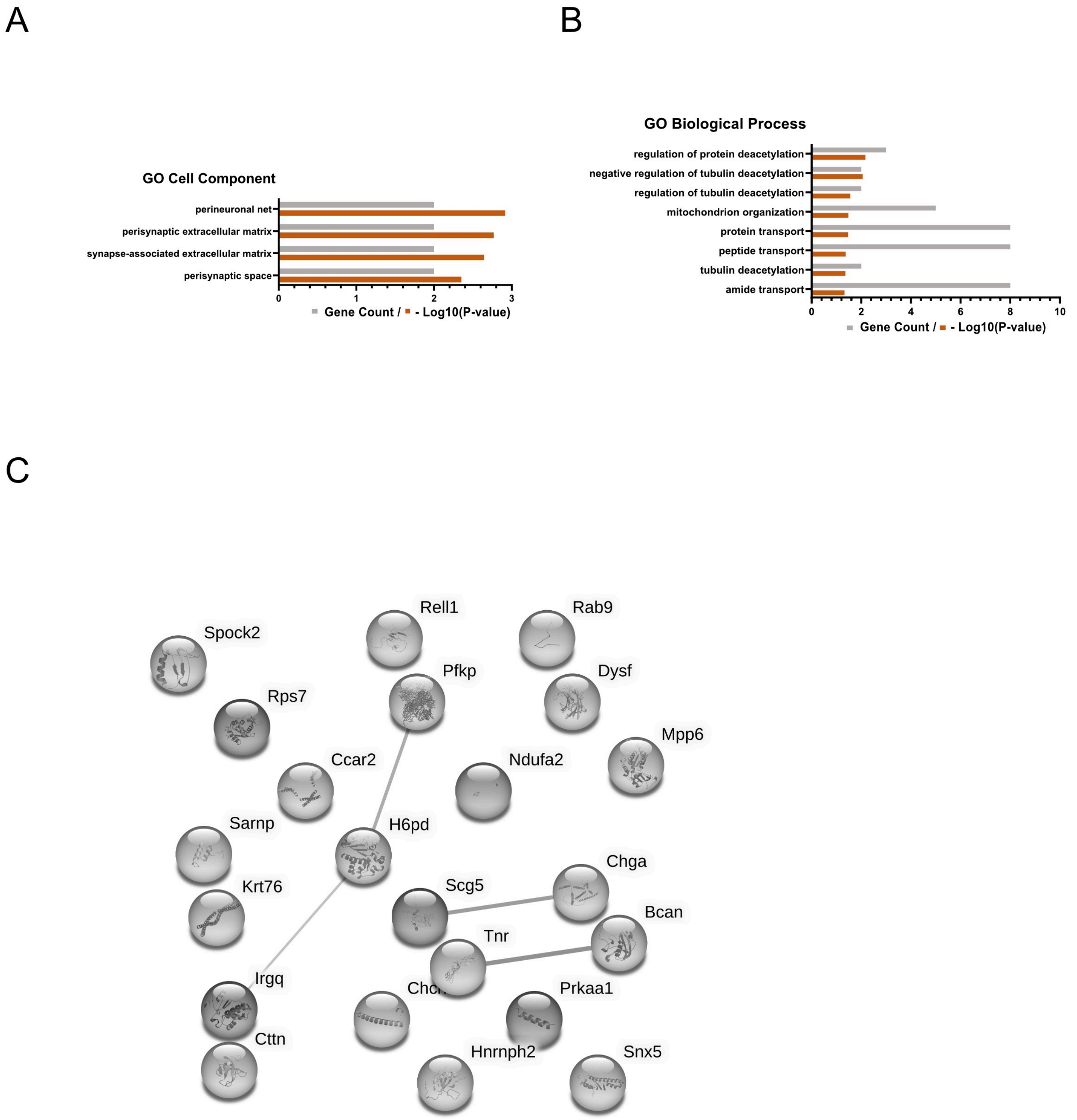


**Supplementary Figure 2**. Gene Ontology (GO) analysis for Cell Component **(A)** and Biological Process **(B)** of proteins differentially upregulated in the media of astrocytes treated with media containing Aß oligomers. The overrepresentation significance threshold was set at the p-value < 0.05. **(C)** Protein-protein interaction network for differentially downregulated proteins generated using STRING. Lines represent protein-protein associations (including but not limited to physical binding) and different thicknesses represent confidence in the interaction. Thicker lines represent higher confidence in the interaction. Circles represent individual proteins.

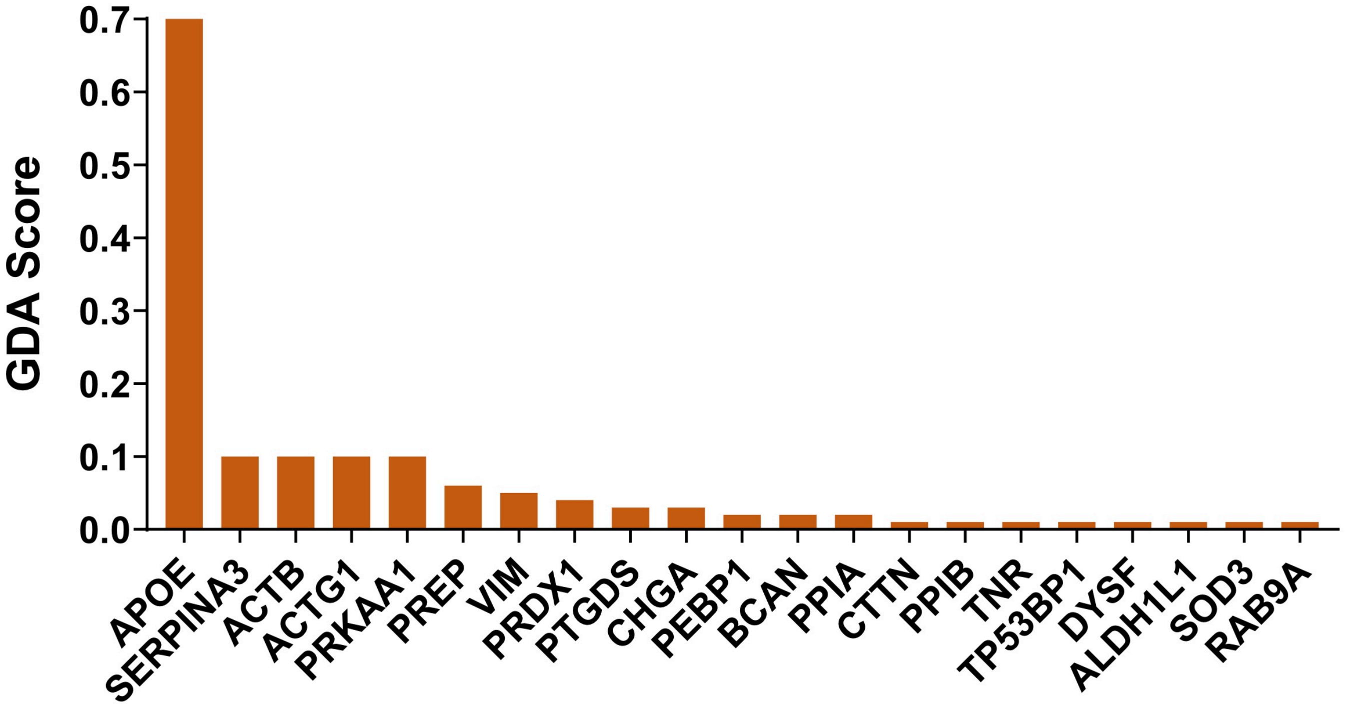


**Supplementary Figure. 3.** Gene-disease association (GDA) scores for the genes coding for proteins identified in this study and found associated with AD using the DisGeNET database. Score shows its association that were found to be associated with Alzheimer’s disease
