## Supplemental Tables for "Proteomics of the astrocyte secretome reveals changes in their response to soluble oligomeric Aß"

**Supplementary Table 1**. List of all identified proteins in the astrocyte secretome upon treatment with APPswe_NCM or WT_NCM, after excluding outliers. In bold, those proteins with WT_ACM vs APPSWE_ACM p value <0.05 who were not found in either APPswe_NCM or WT_NCM. Thick border shows those proteins that were only found in ACM but not in NCM; dotted border highlights those proteins that were found also in NCM and therefore were not analysed further.

| T: Protein IDs | T: Majority protein IDs | T: Protein names | T: Gene names | APPswe_NCM: secretome from APPswe neurons (LFQ intensity) | APPswe_ACM-secretome from astrocytes treated with APPswe_NCM (LFQ intensity) | WT_ACM: secretome from astrocytes treated with WT_NCM (LFQ intensity) | WT_NCM: secretome from WT neurons (LFQ intensity) |
|---|---|---|---|---|---|---|---|
| Q9QUR6 | Q9QUR6 | Prolyl endopeptidase | *Prep* | [illegible] | [illegible] | [illegible] | [illegible] |
| P27773;F6Q404 | P27773 | Protein disulfide-isomerase A3 | *Pdia3* | [illegible] | [illegible] | [illegible] | [illegible] |
| P60710;E9Q1F2;E9Q5F4;G3UZ07;E9Q606;E9Q2D1;F6WX90; | P60710;E9Q1F2;E9Q5F4 | Actin, cytoplasmic 1;Actin, cytoplasmic 1, N-terminally | *Actb* | [illegible] | [illegible] | [illegible] | [illegible] |
| P62911;P17932 | P62911 | 60S ribosomal protein L32 | *Rpl32* | [illegible] | [illegible] | [illegible] | [illegible] |
| O09114-2;O09114 | O09114-2;O09114 | Prostaglandin-H2 D-isomerase | *Ptgds* | [illegible] | [illegible] | [illegible] | [illegible] |
| P70296;D3Z1V4;D6RHS6;E9QLE5;Q8VIN1-2;Q8VIN1 | P70296;D3Z1V4;D6RHS6 | Phosphatidylethanolamine-binding protein 1;Hippoca | *Pebp1* | [illegible] | [illegible] | [illegible] | [illegible] |
| P68033;P68134;A0A1D5RM20 | P68033;P68134 | Actin, alpha cardiac muscle 1;Actin, alpha skeletal mus | *Actc1;Acta1* | [illegible] | [illegible] | [illegible] | [illegible] |
| P08226;A0A1B0GX15;G3UWN5;G3UZM8 | P08226;A0A1B0GX15;G3UWN5 | Apolipoprotein E | *Apoe* | [illegible] | [illegible] | [illegible] | [illegible] |
| Q01149;E0CXI2 | Q01149 | Collagen alpha-2(I) chain | *Col1a2* | [illegible] | [illegible] | [illegible] | [illegible] |
| O09164 | O09164 | Extracellular superoxide dismutase [Cu-Zn] | *Sod3* | [illegible] | [illegible] | [illegible] | [illegible] |
| A0A0G2JE29;D3YUE2;Q61398;D3YWH0;D3YZE3 | A0A0G2JE29;D3YUE2;Q61398;D3YWH0;[ | Procollagen C-endopeptidase enhancer 1 | *Pcolce* | [illegible] | [illegible] | [illegible] | [illegible] |
| P24369 | P24369 | Peptidyl-prolyl cis-trans isomerase B | *Ppib* | [illegible] | [illegible] | [illegible] | [illegible] |
| E9Q1G8;E9Q9F5;O55131;S4R2P6;Q9DA97 | E9Q1G8;E9Q9F5;O55131 | Septin-7 | *Septin 7* | [illegible] | [illegible] | [illegible] | [illegible] |
| Q8R0Y6;A0A0N4SW56;A0A0N4SVT2 | Q8R0Y6 | Cytosolic 10-formyltetrahydrofolate dehydrogenase | *Aldh1l1* | [illegible] | [illegible] | [illegible] | [illegible] |
| P11087;P11087-2;CON__Q862S4 | P11087;P11087-2 | Collagen alpha-1(I) chain | *Col1a1* | [illegible] | [illegible] | [illegible] | [illegible] |
| Q6IRU2 | Q6IRU2 | Tropomyosin alpha-4 chain | *Tpm4* | [illegible] | [illegible] | [illegible] | [illegible] |
| P70399-6;P70399-5;P70399;F6ZRL3 | P70399-6;P70399-5;P70399;F6ZRL3 | Tumor suppressor p53-binding protein 1 | *Tp53bp1;Trp53bp1* | [illegible] | [illegible] | [illegible] | [illegible] |
| Q9EPC1;Q3UF75;D6RHM9;F2Z417 | Q9EPC1;Q3UF75 | Alpha-parvin | *Parva* | [illegible] | [illegible] | [illegible] | [illegible] |
| P35700;B1AXW5;B1AXW6;B1AXW4 | P35700;B1AXW5;B1AXW6;B1AXW4 | Peroxiredoxin-1 | *Prdx1* | [illegible] | [illegible] | [illegible] | [illegible] |
| P20152;A0A0A6YWC8;A2AKJ2;G5E846;A0A2R8W6R6;G3X9 | P20152;A0A0A6YWC8 | Vimentin | *Vim* | [illegible] | [illegible] | [illegible] | [illegible] |
| P17742;A0A1L1SST0;F8VPN3;A0A1L1SRX5;V9GXC1;V9GX3 | P17742;A0A1L1SST0;F8VPN3 | Peptidyl-prolyl cis-trans isomerase A;Peptidyl-prolyl ci | *Ppia;Gm5160* | [illegible] | [illegible] | [illegible] | [illegible] |
| P63260;B1ATY1;G3UYG0 | P63260 | Actin, cytoplasmic 2;Actin, cytoplasmic 2, N-terminally | *Actg1* | [illegible] | [illegible] | [illegible] | [illegible] |

| T: Gene names | C: Student's T-test | C: Student's T-test Significant WT_ACM vs APPSwe_ACM FDR<0.05 | C: Student's T-test Significant WT_ACM vs APPSwe_ACM_ pvalue<0.05 | N: Peptides | N: Sequence coverage [%] | N: Unique sequence coverage [%] | N: Mol. weight [kDa] | N: Score | N: Intensity | N: MS/MS count | N: -Log Student's T-test p-value WT_CM_TG_CM | N: Student's T-test q-value (FDR) WT_CM_TG_CM | LOG2 ratio WT_CM/TG_CM | N: Student's T-test Test statistic WT_CM_TG_CM |
|---|---|---|---|---|---|---|---|---|---|---|---|---|---|---|
| *Prep* | WT_CM_TG_CM | | + | 27 | 49.4 | 49.4 | 80.751 | 250.06 | 12305000000 | 100 | 2.82449 | 0.197 | -1.89925 | -4.00497 |
| *Pdia3* | WT_CM_TG_CM | | + | 52 | 69.3 | 69.3 | 56.678 | 323.31 | 1.2971E+11 | 380 | 1.97523 | 0.2215 | -1.70793 | -2.98262 |
| *Actb* | WT_CM_TG_CM | | + | 49 | 93.3 | 4.5 | 41.736 | 268.94 | 9635900000 | 38 | 2.44771 | 0.159636 | -1.65934 | -3.54846 |
| *Rpl32* | WT_CM_TG_CM | | + | 12 | 59.3 | 59.3 | 15.86 | 137.5 | 15770000000 | 43 | 2.54016 | 0.1538 | -1.62345 | -3.65979 |
| *Ptgds* | WT_CM_TG_CM | | + | 2 | 19 | 19 | 13.941 | 8.4679 | 20372000 | 6 | 1.76299 | 0.256162 | -1.57334 | -2.72766 |
| *Pebp1* | WT_CM_TG_CM | | + | 16 | 84 | 84 | 20.83 | 323.31 | 24935000000 | 106 | 1.98519 | 0.244286 | -1.5329 | -2.99455 |
| *Actc1;Acta1* | WT_CM_TG_CM | | + | 44 | 85.1 | 8.8 | 42.019 | 323.31 | 1.9415E+11 | 342 | 1.70849 | 0.254619 | -1.46468 | -2.66183 |
| *Apoe* | WT_CM_TG_CM | + | + | 45 | 80.4 | 80.4 | 35.866 | 323.31 | 1.4141E+11 | 515 | 4.36141 | 0.016 | -1.45497 | -6.01326 |
| *Col1a2* | WT_CM_TG_CM | | + | 52 | 49.1 | 48.4 | 129.56 | 323.31 | 11749000000 | 331 | 1.44659 | 0.32871 | -1.44104 | -2.34191 |
| *Sod3* | WT_CM_TG_CM | | + | 12 | 55.4 | 55.4 | 27.392 | 323.31 | 5488200000 | 91 | 1.86043 | 0.25 | -1.43871 | -2.84494 |
| *Pcolce* | WT_CM_TG_CM | | + | 3 | 19.9 | 19.9 | 26.424 | 9.3638 | 93988000 | 8 | 1.32236 | 0.354578 | -1.38566 | -2.18719 |
| *Ppib* | WT_CM_TG_CM | | + | 15 | 56.9 | 56.9 | 23.713 | 163.55 | 25631000000 | 114 | 1.46538 | 0.330475 | -1.37326 | -2.36512 |
| *Septin 7* | WT_CM_TG_CM | | + | 36 | 66.6 | 64.5 | 50.648 | 323.31 | 42288000000 | 204 | 1.63597 | 0.2755 | -1.35987 | -2.5739 |

| | | | | | |
|---|---|---|---|---|---|
| Q9D6X6;A0A140LJH8;A0A140LHF1;A0A140LHE7;A0A140LIL0; | Q9D6X6;A0A140LJH8;A0A140LHF1;A0A14 | Serine protease 23 | *Prss23* | 4.15112 | [illegible] |
| Q61239 | Q61239 | Protein farnesyltransferase/geranylgeranyltransferase type | *Fnta* | 0.329368 | [illegible] |
| Q6GQT1 | Q6GQT1 | Alpha-2-macroglobulin-P | *A2mp* | -0.693328 | [illegible] |
| P35283;A2CG35 | P35283;A2CG35 | Ras-related protein Rab-12 | *Rab12* | -0.354142 | [illegible] |
| P24270;A2AL20 | P24270;A2AL20 | Catalase | *Cat* | 0.640325 | [illegible] |
| O88569;O88569-2;O88569-3;A0A0N4SUM2 | O88569;O88569-2;O88569-3 | Heterogeneous nuclear ribonucleoproteins A2/B1 | *Hnrnpa2b1* | -0.628746 | [illegible] |
| Q99LB4;D3YTL5;P24452;D3YZN3;D3YU77;D3Z4K5 | Q99LB4;D3YTL5;P24452 | Macrophage-capping protein | *Capg* | -0.429127 | [illegible] |
| P10711;E9PYD5;P10711-2 | P10711;E9PYD5;P10711-2 | Transcription elongation factor A protein 1 | *Tcea1* | -1.15673 | [illegible] |
| P54818;A0A1Y7VM63 | P54818;A0A1Y7VM63 | Galactocerebrosidase | *Galc* | -0.253889 | [illegible] |
| E9Q3X0;Q9EQK5;D3Z2N7 | E9Q3X0;Q9EQK5 | Major vault protein | *Mvp* | -0.444836 | [illegible] |
| A2A6Q2;Q9JM96;A2A6Q3 | A2A6Q2;Q9JM96;A2A6Q3 | Cdc42 effector protein 4 | *Cdc42ep4* | -0.847655 | [illegible] |
| E9PZ00;Q8BFQ1;K3W4L3;J3QPG5;Q61207 | E9PZ00;Q8BFQ1;K3W4L3;J3QPG5;Q6120 | Prosaposin | *Psap* | -0.899803 | [illegible] |
| P07141;P07141-2;D3Z090;F6RNW8;D3YTW1 | P07141;P07141-2;D3Z090 | Macrophage colony-stimulating factor 1;Processed macrop | *Csf1* | -0.807341 | [illegible] |
| P06797;A0A1Y7VNM3;A0A1Y7VNQ4;A0A1Y7VJC1;A0A1Y7VP4 | P06797;A0A1Y7VNM3;A0A1Y7VNQ4 | Cathepsin L1;Cathepsin L1 heavy chain;Cathepsin L1 light | *Ctsl* | -0.469395 | [illegible] |
| P47867-2;P47867;A0A1L1SV66 | P47867-2;P47867 | Secretogranin-3 | *Scg3* | -0.463684 | [illegible] |
| A8DUK4;P02089;P02088;E9Q223;P02104 | A8DUK4;P02089;P02088;E9Q223 | Hemoglobin subunit beta-2;Hemoglobin subunit beta-1 | *Hbbt1;Hbb-b2;Hbb-b1;Hbb-bs* | -0.235304 | [illegible] |
| P03921 | P03921 | NADH-ubiquinone oxidoreductase chain 5 | *Mtnd5* | -0.336968 | [illegible] |
| Q9Z2T6 | Q9Z2T6 | Keratin, type II cuticular Hb5 | *Krt85* | 0.177827 | [illegible] |
| P01902;F6TIX5;Q9BCZ1;A7VMS2;Q31615;F6T1I5;Q05A75;P06 | P01902 | H-2 class I histocompatibility antigen, K-D alpha chain | *H2-K1* | -0.511894 | [illegible] |
| E9Q9C3;Q9QZQ1-2;Q9QZQ1;E9Q852;E9PYX7;D3YUD2;F7C3 | E9Q9C3;Q9QZQ1-2;Q9QZQ1;E9Q852;E9I | Afadin | *Mllt4* | -0.848173 | [illegible] |
| P01837 | P01837 | Ig kappa chain C region | | -0.315289 | [illegible] |
| A0A0R4J0S1;Q91W92 | A0A0R4J0S1;Q91W92 | Cdc42 effector protein 1 | *Cdc42ep1* | -0.368409 | [illegible] |
| P08905;P17897 | P08905;P17897 | Lysozyme C-2;Lysozyme C-1 | *Lyz2;Lyz1* | -0.588043 | [illegible] |
| Q8C788;Q91ZR2 | Q8C788;Q91ZR2 | Sorting nexin;Sorting nexin-18 | *Snx18* | -0.414551 | [illegible] |
| A2AAY5 | A2AAY5 | SH3 and PX domain-containing protein 2B | *Sh3pxd2b* | -0.807045 | [illegible] |
| K7N6V6;J3QNK3;J3QM74;M0QWX9;L7N298;K7N6T8;E9Q058; | K7N6V6;J3QNK3;J3QM74;M0QWX9;L7N29 | [illegible] | [illegible];Gm3558;Gm2974;Gm10340;Gm3667;Gm3 | -0.212202 | [illegible] |
| Q9CQF9;F7CIP8;A0A0N4SWD3;D3Z275 | Q9CQF9 | Prenylcysteine oxidase | *Pcyox1* | -0.394285 | [illegible] |
| Q9Z1Y4 | Q9Z1Y4 | Thyroid receptor-interacting protein 6 | *Trip6* | -0.364589 | [illegible] |
| P14733;A0A0R4J0Q5;P21619;P21619-2 | P14733 | Lamin-B1 | *Lmnb1* | -0.559587 | [illegible] |
| G3X9V2;P30999;E9Q8Z6;E9Q8Z8;E9Q8Z4;E9Q8Z9;D3Z2H2;D | G3X9V2;P30999;E9Q8Z6;E9Q8Z8;E9Q8Z4 | Catenin delta-1 | *Ctnnd1* | -0.234545 | [illegible] |
| Q8BH95 | Q8BH95 | Enoyl-CoA hydratase, mitochondrial | *Echs1* | -0.176777 | [illegible] |
| Q80YX1;Q80YX1-2;Q80YX1-3;Q80YX1-4;Q80YX1-5;Q80YX0 | Q80YX1;Q80YX1-2;Q80YX1-3;Q80YX1-4;C | Tenascin | *Tnc* | -0.16387 | [illegible] |
| P10148 | P10148 | C-C motif chemokine 2 | *Ccl2* | -0.472234 | [illegible] |
| P70232;P70232-2;A0A0N4SUM4;A0A0N4SUL1 | P70232;P70232-2 | Neural cell adhesion molecule L1-like protein;Processed ne | *Chl1* | -1.35848 | [illegible] |
| Q9D0J8 | Q9D0J8 | Parathymosin | *Ptms* | -0.193774 | [illegible] |
| Q922U1;E0CYD0 | Q922U1 | U4/U6 small nuclear ribonucleoprotein Prp3 | *Prpf3* | -0.393183 | [illegible] |
| Q8K2B3;A0A1Y7VJ55 | Q8K2B3 | Succinate dehydrogenase [ubiquinone] flavoprotein subunit | *Sdha* | -0.892338 | [illegible] |
| Q99LS0 | Q99LS0 | Augurin | *Ecrg4* | -0.860554 | [illegible] |
| Q921I6 | Q921I6 | SH3 domain-binding protein 4 | *Sh3bp4* | 0.725128 | [illegible] |
| Q9EQC5 | Q9EQC5 | N-terminal kinase-like protein | *Scyl1* | 0.972428 | [illegible] |
| P26645 | P26645 | Myristoylated alanine-rich C-kinase substrate | *Marcks* | -1.25987 | [illegible] |
| E9PZS8;Q61387 | E9PZS8;Q61387 | Cytochrome c oxidase subunit 7A-related protein, mitochor | *Cox7a2l* | -0.540209 | [illegible] |
| Q9R0N0 | Q9R0N0 | Galactokinase | *Galk1* | -0.469547 | [illegible] |
| I7HLV2;Q6ZWV3;P86048;A0A1B0GXC3 | I7HLV2;Q6ZWV3;P86048 | 60S ribosomal protein L10;60S ribosomal protein L10-like | *Rpl10;Rpl10l* | -0.612824 | [illegible] |
| Q9EPL2 | Q9EPL2 | Calsyntenin-1;Soluble Alc-alpha;CTF1-alpha | *Clstn1* | -0.252818 | [illegible] |
| A2ARP8;Q9QYR6;Q9QYR6-2 | A2ARP8;Q9QYR6;Q9QYR6-2 | Microtubule-associated protein 1A;MAP1A heavy chain;MA | *Map1a* | -1.06573 | [illegible] |
| Q9CY97;Q9CY97-2 | Q9CY97;Q9CY97-2 | RNA polymerase II subunit A C-terminal domain phosphata | *Ssu72* | -0.176777 | [illegible] |
| P61982 | P61982 | 14-3-3 protein gamma;14-3-3 protein gamma, N-terminally | *Ywhag* | -1.06262 | [illegible] |
| P70268;P70268-2;D6RH37 | P70268;P70268-2;D6RH37 | Serine/threonine-protein kinase N1 | *Pkn1* | 1.03755 | [illegible] |
| Q64523;Q6GSS7;Q64522 | Q64523;Q6GSS7 | Histone H2A type 2-C;Histone H2A type 2-A | *Hist2h2ac;Hist2h2aa1* | 2.3156 | [illegible] |
| Q6A0A9 | Q6A0A9 | Constitutive coactivator of PPAR-gamma-like protein 1 | *FAM120A* | -0.20286 | [illegible] |
| D3YWC9;Q6P9S0;F7D291;F6S054 | D3YWC9;Q6P9S0 | MTSS1-like protein | *Mtss1l* | -0.25087 | [illegible] |
| Q08761 | Q08761 | Vitamin K-dependent protein S | *Pros1* | -0.982185 | [illegible] |
| P26350;A0A087WP98;A0A087WPN6;A0A087WQN2 | P26350 | Prothymosin alpha;Prothymosin alpha, N-terminally proces | *Ptma* | 0.237832 | [illegible] |
| Q6R891;Q6R891-2 | Q6R891 | Neurabin-2 | *Ppp1r9b* | 1.25341 | [illegible] |
| P30681;A0A1B0GQX9 | P30681;A0A1B0GQX9 | High mobility group protein B2 | *Hmgb2* | 1.82205 | [illegible] |
| P43406;A2AKI5;B7ZCI9;F6RWM8;F6W6Q6 | P43406;A2AKI5 | Integrin alpha-V;Integrin alpha-V heavy chain;Integrin alpha | *Itgav* | 0.867003 | [illegible] |
| P14131 | P14131 | 40S ribosomal protein S16 | *Rps16* | -0.220135 | [illegible] |
| Q9R0L6;Q9R0L6-2 | Q9R0L6;Q9R0L6-2 | Pericentriolar material 1 protein | *Pcm1* | -0.619732 | [illegible] |
| A0A0R4IZZ1;Q03157 | A0A0R4IZZ1;Q03157 | Amyloid-like protein 1;C30 | *Aplp1* | -1.7402 | [illegible] |
| Q62426 | Q62426 | Cystatin-B | *Cstb* | -0.271808 | [illegible] |
| Q91XV3 | Q91XV3 | Brain acid soluble protein 1 | *Basp1* | -0.243586 | [illegible] |
| B7ZP47;Q65Z40;F6YCH1 | B7ZP47;Q65Z40;F6YCH1 | Wings apart-like protein homolog | *Wapal* | 1.11684 | [illegible] |
| G3X8Y3;A0A0A6YW80;Q80UM3;A0A0A6YX86;A0A0A6YXF4;Q | G3X8Y3;A0A0A6YW80;Q80UM3 | N-alpha-acetyltransferase 15, NatA auxiliary subunit | *Naa15* | -0.659254 | [illegible] |
| P61255;B1ARA5;B1ARA3 | P61255;B1ARA5;B1ARA3 | 60S ribosomal protein L26 | *Rpl26* | 0.339529 | [illegible] |
| E9Q6L0;J3QP81;A0A087WS18;A0A087WQF1;A0A087WQ95;A( | E9Q6L0;J3QP81;A0A087WS18;A0A087WQ | CLIP-associating protein 1 | *Clasp1* | -0.536937 | [illegible] |
| F8WHM5;Q61543;F6RSH1;E9PWD8 | F8WHM5;Q61543 | Golgi apparatus protein 1 | *Glg1* | -0.796087 | [illegible] |
| P14069 | P14069 | Protein S100-A6 | *S100a6* | -0.488684 | [illegible] |
| Q922R8;Q3TML0 | Q922R8;Q3TML0 | Protein disulfide-isomerase A6 | *Pdia6* | -0.51491 | [illegible] |
| P37804;A0A1L1STN8 | P37804 | Transgelin | *Tagln* | -0.306847 | [illegible] |
| A0A0U1RNT6;A0A0U1RNK6;Q3THS6;A0A0U1RQ95;A0A0U1R( | A0A0U1RNT6;A0A0U1RNK6;Q3THS6 | S-adenosylmethionine synthase isoform type-2 | *Mat2a* | -0.388265 | [illegible] |
| Q5SVR0-2;Q5SVR0 | Q5SVR0-2;Q5SVR0 | TBC1 domain family member 9B | *Tbc1d9b* | -0.176777 | [illegible] |
| P19096;A0A0U1RNJ1;A0A0U1RPP5 | P19096;A0A0U1RNJ1 | Fatty acid synthase;[Acyl-carrier-protein] S-acetyltransfera | *Fasn* | -0.636443 | [illegible] |
| Q61686;D3Z0U2 | Q61686 | Chromobox protein homolog 5 | *Cbx5* | 2.63416 | [illegible] |
| A0A286YDF5;Q69ZN7;E9Q390;Q69ZN7-4;A0A286YCZ3;Q69Z | A0A286YDF5;Q69ZN7;E9Q390;Q69ZN7-4; | Myoferlin | *Myof* | 5.12473 | [illegible] |
| Q8C2E7;Q8C2E7-2 | Q8C2E7;Q8C2E7-2 | WASH complex subunit strumpellin | *Kiaa0196* | -0.176777 | [illegible] |
| P80318;E9Q133;Q3U0I3;F6Q609;F6ZVG8 | P80318;E9Q133;Q3U0I3;F6Q609 | T-complex protein 1 subunit gamma | *Cct3* | 0.754361 | [illegible] |
| Q61035 | Q61035 | Histidine--tRNA ligase, cytoplasmic | *Hars* | -0.492767 | [illegible] |
| Q3UHL6;A0A087WR50;P11276;A0A087WS56;B9EHT6;A0A087 | Q3UHL6;A0A087WR50;P11276;A0A087WS | Fibronectin;Anastellin | *Fn1* | 1.19509 | [illegible] |
| Q9R1P4;A0A1B0GS70 | Q9R1P4;A0A1B0GS70 | Proteasome subunit alpha type-1 | *Psma1* | 0.242471 | [illegible] |
| Q6P1B9;O08539-2;O08539 | Q6P1B9;O08539-2;O08539 | Myc box-dependent-interacting protein 1 | *Bin1* | -0.65574 | [illegible] |
| E9Q397;Q3UGX2;P15508;D3YWH8 | E9Q397;Q3UGX2;P15508 | Spectrin beta chain, erythrocytic | *Sptb* | 0.644284 | [illegible] |
| A0A2R8W6P6;A0A2R8VI39;P23591;A0A2R8VHD0;A0A2R8VKL | A0A2R8W6P6;A0A2R8VI39;P23591;A0A2R | GDP-L-fucose synthase | *Tsta3* | -0.415391 | [illegible] |
| P17426-2;P17426;A0A140LIG7;A0A140LHA6 | P17426-2;P17426 | AP-2 complex subunit alpha-1 | *Ap2a1* | 0.882407 | [illegible] |
| P30355;A0A0J9YUC8 | P30355;A0A0J9YUC8 | Arachidonate 5-lipoxygenase-activating protein | *Alox5ap* | -0.417407 | [illegible] |
| Q9CZX8;D3YUT3;D3Z5R8;D3Z722;D3YUG3;S4R223 | Q9CZX8;D3YUT3;D3Z5R8;D3Z722;D3YUG | 40S ribosomal protein S19 | *Rps19* | -0.454503 | [illegible] |
| A0A0R4J233;A0A1W2P6J7;Q8C650-2;Q8C650;A0A1W2P866 | A0A0R4J233;A0A1W2P6J7;Q8C650-2;Q8C | Septin-10 | *Sep-10* | -0.558303 | [illegible] |
| P16460 | P16460 | Argininosuccinate synthase | *Ass1* | -0.372761 | [illegible] |
| Q8C4G3;Q8C288;A2AQC3;O70228 | Q8C4G3;Q8C288;A2AQC3;O70228 | Phospholipid-transporting ATPase;Probable phospholipid-tr | *Atp9a* | 0.201477 | [illegible] |
| D3YTP8;Q9CY46;Q9QXA5;A0A1B0GRR9;A0A1B0GQZ9 | D3YTP8;Q9CY46;Q9QXA5;A0A1B0GRR9 | U6 snRNA-associated Sm-like protein LSm4 | *Lsm4* | -0.452647 | [illegible] |
| Q9D819 | Q9D819 | Inorganic pyrophosphatase | *Ppa1* | 2.61514 | [illegible] |
| Q9ER67;A2AG47;A2AG46 | Q9ER67 | | *Maged2* | -0.176777 | [illegible] |
| E9PWK1;Q9D379;F6YTS6;D3Z4M3 | E9PWK1;Q9D379 | Epoxide hydrolase 1 | *Ephx1* | -0.437148 | [illegible] |
| Q3U962 | Q3U962 | Collagen alpha-2(V) chain | *Col5a2* | -0.644686 | [illegible] |
| Q62000 | Q62000 | Mimecan | *Ogn* | -0.251083 | [illegible] |
| Q5SWU9;Q5SWU9-2;E9Q4Z2;V9GWS0;D3Z0B3;V9GXY1;A0A( | Q5SWU9;Q5SWU9-2 | Acetyl-CoA carboxylase 1;Biotin carboxylase | *Acaca* | -0.311442 | [illegible] |
| Q9D1R9;A0A0G2JEY6;A0A0G2JGY8 | Q9D1R9;A0A0G2JEY6;A0A0G2JGY8 | 60S ribosomal protein L34 | *Rpl34* | -0.634954 | [illegible] |
| D3YWD3;B1AZA5 | D3YWD3;B1AZA5 | Transmembrane protein 245 | *Tmem245* | -0.253294 | [illegible] |
| Q3U8S1;A2APM5;A2APM3;A2APM4;E9QKM8;Q80X37;A2APM1 | Q3U8S1;A2APM5;A2APM3;A2APM4;E9QKI | CD44 antigen | *Cd44* | -0.237224 | [illegible] |
| A0A0U1RNK7;E9PX48;A2A9M5;A2A9M4;Q8R1A4-2;Q8R1A4;A( | A0A0U1RNK7;E9PX48;A2A9M5;A2A9M4;Q | Dedicator of cytokinesis protein 7 | *Dock7* | -0.412054 | [illegible] |
| Q9QYI3;F7BTP8;A2A5E1 | Q9QYI3;F7BTP8 | DnaJ homolog subfamily C member 7 | *Dnajc7* | 0.622349 | [illegible] |
| F8WI30;Q9CY18;F6RSR4;F7D3B8 | F8WI30;Q9CY18;F6RSR4;F7D3B8 | Sorting nexin-7 | *Snx7* | -0.692025 | [illegible] |
| O70572;D6RGM4 | O70572 | Sphingomyelin phosphodiesterase 2 | *Smpd2* | -1.06546 | [illegible] |
| Q8CBB6;Q8CGP2;Q8CGP1;Q64475;P10853;Q8CGP2-2;Q9D2 | Q8CBB6;Q8CGP2;Q8CGP1;Q64475;P108 | Histone H2B;Histone H2B type 1-P;Histone H2B type 1-K;H | *1h2bk;Hist1h2bb;Hist1h2bf;Hist3h2ba;Hist3h* | 1.35141 | [illegible] |
| P20357;Q80X35;Q80ZL4;E9Q8Y2;G3UZE9;G3UZK9 | P20357;Q80X35;Q80ZL4 | Microtubule-associated protein 2;Microtubule-associated pr | *Map2* | -0.379246 | [illegible] |
| Q8JZQ9;A0A0G2JG48 | Q8JZQ9 | Eukaryotic translation initiation factor 3 subunit B | *Eif3b* | 0.104045 | [illegible] |
| D3Z2Z1;F8WIA1;Q922J3;D3Z3M7;Q922J3-2;F6RCU2;Q8C0S5 | D3Z2Z1;F8WIA1;Q922J3;D3Z3M7;Q922J3- | CAP-Gly domain-containing linker protein 1 | *Clip1* | -0.412468 | [illegible] |
| Q9Z0J0 | Q9Z0J0 | Epididymal secretory protein E1 | *Npc2* | -0.448329 | [illegible] |
| Q8BJD1 | Q8BJD1 | Inter-alpha-trypsin inhibitor heavy chain H5 | *Itih5* | -0.306099 | [illegible] |
| Q9D0J4 | Q9D0J4 | ADP-ribosylation factor-like protein 2 | *Arl2* | -0.435075 | [illegible] |
| Q9QXK7;A0A1Y7VLY3;A0A1Y7VJU9;A0A1Y7VJ97 | Q9QXK7;A0A1Y7VLY3 | Cleavage and polyadenylation specificity factor subunit 3 | *Cpsf3* | -0.25014 | [illegible] |
| Q8BQ46;F6QCI0 | Q8BQ46;F6QCI0 | | *Taf15* | -0.313485 | [illegible] |
| Q9D964 | Q9D964 | Glycine amidinotransferase, mitochondrial | *Gatm* | -0.176777 | [illegible] |
| P63017;Q504P4;D3Z5E2 | P63017;Q504P4 | Heat shock cognate 71 kDa protein | *Hspa8* | -1.05473 | [illegible] |
| P52760 | P52760 | Ribonuclease UK114 | *Hrsp12* | -0.248806 | [illegible] |
| Q91XX1;A0A0A6YWM6;A0A0A6YXZ9;A0A0A6YVS5;O88689-3;( | Q91XX1 | | *Pcdhgc3* | -0.78407 | [illegible] |
| P12032 | P12032 | Metalloproteinase inhibitor 1 | *Timp1* | -0.273198 | [illegible] |
| Q8BL66;A0A1W2P7A6 | Q8BL66 | Early endosome antigen 1 | *Eea1* | -0.520103 | [illegible] |
| Q62093 | Q62093 | Serine/arginine-rich splicing factor 2 | *Srsf2* | -0.297981 | [illegible] |
| Q63918 | Q63918 | Serum deprivation-response protein | *Sdpr* | -0.176777 | [illegible] |
| Q9D0E1-2;Q9D0E1;B8JK33;B8JK32;F6W322;F7C9U3 | Q9D0E1-2;Q9D0E1;B8JK33;B8JK32 | Heterogeneous nuclear ribonucleoprotein M | *Hnrnpm* | -0.597218 | [illegible] |
| E9Q512;H3BJG4;Q8BVJ9;H3BJH8;H3BJU5 | E9Q512;H3BJG4 | | *Trip11* | -0.176777 | [illegible] |
| E9QB01 | E9QB01 | | *Ncam1* | -1.09995 | [illegible] |
| Q3TFD2-2;Q3TFD2;Q3TFD2-3 | Q3TFD2-2;Q3TFD2;Q3TFD2-3 | Lysophosphatidylcholine acyltransferase 1 | *Lpcat1* | -0.449465 | [illegible] |
| A6BLY7 | A6BLY7 | Keratin, type I cytoskeletal 28 | *Krt28* | -0.455417 | [illegible] |
| B1ARU4;E9PVY8;E9QA63;Q9QXZ0;A0A0A0MQA6;Q9QXZ0-3;( | B1ARU4;E9PVY8;E9QA63;Q9QXZ0;A0A0A | Microtubule-actin cross-linking factor 1 | *Macf1* | -0.507512 | [illegible] |
| A0A0R4J1R7;Q9CZL5 | A0A0R4J1R7;Q9CZL5 | Pterin-4-alpha-carbinolamine dehydratase 2 | *Pcbd2* | -0.254033 | [illegible] |
| P48774;E9PVM7;E9PV63;E9Q024;E9Q5L9 | P48774;E9PVM7;E9PV63 | Glutathione S-transferase Mu 5 | *Gstm5* | -0.700602 | [illegible] |
| D3Z0V2;A0A0R4J0X8;Q9ES28-8;Q9ES28-4;Q9ES28-3;Q9ES2 | D3Z0V2;A0A0R4J0X8;Q9ES28-8;Q9ES28- | Rho guanine nucleotide exchange factor 7 | *Arhgef7* | -0.411285 | [illegible] |
| B9EJ54;A0A0J9YUD5;E9Q880 | B9EJ54;A0A0J9YUD5 | | *Nup205* | -0.176777 | [illegible] |
| Q8CAY6;G3XA25;A0A3B2WAL8;Q80X81 | Q8CAY6;G3XA25;A0A3B2WAL8;Q80X81 | Acetyl-CoA acetyltransferase, cytosolic | *Acat2;Acat3* | -0.176777 | [illegible] |
| Q9Z2M7 | Q9Z2M7 | Phosphomannomutase 2 | *Pmm2* | -0.176777 | [illegible] |
| Q922Q8 | Q922Q8 | Leucine-rich repeat-containing protein 59 | *Lrrc59* | -0.449898 | [illegible] |
| B1B0C7;E9PZ16;Q05793 | B1B0C7;E9PZ16;Q05793 | Basement membrane-specific heparan sulfate proteoglycar | *Hspg2* | -0.486057 | [illegible] |
| Q3TL44;A0A1L1ST78;A0A1L1SQN7 | Q3TL44;A0A1L1ST78;A0A1L1SQN7 | NLR family member X1 | *Nlrx1* | -0.37049 | [illegible] |
| Q01853 | Q01853 | Transitional endoplasmic reticulum ATPase | *Vcp* | -0.131678 | [illegible] |
| Q9DCD0 | Q9DCD0 | 6-phosphogluconate dehydrogenase, decarboxylating | *Pgd* | -0.399272 | [illegible] |
| P51859;E0CXA0;E0CYW7 | P51859;E0CXA0;E0CYW7 | Hepatoma-derived growth factor | *Hdgf* | 0.351113 | [illegible] |
| P10605 | P10605 | Cathepsin B;Cathepsin B light chain;Cathepsin B heavy cha | *Ctsb* | -0.611716 | [illegible] |
| O35684;E0CYW0 | O35684;E0CYW0 | Neuroserpin | *Serpini1* | -0.375132 | [illegible] |
| Q5F2E7;Q5F2E7-2 | Q5F2E7;Q5F2E7-2 | Nuclear fragile X mental retardation-interacting protein 2 | *Nufip2* | -0.251886 | [illegible] |
| A2AVJ7;Q99PL5;Q99PL5-11;Q99PL5-10;Q99PL5-9;Q99PL5-8;( | A2AVJ7;Q99PL5 | Ribosome-binding protein 1 | *Rrbp1* | -0.140262 | [illegible] |
| Q61805;A2AC65;F6XKX9 | Q61805 | Lipopolysaccharide-binding protein | *Lbp* | -0.682266 | [illegible] |
| Q91VH6;A0A3F2YNL8;A0A3B2W7C9;A0A3B2W4I2;A0A3B2WC( | Q91VH6;A0A3F2YNL8;A0A3B2W7C9;A0A3 | Protein MEMO1 | *Memo1* | -0.176777 | [illegible] |
| Q62048;Q62048-2;D3Z375 | Q62048;Q62048-2;D3Z375 | Astrocytic phosphoprotein PEA-15 | *Pea15;Pea15a* | -0.423207 | [illegible] |
| P61963 | P61963 | DDB1- and CUL4-associated factor 7 | *Dcaf7* | 1.03537 | [illegible] |
| Q8BPG6;A0A0J9YUK9;A0A0J9YUF6 | Q8BPG6 | Sulfatase-modifying factor 2 | *Sumf2* | -0.376795 | [illegible] |
| E9Q1Y3;E9Q414 | E9Q1Y3;E9Q414 | Apolipoprotein B-100;Apolipoprotein B-48 | *Apob* | -1.05457 | [illegible] |
| A0A087WNY2;A0A087WRT0;D3YZP6;A0A0R4J065;Q8BGE6 | A0A087WNY2;A0A087WRT0;D3YZP6;A0A0 | Cysteine protease ATG4B | *Atg4b* | 2.25122 | [illegible] |
| Q71M36-2;Q71M36;Q71M36-4 | Q71M36-2;Q71M36;Q71M36-4 | Chondroitin sulfate proteoglycan 5 | *Cspg5* | -0.421025 | [illegible] |
| P20029 | P20029 | 78 kDa glucose-regulated protein | *Hspa5* | -0.816136 | [illegible] |
| A0A1B0GR11;Q93092 | A0A1B0GR11;Q93092 | Transaldolase | *Taldo1* | 0.0385968 | [illegible] |
| Q8R1G6;A0A087WPL1;F7C957;E9Q996;REV__A2AP83;REV__ | Q8R1G6;A0A087WPL1;F7C957 | PDZ and LIM domain protein 2 | *Pdlim2* | -0.339143 | [illegible] |
| Q99NH8;Q99NH8-2 | Q99NH8;Q99NH8-2 | Triggering receptor expressed on myeloid cells 2 | *Trem2* | 2.87736 | [illegible] |
| Q9D8N0 | Q9D8N0 | Elongation factor 1-gamma | *Eef1g* | -0.374229 | [illegible] |
| G3UZI2;A0A0R4J259;Q7TMK9-2;G3V018;Q7TMK9;G3UZ48;G | G3UZI2;A0A0R4J259;Q7TMK9-2;G3V018;( | Heterogeneous nuclear ribonucleoprotein Q | *Syncrip* | -0.227605 | [illegible] |
| Q3TLP8;P63001;P60764;A2AC13 | Q3TLP8;P63001 | Ras-related C3 botulinum toxin substrate 1 | *Rac1* | -0.176777 | [illegible] |
| G3X9T8;G3X8Q5 | G3X9T8;G3X8Q5 | | *Cp* | -0.474526 | [illegible] |
| Q8K3X4 | Q8K3X4 | Interferon regulatory factor 2-binding protein-like | *Irf2bpl* | -0.409559 | [illegible] |
| Q8C522;A0A1L1SSA0 | Q8C522 | Endonuclease domain-containing 1 protein | *Endod1* | -0.650707 | [illegible] |
| A0A087WR57;Q91XC8;A0A087WRC6 | A0A087WR57;Q91XC8;A0A087WRC6 | Death-associated protein 1 | *Dap* | -0.573156 | [illegible] |
| Q00PI9 | Q00PI9 | Heterogeneous nuclear ribonucleoprotein U-like protein 2 | *Hnrnpul2* | -0.379051 | [illegible] |
| Q64727;A0A286YDJ4 | Q64727 | Vinculin | *Vcl* | 0.694067 | [illegible] |
| P42932;H3BL49;H3BJB6;H3BLL1;H3BKG2;H3BKR8 | P42932;H3BL49 | T-complex protein 1 subunit theta | *Cct8* | -0.508962 | [illegible] |
| Q61477 | Q61477 | Neuroblastoma suppressor of tumorigenicity 1 | *Nbl1* | -0.358804 | [illegible] |
| Q5EBP8;P49312;P49312-2;V9GXF8 | Q5EBP8;P49312;P49312-2 | Heterogeneous nuclear ribonucleoprotein A1;Heterogeneou | *Hnrnpa1* | -0.682065 | [illegible] |
| Q80X85 | Q80X85 | 28S ribosomal protein S7, mitochondrial | *Mrps7* | -0.293777 | [illegible] |
| Q6URW6-2;Q6URW6-3;K3W4R2;Q6URW6;A0A140LI60;A0A14( | Q6URW6-2;Q6URW6-3;K3W4R2;Q6URW6 | Myosin-14 | *Myh14* | 0.453464 | [illegible] |
| P17751;H7BXC3 | P17751;H7BXC3 | Triosephosphate isomerase | *Tpi1* | 2.52968 | [illegible] |
| Q3UPN1;P29533;P29533-2;A0A0G2JFP9 | Q3UPN1;P29533 | Vascular cell adhesion protein 1 | *Vcam1* | -0.538816 | [illegible] |
| Q99LD8;O08972;G3UZR0 | Q99LD8;O08972;G3UZR0 | N(G),N(G)-dimethylarginine dimethylaminohydrolase 2 | *Ddah2* | -0.441258 | [illegible] |
| A0A1W2P7Y9;Q810U4-2;Q810U4;Q810U4-3;Q810U4-4;Q810U | A0A1W2P7Y9;Q810U4-2;Q810U4;Q810U4- | Neuronal cell adhesion molecule | *Nrcam* | -0.469358 | [illegible] |
| Q07797;E9Q5X5 | Q07797 | Galectin-3-binding protein | *Lgals3bp* | 1.72286 | [illegible] |
| Q9CWK8 | Q9CWK8 | Sorting nexin-2 | *Snx2* | -0.573565 | [illegible] |
| P52480;A0A1L1SU37;A0A1L1SQV8;A0A1L1SSN6;A0A1L1STV8 | P52480 | Pyruvate kinase PKM | *Pkm* | -0.268065 | [illegible] |
| E9QPI5;Q6A026;A0A0J9YV33 | E9QPI5;Q6A026 | Sister chromatid cohesion protein PDS5 homolog A | *Pds5a* | -0.176777 | [illegible] |
| Q7TSK2-2;Q7TSK2;F6XNR8;Q7TSK2-3;F6SXT0 | Q7TSK2-2;Q7TSK2;F6XNR8;Q7TSK2-3 | Seizure protein 6 | *Sez6* | -1.50036 | [illegible] |
| P14602-2;P14602;P14602-3;D3YZ06 | P14602-2;P14602;P14602-3;D3YZ06 | Heat shock protein beta-1 | *Hspb1* | -0.336979 | [illegible] |
| Q8K310;A0A087WSU2;A0A087WSP7;A0A087WQZ1;A0A087W( | Q8K310;A0A087WSU2;A0A087WSP7;A0A0 | Matrin-3 | *Matr3* | -0.61343 | [illegible] |
| H7BWX9;D3Z794;P61957;G3UZX6 | H7BWX9;D3Z794;P61957 | Small ubiquitin-related modifier 2 | *Sumo2* | 0.368381 | [illegible] |
| Q61768;E9QAK5 | Q61768;E9QAK5 | Kinesin-1 heavy chain;Kinesin-like protein | *Kif5b* | -0.833463 | [illegible] |
| P61148;D3Z6H1;D6RCX9;D3Z6E2;D3Z192 | P61148 | Fibroblast growth factor 1 | *Fgf1* | 1.18748 | [illegible] |
| O55222;D3YZA5;A0A1B0GRF6 | O55222;D3YZA5 | Integrin-linked protein kinase | *Ilk* | 0.857973 | [illegible] |
| Q9DCB1;A0A087WSB8;Q9DCB1-2;Q9DCB1-3 | Q9DCB1;A0A087WSB8;Q9DCB1-2;Q9DCB | High mobility group nucleosome-binding domain-containing | *Hmgn3* | -0.555117 | [illegible] |
| P70336;P70336-2;F8VPK5;A0A1Y7VMN0;A0A1Y7VLD1 | P70336;P70336-2;F8VPK5;A0A1Y7VMN0 | Rho-associated protein kinase 2;Rho-associated protein ki | *Rock2* | 0.193367 | [illegible] |
| Q3TMU8;O35098;A0A0R4J1N0;O35098-2 | Q3TMU8;O35098;A0A0R4J1N0;O35098-2 | Dihydropyrimidinase-related protein 4 | *Dpysl4* | -0.248901 | [illegible] |
| Q80XR6;Q20BD0;Q99020 | Q80XR6;Q20BD0;Q99020 | Heterogeneous nuclear ribonucleoprotein A/B | *Hnrnpab* | 1.18057 | [illegible] |
| F6XWD4;Q3UMT1 | F6XWD4;Q3UMT1 | Protein phosphatase 1 regulatory subunit 12C | *Ppp1r12c* | -0.287148 | [illegible] |
| O70579 | O70579 | Peroxisomal membrane protein PMP34 | *Slc25a17* | -0.302516 | [illegible] |
| Q62165 | Q62165 | Dystroglycan;Alpha-dystroglycan;Beta-dystroglycan | *Dag1* | -0.348968 | [illegible] |
| Q3UH59;Q61879;Q5SV64;Q8BXF2 | Q3UH59;Q61879;Q5SV64 | Myosin-10 | *Myh10* | -0.360909 | [illegible] |

| | | | | | | | | |
|---|---|---|---|---|---|---|---|---|
| Z4YK85;M0QWP1;A2ASQ1-3;A2ASQ1-2;A2ASQ1;Z4YJS5 | Z4YK85;M0QWP1;A2ASQ1-3;A2ASQ1-2;A2 | Agrin;Agrin N-terminal 110 kDa subunit;Agrin C-terminal 11 | *Agrn* | -0.120868 | -0.749886 | -0.749886 | -0.749886 | [illegible] |
| O55137;Q9QYR7-2;Q6Q2Z6;Q9QYR7;A0A1Y7VMZ4;Q91YQ6; | O55137 | Acyl-coenzyme A thioesterase 1 | *Acot1* | -0.366347 | -0.366347 | 1.2671 | -0.366347 | [illegible] |
| Q9CZM2;B8JKK2 | Q9CZM2 | 60S ribosomal protein L15 | *Rpl15* | -0.34925 | -0.34925 | -0.34925 | -0.34925 | [illegible] |
| Q9Z1N5;Q8VDW0;G3UXI6;D6RHT5;Q8VDW0-2 | Q9Z1N5;Q8VDW0 | Spliceosome RNA helicase Ddx39b;ATP-dependent RNA h | *Ddx39b;Ddx39a* | -0.222205 | -0.222205 | -0.222205 | -0.222205 | [illegible] |
| Q00612;A3KG36;G3UWD6;P97324 | Q00612;A3KG36 | Glucose-6-phosphate 1-dehydrogenase X;Glucose-6-phos | *G6pdx* | -0.563571 | -0.563571 | -0.563571 | -0.563571 | [illegible] |
| Q9Z1Z0;Q9Z1Z0-2;Q9Z1Z0-4;Q9Z1Z0-3;A0A0J9YUG0 | Q9Z1Z0;Q9Z1Z0-2 | General vesicular transport factor p115 | *Uso1* | 0.357155 | 0.441789 | -0.23429 | -0.139144 | [illegible] |
| A0A3B2WCD8;Q3U0V1;A0A3B2W465;A0A0G2JE55;A0A0G2J | A0A3B2WCD8;Q3U0V1 | Far upstream element-binding protein 2 | *Khsrp* | -0.220558 | 5.26413 | -0.220558 | -0.220558 | [illegible] |
| P58389;A2AWE9;A2AWF0;B7ZDE0;D6RFC2;F6Z6I0 | P58389;A2AWE9;A2AWF0 | Serine/threonine-protein phosphatase 2A activator | *Ppp2r4* | -0.618167 | -0.618167 | -0.618167 | -0.618167 | [illegible] |
| D3YTQ9;P62843 | D3YTQ9;P62843 | 40S ribosomal protein S15 | *Rps15* | -0.235167 | -0.235167 | -0.235167 | -0.235167 | [illegible] |
| Q6ZQ38;D3YWC5 | Q6ZQ38 | Cullin-associated NEDD8-dissociated protein 1 | *Cand1* | -1.45025 | -1.35163 | 0.828808 | 0.757359 | [illegible] |
| Q9WTQ5;Q9WTQ5-2 | Q9WTQ5;Q9WTQ5-2 | A-kinase anchor protein 12 | *Akap12* | -0.846945 | -0.846945 | -0.478426 | -0.846945 | [illegible] |
| A0A0G2JGD2;P07091 | A0A0G2JGD2;P07091 | Protein S100-A4 | *S100a4* | -0.241443 | -0.241443 | -0.241443 | -0.241443 | [illegible] |
| Q9QUM9;E0CXB1;E0CYT2 | Q9QUM9;E0CXB1 | Proteasome subunit alpha type-6 | *Psma6* | -0.299932 | -0.299932 | -0.299932 | -0.299932 | [illegible] |
| P16125;D3Z7F0;A0A0N4SVV8 | P16125;D3Z7F0 | L-lactate dehydrogenase B chain;L-lactate dehydrogenase | *Ldhb* | -0.410336 | -0.184687 | -0.423956 | -0.423956 | [illegible] |
| P47962;D3YYV8 | P47962;D3YYV8 | 60S ribosomal protein L5 | *Rpl5* | -0.317387 | 4.3408 | -0.317387 | -0.317387 | [illegible] |
| P21981;G3UXE8 | P21981 | Protein-glutamine gamma-glutamyltransferase 2 | *Tgm2* | -0.374585 | -0.374585 | -0.374585 | -0.374585 | [illegible] |
| P35564 | P35564 | Calnexin | *Canx* | -0.652608 | -0.652608 | -0.652608 | -0.652608 | [illegible] |
| P49722;Q8BKE0 | P49722;Q8BKE0 | Proteasome subunit alpha type-2 | *Psma2* | 0.74451 | 0.893386 | 3.95112 | -0.440203 | [illegible] |
| E9Q9E1;E9PVC5;Q6NZJ6;E9PVC6;Q6NZJ6-2;A0A0J9YUS5;A | E9Q9E1;E9PVC5;Q6NZJ6;E9PVC6;Q6NZJ | Eukaryotic translation initiation factor 4 gamma 1 | *Eif4g1* | -0.728785 | -0.728785 | -0.728785 | -0.728785 | [illegible] |
| Q91W50;Q8JZN2;A0A0G2JF72;A0A0G2JDJ7;A0A0G2JE62 | Q91W50;Q8JZN2;A0A0G2JF72 | Cold shock domain-containing protein E1 | *Csde1* | -0.525414 | -0.525414 | -0.525414 | -0.525414 | [illegible] |
| P58404-2;P58404;V9GXE7;V9GXT1;V9GXI9 | P58404-2;P58404;V9GXE7 | Striatin-4 | *Strn4* | -0.176777 | -0.176777 | -0.176777 | -0.176777 | [illegible] |
| Q60963;E9Q330;E9Q6J0;E9Q4T5 | Q60963;E9Q330 | Platelet-activating factor acetylhydrolase | *Pla2g7* | -0.456063 | 2.5754 | -0.456063 | -0.456063 | [illegible] |
| Q80UG5-3;A2A6U3;Q80UG5;Q80UG5-2;A2A6U5;A8Y5D3;REV | Q80UG5-3;A2A6U3;Q80UG5;Q80UG5-2 | Septin-9 | *Sep-09* | -0.546828 | -0.546828 | -0.546828 | -0.546828 | [illegible] |
| P10833;A0A1B0GRG1;A0A1B0GRT5 | P10833 | Ras-related protein R-Ras | *Rras* | 1.63278 | 1.97032 | 0.765401 | 0.742314 | [illegible] |
| A0A0N4SW28;Q9DAS9;A0A0N4SVT3 | A0A0N4SW28;Q9DAS9;A0A0N4SVT3 | Guanine nucleotide-binding protein G(I)/G(S)/G(O) subunit | *Gng12* | -0.176777 | -0.176777 | -0.176777 | -0.176777 | [illegible] |
| Q9R1P1;D3YUM8 | Q9R1P1 | Proteasome subunit beta type-3 | *Psmb3* | -0.176777 | -0.176777 | -0.176777 | -0.176777 | [illegible] |
| A0A1L1STE6;Q9D6R2;Q9D6R2-2 | A0A1L1STE6;Q9D6R2;Q9D6R2-2 | Isocitrate dehydrogenase [NAD] subunit alpha, mitochondri | *Idh3a* | -0.463739 | -0.463739 | -0.463739 | -0.463739 | [illegible] |
| Q3UKN6;P81117;S4R2R9 | Q3UKN6;P81117 | Nucleobindin-2;Nesfatin-1 | *Nucb2* | -0.252992 | -0.252992 | -0.252992 | -0.252992 | [illegible] |
| [illegible] | [illegible] | [illegible] | [illegible] | [illegible] | [illegible] | [illegible] | [illegible] | [illegible] |

[illegible]

| | | | | | | | | |
|---|---|---|---|---|---|---|---|---|
| E9PYI8;Q9JMA1 | E9PYI8;Q9JMA1 | Ubiquitin carboxyl-terminal hydrolase;Ubiquitin carboxyl-ter | *Usp14* | -0.346337 | -0.346337 | -0.346337 | 3.4644 | [illegible] |
| E9PYN1;Q8R5M8-4;Q8R5M8-3;Q8R5M8-2;Q8R5M8;Q8R5M8-[illegible] | E9PYN1;Q8R5M8-4;Q8R5M8-3;Q8R5M8-2 | Cell adhesion molecule 1 | *Cadm1* | 1.26717 | -0.635606 | 1.43651 | 1.2159 | [illegible] |
| E9PYT0;P97393;A0A1W2P6H9 | E9PYT0;P97393 | Rho GTPase-activating protein 5 | *Arhgap5* | -0.176777 | -0.176777 | -0.176777 | -0.176777 | [illegible] |
| E9PZ43;A0A0G2JDN7;P27546-4 | E9PZ43;A0A0G2JDN7;P27546-4 | Microtubule-associated protein;Microtubule-associated pro | *Map4* | 2.82332 | 3.7211 | -0.34398 | -0.34398 | [illegible] |
| E9PZD8;Q61147;G3UXG1;G3UWP5;G3UZ53 | E9PZD8;Q61147 | Ceruloplasmin | *Cp* | -0.53669 | -0.53669 | -0.53669 | -0.53669 | [illegible] |
| E9PZF0;Q01768 | E9PZF0;Q01768 | Nucleoside diphosphate kinase;Nucleoside diphosphate kir | *Gm20390;Nme2* | 3.17339 | 4.0308 | 0.708689 | 0.759354 | [illegible] |
| E9Q0G1;Q9CQJ6 | E9Q0G1;Q9CQJ6 | Density-regulated protein | *Denr* | -0.411821 | -0.411821 | -0.411821 | -0.411821 | [illegible] |
| F8WGN6;E9Q0J5;Q9QXL2-3;Q9QXL2-2;Q9QXL2-4;Q9QXL2;[illegible] | F8WGN6;E9Q0J5;Q9QXL2-3;Q9QXL2-2;Q[illegible] | Kinesin-like protein;Kinesin-like protein KIF21A | *Kif21a* | -0.176777 | -0.176777 | -0.176777 | -0.176777 | [illegible] |
| E9Q1J7;Q99MN9;D3YZC1;A0A087WQV1 | E9Q1J7;Q99MN9 | Propionyl-CoA carboxylase beta chain, mitochondrial | *Pccb* | 1.79151 | 1.41881 | -0.439072 | -0.0762718 | [illegible] |
| E9Q1S3;Q01405 | E9Q1S3;Q01405 | Protein transport protein Sec23A | *Sec23a* | -0.319981 | -0.319981 | -0.319981 | -0.319981 | [illegible] |
| E9Q1X8;O08532-3;O08532-2;O08532-4;O08532-5;O08532;Q8[illegible] | E9Q1X8;O08532-3;O08532-2;O08532-4;O[illegible] | Voltage-dependent calcium channel subunit alpha-2/delta-1 | *Cacna2d1* | 5.48008 | -0.176777 | -0.176777 | -0.176777 | [illegible] |
| E9Q242;P54822;E9Q3T7;A0A140LHU1;A0A0G2JFI8;E9Q0A0 | E9Q242;P54822;E9Q3T7 | Adenylosuccinate lyase | *Adsl* | -0.305263 | -0.305263 | 2.57925 | -0.305263 | [illegible] |
| E9Q3E2;Q8CC35-2;Q8CC35-3;Q8CC35 | E9Q3E2;Q8CC35-2;Q8CC35-3;Q8CC35 | Synaptopodin | *Synpo* | -0.318711 | 4.70817 | -0.318711 | -0.318711 | [illegible] |
| E9Q3M0;Q62432-2;Q62432;A2CG44;Q8BUN5;F7BUP2;E9Q5P[illegible] | E9Q3M0;Q62432-2;Q62432;A2CG44;Q8BU[illegible] | Mothers against decapentaplegic homolog;Mothers against | *Smad2;Smad3* | -0.33211 | -0.33211 | -0.33211 | -0.33211 | [illegible] |
| E9Q586;E9Q3M3;O08788-2;O08788;D3YX34;D3Z2M9;D3YYG[illegible] | E9Q586;E9Q3M3;O08788-2;O08788;D3YX[illegible] | Dynactin subunit 1 | *Dctn1* | -0.752509 | -0.752509 | -0.752509 | -0.752509 | [illegible] |
| E9Q3P4;E9PY14 | E9Q3P4;E9PY14 | | *Cenpf* | -0.253487 | -0.253487 | -0.253487 | -0.253487 | [illegible] |
| E9Q4G8;E9Q3Q6;Q61490;Q5MPX5;F6QH25 | E9Q4G8;E9Q3Q6;Q61490 | CD166 antigen | *Alcam* | -0.299335 | -0.299335 | -0.299335 | -0.299335 | [illegible] |
| E9Q4D0;F6XJP7;H3BJL4;H3BLN2;Q04592-2;Q04592 | E9Q4D0;F6XJP7;H3BJL4 | | *Pcsk6* | -0.176777 | -0.176777 | -0.176777 | -0.176777 | [illegible] |
| E9Q4M4;Q91VN4 | E9Q4M4;Q91VN4 | MICOS complex subunit Mic25 | *Chchd6* | -0.642728 | -0.642728 | 0.149983 | -0.642728 | [illegible] |
| E9Q5B5;O08528;E9Q9M6 | E9Q5B5;O08528 | Hexokinase;Hexokinase-2 | *Hk2* | -1.19901 | -0.0976928 | -0.035662 | -1.19901 | [illegible] |
| F8WH23;E9Q5D9;Q61581 | F8WH23;E9Q5D9;Q61581 | Insulin-like growth factor-binding protein 7 | *Igfbp7* | -0.523401 | -0.523401 | -0.523401 | -0.523401 | [illegible] |
| E9Q634;G3V011;Q8CG29;P70248;A0A1L1STM1 | E9Q634 | Unconventional myosin-Ie | *Myo1e* | -0.176777 | -0.176777 | -0.176777 | -0.176777 | [illegible] |
| E9Q6R7;A0A1W2P7C0;Q61636;A0A1W2P8G5;A0A1W2P8A5;A[illegible] | E9Q6R7 | | *Utrn* | -0.403323 | -0.403323 | -0.403323 | -0.403323 | [illegible] |
| E9Q704;F8VQA4;P97467;F6TGI4;F6T0D3 | E9Q704;F8VQA4;P97467 | Peptidyl-glycine alpha-amidating monooxygenase;Peptidylg[illegible] | *Pam* | 1.41784 | -0.520226 | 1.52601 | 3.09896 | [illegible] |
| E9Q718;Q9R0B9 | E9Q718;Q9R0B9 | Procollagen-lysine,2-oxoglutarate 5-dioxygenase 2 | *Plod2* | -0.290731 | -0.290731 | -0.290731 | -0.290731 | [illegible] |
| G3X9Y5;E9Q735;Q8VDS1;E9Q442 | G3X9Y5;E9Q735 | Ubiquitin conjugation factor E4 A | *Ube4a* | -0.176777 | -0.176777 | -0.176777 | -0.176777 | [illegible] |
| E9Q740;F8VQC1 | E9Q740;F8VQC1 | Signal recognition particle subunit SRP72 | *Srp72* | -0.176777 | -0.176777 | -0.176777 | -0.176777 | [illegible] |
| E9Q7A5;E9Q8R5;Q99JY4 | E9Q7A5;E9Q8R5;Q99JY4 | TraB domain-containing protein | *Trabd* | -0.435877 | 0.899609 | -0.435877 | -0.435877 | [illegible] |
| [illegible] | [illegible] | [illegible] | [illegible] | [illegible] | [illegible] | [illegible] | [illegible] | [illegible] |

| | | | | | |
|---|---|---|---|---|---|
| O70435;E0CX62;F7BWF9;E0CZ34;F8WH02;E0CYL6 | O70435;E0CX62 | Proteasome subunit alpha type-3 | *Psma3* | -0.668198 | [illegible] |
| Q8BH40;O70439 | Q8BH40;O70439 | Syntaxin-7 | *Stx7* | 3.53327 | [illegible] |
| O70456 | O70456 | 14-3-3 protein sigma | *Sfn* | -0.274629 | [illegible] |
| O70475;D3YXP9 | O70475 | UDP-glucose 6-dehydrogenase | *Ugdh* | -0.802749 | [illegible] |
| O70503;O70503-2 | O70503 | Very-long-chain 3-oxoacyl-CoA reductase | *Hsd17b12* | -0.425187 | [illegible] |
| O88322;A0A286YCQ5 | O88322;A0A286YCQ5 | Nidogen-2 | *Nid2* | -0.176777 | [illegible] |
| O88325 | O88325 | | *Naglu* | -0.176777 | [illegible] |
| O88327;B1AZA7 | O88327 | Alpha-catulin | *Ctnnal1* | -0.647215 | [illegible] |
| O88342;A0A0J9YU05 | O88342 | WD repeat-containing protein 1 | *Wdr1* | 0.854828 | [illegible] |
| O88343;A7E1Z5;O88343-2 | O88343;A7E1Z5;O88343-2 | Electrogenic sodium bicarbonate cotransporter 1;Anion exc | *Slc4a4* | -0.357708 | [illegible] |
| O88531;B1B0P8;B1B0P9 | O88531 | Palmitoyl-protein thioesterase 1 | *Ppt1* | -0.176777 | [illegible] |
| O88543;F6YCA7;D3Z036 | O88543 | COP9 signalosome complex subunit 3 | *Cops3* | 1.52955 | [illegible] |
| O88544;D3YV99;F6QTS1;D3Z1R9;D6RG47 | O88544;D3YV99;F6QTS1;D3Z1R9 | COP9 signalosome complex subunit 4 | *Cops4* | 1.02345 | [illegible] |
| O88685;B7ZCF1;A2AGN7;A0A087WPH7;F6Q2E3 | O88685;B7ZCF1;A2AGN7;A0A087WPH7 | 26S protease regulatory subunit 6A | *Psmc3* | -0.202182 | [illegible] |
| O88696 | O88696 | ATP-dependent Clp protease proteolytic subunit, mitochonc | *Clpp* | -0.176777 | [illegible] |
| O88952;Q8JZS0;Q3TUM0;O88951 | O88952 | Protein lin-7 homolog C | *Lin7c* | -0.297706 | [illegible] |
| O88958;D3Z0R5 | O88958;D3Z0R5 | Glucosamine-6-phosphate isomerase 1 | *Gnpda1* | -0.254121 | [illegible] |
| O88968 | O88968 | Transcobalamin-2 | *Tcn2* | -0.45436 | [illegible] |
| O89001 | O89001 | Carboxypeptidase D | *Cpd* | -0.35895 | [illegible] |
| O89023 | O89023 | Tripeptidyl-peptidase 1 | *Tpp1* | -0.310948 | [illegible] |
| O89051;A0A2I3BQ30;A0A2I3BPW7 | O89051 | Integral membrane protein 2B;BRI2, membrane form;BRI2 i | *Itm2b* | -0.319931 | [illegible] |
| O89053;G3UYK8;A0A0U1RPY8;D3YW57;G3UX53;D3YXM2 | O89053;G3UYK8 | Coronin-1A;Coronin | *Coro1a* | -0.528181 | [illegible] |
| O89079;D3Z315;F6YFR7;E9Q6I5 | O89079;D3Z315 | Coatomer subunit epsilon | *Cope* | 0.267271 | [illegible] |
| Q8BG13;O89086;S4R2M6 | Q8BG13;O89086 | RNA-binding protein 3 | *Rbm3* | -0.176777 | [illegible] |
| O89112;B2KGR2 | O89112;B2KGR2 | LanC-like protein 1 | *Lancl1* | -0.417901 | [illegible] |
| P00493 | P00493 | Hypoxanthine-guanine phosphoribosyltransferase | *Hprt1* | 4.00622 | [illegible] |
| P01029 | P01029 | Complement C4-B;Complement C4 beta chain;Complement | *C4b* | -0.389476 | [illegible] |
| P01899;P01897;P01896;E9PWT4;E9QJR9;P14428;P14429;P03 | P01899;P01897 | H-2 class I histocompatibility antigen, D-B alpha chain;H-2 | *H2-D1;H2-L* | -0.651668 | [illegible] |
| P04627;B1AUN8;E9Q0P8;Q8CAD1;B1AUP0;A0A0N4SWH5;A0A | P04627;B1AUN8;E9Q0P8;Q8CAD1 | Serine/threonine-protein kinase A-Raf | *Araf* | -0.253936 | [illegible] |
| P05063 | P05063 | Fructose-bisphosphate aldolase C | *Aldoc* | -0.258953 | [illegible] |
| P05201;F7ALS6 | P05201 | Aspartate aminotransferase, cytoplasmic | *Got1* | -0.495476 | [illegible] |
| P05202 | P05202 | Aspartate aminotransferase, mitochondrial | *Got2* | -0.389927 | [illegible] |
| P06537-5;P06537-3;P06537;E9PUR6;P06537-4;P06537-2;E9PY | P06537-5;P06537-3;P06537;E9PUR6;P065 | Glucocorticoid receptor | *Nr3c1* | -1.0304 | [illegible] |
| P06745;A0A0U1RP97;A0A0U1RQ72;A0A0U1RQ18 | P06745 | Glucose-6-phosphate isomerase | *Gpi* | 0.879847 | [illegible] |
| P06837 | P06837 | Neuromodulin | *Gap43* | -0.643175 | [illegible] |
| P07309 | P07309 | Transthyretin | *Ttr* | -0.117116 | [illegible] |
| P07356;B0V2N7;B0V2N8;B0V2N5 | P07356;B0V2N7;B0V2N8;B0V2N5 | Annexin A2;Annexin | *Anxa2* | -0.466175 | [illegible] |
| P07724 | P07724 | Serum albumin | *Alb* | -1.12228 | [illegible] |
| P07901;B7ZC50;A2A6A2;B7ZC49;REV__Q8BL66 | P07901 | Heat shock protein HSP 90-alpha | *Hsp90aa1* | -0.430246 | [illegible] |
| P08121;A0A087WPS3;A0A087WPJ5 | P08121 | Collagen alpha-1(III) chain | *Col3a1* | -0.383917 | [illegible] |
| P08207 | P08207 | Protein S100-A10 | *S100a10* | -0.200545 | [illegible] |
| P08228 | P08228 | Superoxide dismutase [Cu-Zn] | *Sod1* | -0.244236 | [illegible] |
| P08249;A0A0G2JF23;A0A0G2JGY4 | P08249 | Malate dehydrogenase, mitochondrial | *Mdh2* | -0.249555 | [illegible] |
| P08752;A0A0A6YWA9;B2RSH2;A0A0A6YXN0;A0A0A6YXC2;A2A | P08752;A0A0A6YWA9 | Guanine nucleotide-binding protein G(i) subunit alpha-2 | *Gnai2* | 4.02949 | [illegible] |
| P09055;P09055-2 | P09055;P09055-2 | Integrin beta-1 | *Itgb1* | -0.196643 | [illegible] |
| P09103;E9Q8G8 | P09103 | Protein disulfide-isomerase | *P4hb* | -0.436927 | [illegible] |
| P09405;A0A087WRM5 | P09405 | Nucleolin | *Ncl* | -0.674906 | [illegible] |
| P09470;F6QCP8;P09470-2 | P09470;F6QCP8;P09470-2 | Angiotensin-converting enzyme;Angiotensin-converting enz | *Ace* | -0.253873 | [illegible] |
| P09581 | P09581 | Macrophage colony-stimulating factor 1 receptor | *Csf1r* | -0.293208 | [illegible] |
| P10107 | P10107 | Annexin A1 | *Anxa1* | 1.38431 | [illegible] |
| P10493 | P10493 | Nidogen-1 | *Nid1* | 0.630338 | [illegible] |
| P10639 | P10639 | Thioredoxin | *Txn* | 0.148949 | [illegible] |
| P10852;P10852-2;A0A0U1RP98;A0A0U1RPK4;A0A0U1RPL8;A( | P10852;P10852-2;A0A0U1RP98 | 4F2 cell-surface antigen heavy chain | *Slc3a2* | -0.176777 | [illegible] |
| P10922 | P10922 | Histone H1.0;Histone H1.0, N-terminally processed | *H1f0* | -0.254103 | [illegible] |
| P11499;E9Q3D6;E9PX27;E9Q0C3;D3Z1R1 | P11499 | Heat shock protein HSP 90-beta | *Hsp90ab1* | -0.203437 | [illegible] |
| P11531;A2A9Z1;A2A9Z2 | P11531;A2A9Z1;A2A9Z2 | Dystrophin | *Dmd* | -0.301538 | [illegible] |
| P11688 | P11688 | Integrin alpha-5;Integrin alpha-5 heavy chain;Integrin alpha- | *Itga5* | 3.30934 | [illegible] |
| Q542I8;P11835;M0QWA7;D3YYP8 | Q542I8;P11835;M0QWA7 | Integrin beta;Integrin beta-2 | *Itgb2* | -0.544832 | [illegible] |
| P11881-8;P11881-7;P11881-6;P11881-5;P11881-4;P11881-2;P1 | P11881-8;P11881-7;P11881-6;P11881-5;P1 | Inositol 1,4,5-trisphosphate receptor type 1 | *Itpr1* | 3.32076 | [illegible] |
| P11983;P11983-2;F2Z483;A0A3B2WDE2 | P11983;P11983-2 | T-complex protein 1 subunit alpha | *Tcp1* | -0.236949 | [illegible] |
| P12265;Q99KJ6;D3YY48 | P12265 | Beta-glucuronidase | *Gusb* | -0.143713 | [illegible] |
| P12382;A0A1W2P7T1 | P12382 | ATP-dependent 6-phosphofructokinase, liver type | *Pfkl* | -1.05812 | [illegible] |
| P12970;V9GX35 | P12970 | 60S ribosomal protein L7a | *Rpl7a* | -0.80145 | [illegible] |
| P13011;Q99PL7;P13516;Q6T707 | P13011;Q99PL7;P13516 | Acyl-CoA desaturase 2;Acyl-CoA desaturase 1 | *Scd2;Scd3;Scd1* | -0.176777 | [illegible] |
| P13020-2;P13020;A6PWS5 | P13020-2;P13020 | Gelsolin | *Gsn* | -0.331295 | [illegible] |
| P13597-2;P13597 | P13597-2;P13597 | Intercellular adhesion molecule 1 | *Icam1* | -0.254058 | [illegible] |
| P14094;A0A0A6YX05 | P14094 | Sodium/potassium-transporting ATPase subunit beta-1 | *Atp1b1* | -0.624207 | [illegible] |
| P14106 | P14106 | Complement C1q subcomponent subunit B | *C1qb* | -0.176777 | [illegible] |
| P14148;F6XI62 | P14148;F6XI62 | 60S ribosomal protein L7 | *Rpl7* | -0.389554 | [illegible] |
| P14152;B1ATQ3 | P14152 | Malate dehydrogenase, cytoplasmic | *Mdh1* | -0.793047 | [illegible] |
| P14206;A0A1L1SUK3;A0A1L1SRW0 | P14206;A0A1L1SUK3 | 40S ribosomal protein SA | *Rpsa* | -0.176777 | [illegible] |
| P14576;E9PXC0;A0A1Y7VJJ0;A0A1W2P809;P14576-2 | P14576;E9PXC0 | Signal recognition particle 54 kDa protein | *Srp54;Srp54c* | -0.501325 | [illegible] |
| P14869;D3YVM5;S4R1N1 | P14869;D3YVM5;S4R1N1 | 60S acidic ribosomal protein P0 | *Rplp0* | -0.351212 | [illegible] |
| P14873 | P14873 | Microtubule-associated protein 1B;MAP1B heavy chain;MA | *Map1b* | -0.311969 | [illegible] |
| P14901 | P14901 | Heme oxygenase 1 | *Hmox1* | -0.647514 | [illegible] |
| P15105;D3YVK1;D3Z121 | P15105 | Glutamine synthetase | *Glul* | 1.69268 | [illegible] |
| P15209-4;P15209-3;P15209;A0A286YDA0;A0A286YCV8;A0A28 | P15209-4;P15209-3;P15209;A0A286YDA0;A | BDNF/NT-3 growth factors receptor | *Ntrk2* | -0.306666 | [illegible] |
| P15626;D3YX76;Q8R5I6;F6Y363;D3YVP9;G5E8M7;O35660 | P15626;D3YX76 | Glutathione S-transferase Mu 2 | *Gstm2* | -0.237277 | [illegible] |
| P15864 | P15864 | Histone H1.2 | *Hist1h1c* | -0.419661 | [illegible] |
| P16045;A0A2R8VHJ0 | P16045 | Galectin-1 | *Lgals1* | -0.453337 | [illegible] |
| P16330-2;P16330 | P16330-2;P16330 | 2,3-cyclic-nucleotide 3-phosphodiesterase | *Cnp* | -0.253321 | [illegible] |
| P16332;A0A3B2WCT0 | P16332 | Methylmalonyl-CoA mutase, mitochondrial | *Mut* | 2.23699 | [illegible] |
| P17427;A0A0J9YUA7;A0A140LHG0 | P17427 | AP-2 complex subunit alpha-2 | *Ap2a2* | -0.334896 | [illegible] |
| P17710-3;G3UVV4;P17710-4;P17710;P17710-2;D3YYR4;D3Z3( | P17710-3;G3UVV4;P17710-4;P17710;P177 | Hexokinase-1;Hexokinase | *Hk1* | -0.501447 | [illegible] |
| Q61696;P17879 | Q61696;P17879 | Heat shock 70 kDa protein 1A;Heat shock 70 kDa protein 1 | *Hspa1a;Hspa1b* | -0.554136 | [illegible] |
| P18242;F8WIR1;F6Y6L6;A0A1B0GT66 | P18242;F8WIR1;F6Y6L6;A0A1B0GT66 | Cathepsin D | *Ctsd* | -0.522535 | [illegible] |
| P18406 | P18406 | Protein CYR61 | *Cyr61* | -0.176777 | [illegible] |
| P18872;D3Z2M7;F7BLT7;F6W1B2;F6WC15 | P18872;D3Z2M7 | Guanine nucleotide-binding protein G(o) subunit alpha | *Gnao1* | -0.246346 | [illegible] |
| P19157;P46425;Q8VC73;F6RWR5 | P19157;P46425 | Glutathione S-transferase P 1;Glutathione S-transferase P | *Gstp1;Gstp2* | -0.176777 | [illegible] |
| P19253;A0A1B0GSQ6;A0A1B0GS68;A0A1B0GTA1;A0A1B0GS | P19253;A0A1B0GSQ6;A0A1B0GS68;A0A1E | 60S ribosomal protein L13a | *Rpl13a* | -0.248941 | [illegible] |
| P19324;A0A140LHR4;A0A140LHK0;A0A140LHD6 | P19324 | Serpin H1 | *Serpinh1* | 2.55493 | [illegible] |
| Q9D881;P19536;F7C106;A0A0A6YVR0 | Q9D881;P19536 | Cytochrome c oxidase subunit 5B, mitochondrial | *Cox5b* | -0.310764 | [illegible] |
| P19783;M0QWX7;M0QWX2;D6RG40 | P19783;M0QWX7 | Cytochrome c oxidase subunit 4 isoform 1, mitochondrial | *Cox4i1* | -0.253966 | [illegible] |
| P20060 | P20060 | Beta-hexosaminidase subunit beta | *Hexb* | 0.1025 | [illegible] |
| P21278 | P21278 | Guanine nucleotide-binding protein subunit alpha-11 | *Gna11* | -0.176777 | [illegible] |
| P21279 | P21279 | Guanine nucleotide-binding protein G(q) subunit alpha | *Gnaq* | 3.7072 | [illegible] |
| P21956-2;P21956 | P21956-2;P21956 | Lactadherin | *Mfge8* | -0.438954 | [illegible] |
| Q544X6;P22315 | Q544X6;P22315 | Ferrochelatase;Ferrochelatase, mitochondrial | *Fech* | -0.176777 | [illegible] |
| Q8CBB7;P22892;O88512 | Q8CBB7;P22892 | AP-1 complex subunit gamma-1 | *Ap1g1* | -0.306346 | [illegible] |
| P23116;REV__G3UZX7;REV__G3UYR9;REV__G3UWR2;REV_ | P23116 | Eukaryotic translation initiation factor 3 subunit A | *Eif3a* | -0.176777 | [illegible] |
| Q9DCC5;P23198;D3Z1A9;D3Z313 | Q9DCC5;P23198 | Chromobox protein homolog 3 | *Cbx3* | -0.489541 | [illegible] |
| Q543K9;P23492;A0A2I3BQH2;Q9D8C9;A0A2I3BS22 | Q543K9;P23492;A0A2I3BQH2 | Purine nucleoside phosphorylase | *Pnp* | -0.514926 | [illegible] |
| P23819-4;P23819-3;E9QKC0;G5E8H1;P23819;A0A0A6YW90;P2 | P23819-4;P23819-3;E9QKC0;G5E8H1;P23 | Glutamate receptor 2 | *Gria2* | 0.456943 | [illegible] |
| P23927;A0A1L1SRG5 | P23927;A0A1L1SRG5 | Alpha-crystallin B chain | *Cryab* | 1.19871 | [illegible] |
| P24527 | P24527 | Leukotriene A-4 hydrolase | *Lta4h* | -0.32822 | [illegible] |
| P24549;A0A286YDG6;A0A286YCZ0 | P24549 | Retinal dehydrogenase 1 | *Aldh1a1* | 4.55362 | [illegible] |
| P25444;D3YVC1;D3YWJ3;D3Z536 | P25444;D3YVC1;D3YWJ3 | 40S ribosomal protein S2 | *Rps2* | -0.279572 | [illegible] |
| P26039;A2AIM2;F6SX70;F6S1V7 | P26039 | Talin-1 | *Tln1* | -0.588674 | [illegible] |
| P26043;Q7TSG6 | P26043;Q7TSG6 | Radixin | *Rdx* | -0.462557 | [illegible] |
| Q8C483;P26638;A2AFS0;A2AFS1 | Q8C483;P26638;A2AFS0;A2AFS1 | Serine--tRNA ligase, cytoplasmic | *Sars* | -0.290859 | [illegible] |
| P26883;F6X9I3 | P26883 | Peptidyl-prolyl cis-trans isomerase FKBP1A | *Fkbp1a* | -0.458573 | [illegible] |
| P27546-2;P27546;P27546-3;A0A0G2JFH2;Q78TF3 | P27546-2;P27546;P27546-3 | Microtubule-associated protein 4 | *Map4* | 3.42914 | [illegible] |
| P27612;F7D1R5 | P27612 | Phospholipase A-2-activating protein | *Plaa* | -0.176777 | [illegible] |
| P27659;A0A087WQK0;A0A2R8VHN4;A0A087WNS0;Q9CQ09;E | P27659 | 60S ribosomal protein L3 | *Rpl3* | -0.518606 | [illegible] |
| P28271;Q811J3 | P28271 | Cytoplasmic aconitate hydratase | *Aco1* | -0.245529 | [illegible] |
| P28301 | P28301 | Protein-lysine 6-oxidase | *Lox* | -0.25119 | [illegible] |
| P28474;A0A0G2JGS3;Q64437 | P28474 | Alcohol dehydrogenase class-3 | *Adh5* | 1.38814 | [illegible] |
| P28653 | P28653 | Biglycan | *Bgn* | -0.435758 | [illegible] |
| P28654 | P28654 | Decorin | *Dcn* | -0.266605 | [illegible] |
| Q3TF41;Q8BSH9;P28656 | Q3TF41;Q8BSH9;P28656 | Nucleosome assembly protein 1-like 1 | *Nap1l1* | 4.17016 | [illegible] |
| P28658 | P28658 | Ataxin-10 | *Atxn10* | 0.0770399 | [illegible] |
| P28661-4;P28661-2;P28661;P28661-3;Q5ND13;Q5ND18;Q5ND | P28661-4;P28661-2;P28661;P28661-3 | Septin-4 | *Sep-04* | -0.24017 | [illegible] |
| P28667 | P28667 | MARCKS-related protein | *Marcksl1* | -0.856323 | [illegible] |
| P28740-2;E0CZ72;P28740;P28740-1;F8VQ42;Q8C0N1 | P28740-2;E0CZ72;P28740;P28740-1;F8VQ | Kinesin-like protein KIF2A;Kinesin-like protein | *Kif2a* | -0.24982 | [illegible] |
| Q68FM4;P28828;A0A286YDL1;A0A286YCW9;B1AQN2;Q99M8( | Q68FM4;P28828;A0A286YDL1 | Protein-tyrosine-phosphatase;Receptor-type tyrosine-prote | *Ptprm* | -0.176777 | [illegible] |
| P29268;H3BJW0;H3BK14 | P29268;H3BJW0 | Connective tissue growth factor | *Ctgf* | -0.250457 | [illegible] |
| P29341;Q9D4E6;Q62029;A0A2I3BR37;F6ZAX1;Q8C7D3;A2A5N | P29341 | Polyadenylate-binding protein 1 | *Pabpc1* | 0.774743 | [illegible] |
| P29758 | P29758 | Ornithine aminotransferase, mitochondrial | *Oat* | -0.211608 | [illegible] |
| P30285;E9Q9E4;E9PZX7;D3YWB0;D6RHS5 | P30285;E9Q9E4;E9PZX7 | Cyclin-dependent kinase 4 | *Cdk4* | 1.41204 | [illegible] |
| P30416;F6S2D5;F7CAT1 | P30416 | Peptidyl-prolyl cis-trans isomerase FKBP4;Peptidyl-prolyl ci | *Fkbp4* | -0.429188 | [illegible] |
| Q3UZG4;P31230;A0A0G2JEU9;A0A0G2JDV0;A0A0G2JDW6;A( | Q3UZG4;P31230 | Aminoacyl tRNA synthase complex-interacting multifunctior | *Aimp1* | -0.33827 | [illegible] |
| P31786;Q4VWZ5;D3Z563;M0QWU8 | P31786;Q4VWZ5;D3Z563 | Acyl-CoA-binding protein | *Dbi* | -0.477565 | [illegible] |
| P32020;P32020-2;A2APS4 | P32020;P32020-2 | Non-specific lipid-transfer protein | *Scp2* | -0.176777 | [illegible] |
| P32233;F7BQ65 | P32233 | Developmentally-regulated GTP-binding protein 1 | *Drg1* | -0.303463 | [illegible] |
| P32261 | P32261 | Antithrombin-III | *Serpinc1* | -0.176777 | [illegible] |
| P32921-2;P32921 | P32921-2;P32921 | Tryptophan--tRNA ligase, cytoplasmic;T1-TrpRS;T2-TrpRS | *Wars* | -0.543623 | [illegible] |
| P34022;H7BX22 | P34022;H7BX22 | Ran-specific GTPase-activating protein | *Ranbp1* | -0.400768 | [illegible] |
| P34152;P34152-4;P34152-2;P34152-6;P34152-5;P34152-7;P34 | P34152;P34152-4;P34152-2;P34152-6;P34 | Focal adhesion kinase 1 | *Ptk2* | -0.250045 | [illegible] |
| P35235-2;P35235 | P35235-2;P35235 | Tyrosine-protein phosphatase non-receptor type 11 | *Ptpn11* | 1.18187 | [illegible] |
| Q8C266;P35278;A2A5F5;A2A5F6 | Q8C266;P35278 | Ras-related protein Rab-5C | *Rab5c* | -0.176777 | [illegible] |
| P35282;A0A1W2P6Z4 | P35282 | Ras-related protein Rab-21 | *Rab21* | -0.176777 | [illegible] |
| P35293 | P35293 | Ras-related protein Rab-18 | *Rab18* | -0.176777 | [illegible] |
| P35505;D3Z2R9 | P35505 | Fumarylacetoacetase | *Fah* | 0.923386 | [illegible] |
| Q543V3;P35569 | Q543V3;P35569 | Insulin receptor substrate 1 | *Irs1* | -0.254113 | [illegible] |
| P35585;D3YZ71;S4R1Q4;A0A1L1SUA0;A0A0R4IZX6;A0A0R4J | P35585;D3YZ71 | AP-1 complex subunit mu-1 | *Ap1m1* | -0.531221 | [illegible] |
| P35762;A0A140LJL0 | P35762;A0A140LJL0 | CD81 antigen | *Cd81* | -0.176777 | [illegible] |
| P35802 | P35802 | Neuronal membrane glycoprotein M6-a | *Gpm6a* | -0.312681 | [illegible] |
| P35980;A0A1B0GSS8;A0A1B0GQU8;A0A1B0GSA8;A0A1B0GS | P35980;A0A1B0GSS8;A0A1B0GQU8;A0A1I | 60S ribosomal protein L18 | *Rpl18* | -0.341196 | [illegible] |
| P36423;F6SG79 | P36423 | Thromboxane-A synthase | *Tbxas1* | -0.295941 | [illegible] |
| P36916;P36916-2 | P36916;P36916-2 | Guanine nucleotide-binding protein-like 1 | *Gnl1* | -0.559745 | [illegible] |
| P37040;Q05DV1;E9PVT9;E9Q997 | P37040;Q05DV1 | NADPH--cytochrome P450 reductase | *Por* | -0.649208 | [illegible] |
| P38647 | P38647 | Stress-70 protein, mitochondrial | *Hspa9* | -0.554985 | [illegible] |
| P39038;Q80ZV4 | P39038;Q80ZV4 | Cadherin-4 | *Cdh4* | -0.363373 | [illegible] |
| P39054-2;G3UZZ3 | P39054-2;G3UZZ3 | | *Dnm2* | -0.604688 | [illegible] |
| P40124;B1ARS0 | P40124 | Adenylyl cyclase-associated protein 1 | *Cap1* | -0.646902 | [illegible] |
| P40142;A0A286YE28;E0CY51;A0A0R4J2A3;Q9D4D4 | P40142;A0A286YE28 | Transketolase | *Tkt* | -0.496522 | [illegible] |
| P40240 | P40240 | CD9 antigen | *Cd9* | -0.176777 | [illegible] |
| P41233 | P41233 | ATP-binding cassette sub-family A member 1 | *Abca1* | -0.253841 | [illegible] |
| P41241;A0A1L1SQQ5;D3YVQ8;A0A1L1SR46;A0A1L1STA1;D6F | P41241 | Tyrosine-protein kinase CSK | *Csk* | -0.176777 | [illegible] |
| P42128;A0A0G2JG71 | P42128;A0A0G2JG71 | Forkhead box protein K1 | *Foxk1* | -0.314574 | [illegible] |
| P42208;E9Q3V6;F6WYM0;D3YYB1;D3Z3C0;D3Z1S1;F6UKN5;[ | P42208;E9Q3V6;F6WYM0;D3YYB1;D3Z3C( | Septin-2 | *Sep-02* | -0.251192 | [illegible] |
| P42669 | P42669 | Transcriptional activator protein Pur-alpha | *Pura* | -0.176777 | [illegible] |
| P43274 | P43274 | Histone H1.4 | *Hist1h1e* | -0.176777 | [illegible] |
| P43275 | P43275 | Histone H1.1 | *Hist1h1a* | -0.25411 | [illegible] |
| P43276 | P43276 | Histone H1.5 | *Hist1h1b* | -0.438997 | [illegible] |
| P43277 | P43277 | Histone H1.3 | *Hist1h1d* | 0.893834 | [illegible] |
| P45591;A0A1Y7VJ71 | P45591 | Cofilin-2 | *Cfl2* | 5.48008 | [illegible] |
| P45878 | P45878 | Peptidyl-prolyl cis-trans isomerase FKBP2 | *Fkbp2* | 3.20313 | [illegible] |
| P46460;G3UX86;G3UX98 | P46460 | Vesicle-fusing ATPase | *Nsf* | -0.461776 | [illegible] |
| P46471;Q8BVQ9 | P46471;Q8BVQ9 | 26S protease regulatory subunit 7 | *Psmc2* | -0.285699 | [illegible] |
| P46664 | P46664 | Adenylosuccinate synthetase isozyme 2 | *Adss* | -0.252383 | [illegible] |
| P46737;A3KGA8;P46737-2 | P46737 | Lys-63-specific deubiquitinase BRCC36 | *Brcc3* | -0.176777 | [illegible] |
| P46935;Q3V335;V9GWV8;V9GXK3;A0A286YDV4;V9GXS6;A0A | P46935 | E3 ubiquitin-protein ligase NEDD4 | *Nedd4* | -0.373904 | [illegible] |
| P46938;G3UYW7;P46938-2;G3UYV4;G3UYA6;G3UY62 | P46938;G3UYW7;P46938-2;G3UYV4;G3UY | Transcriptional coactivator YAP1 | *Yap1* | -0.176777 | [illegible] |
| P47199;A0A0A6YXR4;V9GXY8;A0A0A6YVS7;D3YUG9;D3Z2X0 | P47199;A0A0A6YXR4;V9GXY8;A0A0A6YVS | Quinone oxidoreductase | *Cryz* | 1.69616 | [illegible] |
| Q5RKN9;P47753;A0A0G2JE27 | Q5RKN9;P47753 | F-actin-capping protein subunit alpha-1 | *Capza1* | -0.23007 | [illegible] |
| P47754;D6RCW7;A0A0N4SVM0 | P47754;D6RCW7 | F-actin-capping protein subunit alpha-2 | *Capza2* | 3.4321 | [illegible] |
| P47757-2;A2AMW0;P47757-4 | P47757-2;A2AMW0;P47757-4 | F-actin-capping protein subunit beta | *Capzb* | 3.69826 | [illegible] |
| P47791;P47791-2 | P47791;P47791-2 | Glutathione reductase, mitochondrial | *Gsr* | 0.970054 | [illegible] |
| P47856-2;P47856;D3YYD9;D3YYE0 | P47856-2;P47856 | Glutamine--fructose-6-phosphate aminotransferase [isome | *Gfpt1* | 4.41202 | [illegible] |
| P47857;P47857-2;P47857-3;A0A2R8VHV2;A0A2R8VHA5 | P47857;P47857-2;P47857-3 | ATP-dependent 6-phosphofructokinase, muscle type | *Pfkm* | 1.64545 | [illegible] |
| P47911;A0A0J9YU32 | P47911 | 60S ribosomal protein L6 | *Rpl6* | -0.215099 | [illegible] |
| P48036;A0A0G2JGQ0 | P48036 | Annexin A5 | *Anxa5* | -0.176777 | [illegible] |
| P48722;P48722-2;E0CY23;A0A0N4SVU2;A0A0N4SW76 | P48722;P48722-2;E0CY23 | Heat shock 70 kDa protein 4L | *Hspa4l* | 0.396459 | [illegible] |
| P48758;A0A338P684;A0A1B0GRG8 | P48758;A0A338P684 | Carbonyl reductase [NADPH] 1 | *Cbr1* | -0.292331 | [illegible] |
| P48759 | P48759 | Pentraxin-related protein PTX3 | *Ptx3* | 1.47307 | [illegible] |
| P49443;Q9EQE2;A0A1Y7VJZ4;A0A1Y7VNK7;A0A1Y7VL98 | P49443;Q9EQE2 | Protein phosphatase 1A | *Ppm1a* | 0.263464 | [illegible] |
| P50171;P50171-2;G3UX44 | P50171;P50171-2;G3UX44 | Estradiol 17-beta-dehydrogenase 8 | *Hsd17b8;H2-Ke6* | -0.176777 | [illegible] |
| P50228 | P50228 | C-X-C motif chemokine 5;GCP-2(1-78);GCP-2(9-78) | *Cxcl5* | -0.382423 | [illegible] |
| P50396;B7FAU8;D6RI86 | P50396 | Rab GDP dissociation inhibitor alpha | *Gdi1* | -0.422786 | [illegible] |
| P50427 | P50427 | Steryl-sulfatase | *Sts* | -0.176777 | [illegible] |

| | | | | |
|---|---|---|---|---|
| P50516;P50516-2;D3Z1B9;D3YWH3;D3YZ23 | P50516;P50516-2 | V-type proton ATPase catalytic subunit A | *Atp6v1a* | -0.168439 |
| P50518;A0A0N4SW07;A0A0N4SW34;A0A0N4SWA3;Q9D593 | P50518;A0A0N4SW07;A0A0N4SW34 | V-type proton ATPase subunit E 1 | *Atp6v1e1* | 1.1599 |
| P50543 | P50543 | Protein S100-A11 | *S100a11* | 1.46084 |
| P51174;A0A0R4J083 | P51174;A0A0R4J083 | Long-chain specific acyl-CoA dehydrogenase, mitochondria | *Acadl* | -0.176777 |
| P51410;A0A0G2JES3;A0A140T8T4;D3Z629;D3YZT0;A0A0G2JI | P51410;A0A0G2JES3;A0A140T8T4;D3Z62 | 60S ribosomal protein L9 | *Rpl9* | -0.176777 |
| P51432 | P51432 | 1-phosphatidylinositol 4,5-bisphosphate phosphodiesterase | *Plcb3* | -0.176777 |
| Q8BGZ6;P51569;A2BDV6 | Q8BGZ6;P51569 | Alpha-galactosidase A | *Gla* | 1.08797 |
| P51655 | P51655 | Glypican-4;Secreted glypican-4 | *Gpc4* | 0.220461 |
| P51660 | P51660 | Peroxisomal multifunctional enzyme type 2;(3R)-hydroxyac | *Hsd17b4* | -0.176777 |
| P51880;E9Q0H6 | P51880;E9Q0H6 | Fatty acid-binding protein, brain | *Fabp7* | -0.308316 |
| P51881 | P51881 | ADP/ATP translocase 2;ADP/ATP translocase 2, N-termina | *Slc25a5* | -0.202137 |
| P51910 | P51910 | Apolipoprotein D | *Apod* | -0.292977 |
| P52019;Q3TSS7 | P52019;Q3TSS7 | Squalene monooxygenase | *Sqle* | 0.713372 |
| P52293;A2A600;A2A601;A6PW68;F2Z431;A2A602 | P52293 | Importin subunit alpha-1 | *Kpna2* | -0.176777 |
| P52503;A0A1Y7VM38;A0A1Y7VKP8 | P52503;A0A1Y7VM38;A0A1Y7VKP8 | NADH dehydrogenase [ubiquinone] iron-sulfur protein 6, mi | *Ndufs6* | -0.176777 |
| P53810;J3QQ30;J3QPW1;F8WGG5 | P53810;J3QQ30;J3QPW1 | Phosphatidylinositol transfer protein alpha isoform | *Pitpna* | -0.379079 |
| P54071;D6RIL6;A0A0U1RP68;A0A0U1RPR1 | P54071;D6RIL6 | Isocitrate dehydrogenase [NADP], mitochondrial | *Idh2* | -0.385604 |
| P54116 | P54116 | Erythrocyte band 7 integral membrane protein | *Stom* | 1.22057 |
| P54775;A0A140LIZ5 | P54775;A0A140LIZ5 | 26S protease regulatory subunit 6B | *Psmc4* | -0.311398 |
| P54823 | P54823 | Probable ATP-dependent RNA helicase DDX6 | *Ddx6* | -0.176777 |
| P54923 | P54923 | [Protein ADP-ribosylarginine] hydrolase | *Adprh* | -0.197209 |
| P55065;A2A5K2;A2A5K3;A2A5K4 | P55065;A2A5K2;A2A5K3 | Phospholipid transfer protein | *Pltp* | -0.417008 |
| P55096;A0A0G2JDI9 | P55096;A0A0G2JDI9 | ATP-binding cassette sub-family D member 3 | *Abcd3* | -0.478235 |
| P55194;A2A5V3;A2A5V2;S4R2D3;A0A2R8W6I7;S4R2T6 | P55194;A2A5V3;A2A5V2;S4R2D3;A0A2R8V | SH3 domain-binding protein 1 | *Sh3bp1* | 3.65355 |
| P55258;D3YWL1;D3YW33;Q9CXS2;Q3TYH2;A0A286YDR2;A2A | P55258 | Ras-related protein Rab-8A | *Rab8a* | 4.49451 |
| P55264-2;P55264;A0A286YCD7;A0A286YCN3 | P55264-2;P55264;A0A286YCD7 | Adenosine kinase | *Adk* | -0.369617 |
| P55821 | P55821 | Stathmin-2 | *Stmn2* | -0.455066 |
| Q3U4W8;P56399;D3YYA5;D3Z4K7 | Q3U4W8;P56399 | Ubiquitin carboxyl-terminal hydrolase;Ubiquitin carboxyl-ter | *Usp5* | -0.339074 |
| P56480 | P56480 | ATP synthase subunit beta, mitochondrial | *Atp5b* | 1.55292 |
| Q91YZ2;P56546;P56546-2;E9PWQ9;E9Q0T4 | Q91YZ2;P56546;P56546-2;E9PWQ9 | C-terminal-binding protein 2 | *Ctbp2* | -0.252856 |
| P56564;O35544 | P56564 | Excitatory amino acid transporter 1 | *Slc1a3* | -0.238705 |
| P57759;F8WJI4;F8WIM7;D6RG87 | P57759 | Endoplasmic reticulum resident protein 29 | *Erp29* | 0.867788 |
| P58466 | P58466 | Carboxy-terminal domain RNA polymerase II polypeptide A | *Ctdsp1* | -0.39496 |
| P58871;Z4YJL4 | P58871;Z4YJL4 | 182 kDa tankyrase-1-binding protein | *Tnks1bp1* | -0.335162 |
| P59017 | P59017 | Bcl-2-like protein 13 | *Bcl2l13* | -0.176777 |
| P60122;D3YW60 | P60122 | RuvB-like 1 | *Ruvbl1* | -0.370938 |
| P60335;A0A087WR61 | P60335 | Poly(rC)-binding protein 1 | *Pcbp1* | -0.176777 |
| P60670-2;P60670 | P60670-2;P60670 | Nuclear protein localization protein 4 homolog | *Nploc4* | -0.382368 |
| P60766;P60766-1 | P60766;P60766-1 | Cell division control protein 42 homolog | *Cdc42* | -0.411256 |
| P60843 | P60843 | Eukaryotic initiation factor 4A-I | *Eif4a1* | -0.294952 |
| P61027 | P61027 | Ras-related protein Rab-10 | *Rab10* | -0.176777 |
| P61161 | P61161 | Actin-related protein 2 | *Actr2* | 1.59655 |
| P61164 | P61164 | Alpha-centractin | *Actr1a* | -0.560536 |
| P61202;P61202-2;A2AQE4 | P61202;P61202-2;A2AQE4 | COP9 signalosome complex subunit 2 | *Cops2* | -0.254076 |
| P61222 | P61222 | ATP-binding cassette sub-family E member 1 | *Abce1* | 2.01146 |
| P61358;A2A4Q0 | P61358 | 60S ribosomal protein L27 | *Rpl27* | 5.48008 |
| P61458;A0A1W2P7A4 | P61458;A0A1W2P7A4 | Pterin-4-alpha-carbinolamine dehydratase | *Pcbd1* | -0.625009 |
| P61514;A0A1D5RL86 | P61514 | 60S ribosomal protein L37a | *Rpl37a* | -0.369821 |
| P61793-2;P61793;A2AMJ0;A2AMI9 | P61793-2;P61793 | Lysophosphatidic acid receptor 1 | *Lpar1* | 1.82962 |
| P61922;P61922-2;F7C9G3;Q3TUE8;F6RN86 | P61922;P61922-2 | 4-aminobutyrate aminotransferase, mitochondrial | *Abat* | -0.176777 |
| P61971 | P61971 | Nuclear transport factor 2 | *Nutf2* | 2.59037 |
| P62075;A0A1W2P7H2;A0A1W2P756 | P62075;A0A1W2P7H2 | Mitochondrial import inner membrane translocase subunit T | *Timm13* | -0.176777 |
| P62137 | P62137 | Serine/threonine-protein phosphatase PP1-alpha catalytic s | *Ppp1ca* | 2.87571 |
| P62141;A0A0J9YUU8;A0A0J9YUG2 | P62141 | Serine/threonine-protein phosphatase PP1-beta catalytic su | *Ppp1cb* | -0.176777 |
| P62192 | P62192 | 26S protease regulatory subunit 4 | *Psmc1* | -0.357493 |
| P62196;Q8K1K2 | P62196;Q8K1K2 | 26S protease regulatory subunit 8 | *Psmc5* | -0.417808 |
| P62242 | P62242 | 40S ribosomal protein S8 | *Rps8* | -0.260791 |
| P62307 | P62307 | Small nuclear ribonucleoprotein F | *Snrpf* | 1.72925 |
| P62309 | P62309 | Small nuclear ribonucleoprotein G | *Snrpg* | -0.306093 |
| P62315 | P62315 | Small nuclear ribonucleoprotein Sm D1 | *Snrpd1* | -0.176777 |
| P62320;A0A1W2P7K5 | P62320;A0A1W2P7K5 | Small nuclear ribonucleoprotein Sm D3 | *Snrpd3* | 1.1853 |
| P62334;A0A2I3BQ79 | P62334 | 26S protease regulatory subunit 10B | *Psmc6* | -0.176777 |
| P62492;E9PZB2;E9Q6B3 | P62492 | Ras-related protein Rab-11A | *Rab11a* | -0.362261 |
| P62627;A2AVR9 | P62627;A2AVR9 | Dynein light chain roadblock-type 1 | *Dynlrb1* | -0.267066 |
| P62717;A0A1D5RLW5;A0A1D5RM85;A0A1D5RME4;A0A1D5RN | P62717;A0A1D5RLW5;A0A1D5RM85;A0A1I | 60S ribosomal protein L18a | *Rpl18a* | -0.176777 |
| P62737;P63268;D3YZY0;D3Z2K3;A0A0U1RQ96 | P62737;P63268;D3YZY0 | Actin, aortic smooth muscle;Actin, gamma-enteric smooth r | *Acta2;Actg2* | -0.242037 |
| P62751;D3YTY6 | P62751 | 60S ribosomal protein L23a | *Rpl23a* | 2.48424 |
| P62754 | P62754 | 40S ribosomal protein S6 | *Rps6* | -0.195714 |
| P62814;Q91YH6;A0A0U1RNU9 | P62814 | V-type proton ATPase subunit B, brain isoform | *Atp6v1b2* | 1.47455 |
| P62835;A0A0G2JE52;A0A0G2JDL9 | P62835;A0A0G2JE52;A0A0G2JDL9 | Ras-related protein Rap-1A | *Rap1a* | -0.279835 |
| P62849-2;P62849-3;P62849;A0A286YEB7 | P62849-2;P62849-3;P62849;A0A286YEB7 | 40S ribosomal protein S24 | *Rps24* | -0.176777 |
| P62855 | P62855 | 40S ribosomal protein S26 | *Rps26* | -0.176777 |
| P62874;H3BKR2;H3BLF7;Q61011 | P62874;H3BKR2;H3BLF7 | Guanine nucleotide-binding protein G(I)/G(S)/G(T) subunit | *Gnb1* | -0.354225 |
| P62880;E9QKR0;D3Z1M1;D3YZX3;D3Z1T4;E9PWM7;P29387;A | P62880;E9QKR0;D3Z1M1;D3YZX3;D3Z1T4 | Guanine nucleotide-binding protein G(I)/G(S)/G(T) subunit | *Gnb2* | -0.351478 |
| P62918 | P62918 | 60S ribosomal protein L8 | *Rpl8* | -0.274391 |
| P62965 | P62965 | Cellular retinoic acid-binding protein 1 | *Crabp1* | -0.176777 |
| P63005;P63005-2;Q5SW16 | P63005;P63005-2 | Platelet-activating factor acetylhydrolase IB subunit alpha | *Pafah1b1* | -0.0597865 |
| P63028;D3YU75 | P63028;D3YU75 | Translationally-controlled tumor protein | *Tpt1* | 0.00424666 |
| P63037;B1AXY1;B1AXY0;B1AXX9 | P63037;B1AXY1;B1AXY0 | DnaJ homolog subfamily A member 1 | *Dnaja1* | -0.215822 |
| P63038;P63038-2;D3Z2F2;D3Z7J9 | P63038 | 60 kDa heat shock protein, mitochondrial | *Hspd1* | -0.518986 |
| P63242;A0A0A0MQM0;Q8BGY2;J3QPS8 | P63242;A0A0A0MQM0;Q8BGY2 | Eukaryotic translation initiation factor 5A-1;Eukaryotic trans | *Eif5a;Eif5a2* | -0.228614 |
| P63248;A0A0A6YXS7 | P63248;A0A0A6YXS7 | cAMP-dependent protein kinase inhibitor alpha | *Pkia* | -0.176777 |
| P63325 | P63325 | 40S ribosomal protein S10 | *Rps10* | -0.190338 |
| P63328-2;P63328 | P63328-2;P63328 | Serine/threonine-protein phosphatase 2B catalytic subunit a | *Ppp3ca* | -0.253089 |
| P63330 | P63330 | Serine/threonine-protein phosphatase 2A catalytic subunit a | *Ppp2ca* | -0.540972 |
| P67984 | P67984 | 60S ribosomal protein L22 | *Rpl22* | -0.253991 |
| P68254-2;P68254;F6YY69;F6VW30 | P68254-2;P68254;F6YY69;F6VW30 | 14-3-3 protein theta | *Ywhaq* | -0.607633 |
| P68369;P05214 | P68369;P05214 | Tubulin alpha-1A chain;Tubulin alpha-3 chain | *Tuba1a;Tuba3a* | -0.328024 |
| P68372;G3UZR1 | P68372 | Tubulin beta-4B chain | *Tubb4b* | -0.240407 |
| P68510 | P68510 | 14-3-3 protein eta | *Ywhah* | -0.36867 |
| P70122;F6TN03;D6REV5 | P70122;F6TN03 | Ribosome maturation protein SBDS | *Sbds* | 4.32997 |
| P70168 | P70168 | Importin subunit beta-1 | *Kpnb1* | -0.43244 |
| P70195 | P70195 | Proteasome subunit beta type-7 | *Psmb7* | -0.4459 |
| P70202 | P70202 | Latexin | *Lxn* | -0.29251 |
| Q3UGN9;P70297;A2ARJ0;A0A0A6YVP6 | Q3UGN9;P70297;A2ARJ0 | Signal transducing adapter molecule 1 | *Stam* | -0.312546 |
| P70372 | P70372 | ELAV-like protein 1 | *Elavl1* | -0.176777 |
| Q4FE56;P70398;E9PWA9;F8VPU6;G3UZS3 | Q4FE56;P70398 | Ubiquitin carboxyl-terminal hydrolase;Probable ubiquitin car | *Usp9x* | 2.93636 |
| P70441;P70441-2 | P70441 | Na(+)/H(+) exchange regulatory cofactor NHE-RF1 | *Slc9a3r1* | -0.238434 |
| P70697;A0A0A0MQG7 | P70697 | Uroporphyrinogen decarboxylase | *Urod* | -0.284397 |
| P70698 | P70698 | CTP synthase 1 | *Ctps1* | -0.330711 |
| P70699;F6R5R5;A2AFL5;F6VEG4;A2AFL3 | P70699 | Lysosomal alpha-glucosidase | *Gaa* | -0.531553 |
| P80314;A0A1W2P7B7;A0A1W2P828;A0A1W2P6Q3;A0A1W2P8 | P80314;A0A1W2P7B7 | T-complex protein 1 subunit beta | *Cct2* | -0.799022 |
| P80315;G5E839;G3UYW5;G3UXF3;G3UXG2 | P80315;G5E839 | T-complex protein 1 subunit delta | *Cct4* | -0.212389 |
| P80317;Q61390;B1AT05 | P80317 | T-complex protein 1 subunit zeta | *Cct6a* | -0.20887 |
| P82198;Q3UXJ2 | P82198;Q3UXJ2 | Transforming growth factor-beta-induced protein ig-h3 | *Tgfbi* | -0.500535 |
| P83741-4;P83741-5;P83741;P83741-2;P83741-3;P83741-6;F7C | P83741-4;P83741-5;P83741;P83741-2;P837 | Serine/threonine-protein kinase WNK1 | *Wnk1* | -0.320149 |
| P84084 | P84084 | ADP-ribosylation factor 5 | *Arf5* | -0.193449 |
| P84089;G3UW85;A0A1W2P7H9;A0A1W2P7T3 | P84089;G3UW85;A0A1W2P7H9;A0A1W2P7 | Enhancer of rudimentary homolog | *Erh* | -0.291997 |
| P97290;A2ATR8 | P97290;A2ATR8 | Plasma protease C1 inhibitor | *Serping1* | 2.87828 |
| P97300-1;P97300;H3BIX4;Z4YLB7;P97300-3;A0A0A0MQN8;H3I | P97300-1;P97300;H3BIX4;Z4YLB7;P97300- | Neuroplastin | *Nptn* | -0.176777 |
| P97310;A0A0N4SVY2 | P97310 | DNA replication licensing factor MCM2 | *Mcm2* | 1.16193 |
| P97315 | P97315 | Cysteine and glycine-rich protein 1 | *Csrp1* | -0.176777 |
| P97326 | P97326 | Cadherin-6 | *Cdh6* | -0.176777 |
| P97346;P97346-2 | P97346;P97346-2 | Nucleoredoxin | *Nxn* | -0.340616 |
| P97351 | P97351 | 40S ribosomal protein S3a | *Rps3a* | 0.771773 |
| P97372;G3X9V0;E0CZ90 | P97372;G3X9V0 | Proteasome activator complex subunit 2 | *Psme2* | -0.368776 |
| P97384;D3Z7U0 | P97384;D3Z7U0 | Annexin A11;Annexin | *Anxa11* | -0.300366 |
| P97390 | P97390 | Vacuolar protein sorting-associated protein 45 | *Vps45* | -0.176777 |
| P97429;A0A0N4SW89;D3Z0S1;A0A0N4SV57;F7ANV6;S4R1F2 | P97429;A0A0N4SW89;D3Z0S1;A0A0N4SV | Annexin A4;Annexin | *Anxa4* | -0.176777 |
| P97434;P97434-2;P97434-5;F6RND9;P97434-4;P97434-3;F6S | P97434;P97434-2;P97434-5;F6RND9;P974 | Myosin phosphatase Rho-interacting protein | *Mprip* | -0.35972 |
| P97447;A2AEX8;A2AEX6;Q8CDC8 | P97447;A2AEX8;A2AEX6 | Four and a half LIM domains protein 1 | *Fhl1* | -1.07254 |
| P97450;E9QAD6 | P97450;E9QAD6 | ATP synthase-coupling factor 6, mitochondrial | *Atp5j* | -0.253255 |
| P97494 | P97494 | Glutamate--cysteine ligase catalytic subunit | *Gclc* | -0.176777 |
| P97797-2;P97797;Q6P6I8;E0CYM8;P97797-3;A0A0R4J1Z7;E9 | P97797-2;P97797;Q6P6I8;E0CYM8 | Tyrosine-protein phosphatase non-receptor type substrate | *Sirpa* | -0.246866 |
| P97807-2;P97807;H3BKG7 | P97807-2;P97807 | Fumarate hydratase, mitochondrial | *Fh* | 0.70413 |
| P97855 | P97855 | Ras GTPase-activating protein-binding protein 1 | *G3bp1* | -0.322199 |
| P98192;Q05DV2 | P98192;Q05DV2 | Dihydroxyacetone phosphate acyltransferase | *Gnpat* | -0.176777 |
| P98203-4;P98203-5;P98203-2;P98203;A0A338P7H0 | P98203-4;P98203-5;P98203-2;P98203 | Armadillo repeat protein deleted in velo-cardio-facial syndro | *Arvcf* | -0.395392 |
| P99024 | P99024 | Tubulin beta-5 chain | *Tubb5* | -0.355602 |
| P99026 | P99026 | Proteasome subunit beta type-4 | *Psmb4* | -0.303912 |
| P99028 | P99028 | Cytochrome b-c1 complex subunit 6, mitochondrial | *Uqcrh* | -0.237752 |
| P99029-2;P99029;G3UZJ4;H3BJQ7 | P99029-2;P99029;G3UZJ4;H3BJQ7 | Peroxiredoxin-5, mitochondrial | *Prdx5* | -0.643656 |
| Q01705-4;Q01705-3;Q01705;Q01705-2 | Q01705-4;Q01705-3;Q01705;Q01705-2 | Neurogenic locus notch homolog protein 1;Notch 1 extracel | *Notch1* | -0.370758 |
| Q9D031;Q01730;A2AUR7;B1AYQ0;E0CXG5;A0A0A6YWZ2 | Q9D031;Q01730;A2AUR7 | Ras suppressor protein 1 | *Rsu1* | -0.176777 |
| Q02053;P31254 | Q02053 | Ubiquitin-like modifier-activating enzyme 1 | *Uba1* | -0.457851 |
| Q02248;E9Q6A9;F7BAC9;D3YUH4;F6QZ47;F7CRC6;D3Z5Q1; | Q02248;E9Q6A9 | Catenin beta-1 | *Ctnnb1* | -0.59453 |
| Q03146-2;Q03146;G3UXE5;D6RII6;B0V2I0;B0V2H9;E9PW79;B | Q03146-2;Q03146;G3UXE5 | Epithelial discoidin domain-containing receptor 1 | *Ddr1* | -0.247094 |
| Q03265;D3Z6F5;D6RJ16 | Q03265;D3Z6F5 | ATP synthase subunit alpha, mitochondrial;ATP synthase s | *Atp5a1* | -0.68384 |
| Q03350 | Q03350 | Thrombospondin-2 | *Thbs2* | -0.176777 |
| Q03366 | Q03366 | C-C motif chemokine 7 | *Ccl7* | -0.30233 |
| Q03963 | Q03963 | Interferon-induced, double-stranded RNA-activated protein | *Eif2ak2* | -0.250336 |
| Q04447;A0A0J9YKD4;P07310 | Q04447 | Creatine kinase B-type | *Ckb* | -0.412984 |
| Q04690-2;Q04690;A0A1W2P863;Q04690-4;Q04690-3;Q5SYH9 | Q04690-2;Q04690;A0A1W2P863;Q04690-4 | Neurofibromin | *Nf1* | 2.47259 |
| Q04735-2;Q04735;E3W997;Q99J95-3;Q80YP0;Q99J95-2;O354 | Q04735-2;Q04735 | Cyclin-dependent kinase 16 | *Cdk16* | -0.253861 |
| Q05816 | Q05816 | Fatty acid-binding protein, epidermal | *Fabp5* | -0.176777 |
| Q08024-2;Q08024-4;Q08024-3;Q08024 | Q08024-2 | Core-binding factor subunit beta | *Cbfb* | -0.254062 |
| Q08093;D3Z7R6 | Q08093;D3Z7R6 | Calponin-2;Calponin | *Cnn2* | -0.176777 |
| Q0VBD0 | Q0VBD0 | Integrin beta | *Itgb8* | -0.176777 |
| Q11011;E9Q039;F6QYF8;E9Q6F4;F6V7K3;F7ANF4;F2Z3V5 | Q11011;E9Q039;F6QYF8;E9Q6F4 | Puromycin-sensitive aminopeptidase | *Npepps* | 0.47865 |
| Q3B7Z2;Q3B7Z2-2;Q5F209;Q5QNQ6 | Q3B7Z2;Q3B7Z2-2 | Oxysterol-binding protein 1 | *Osbp* | -0.650166 |
| Q3TCN2;Q3TCN2-2 | Q3TCN2;Q3TCN2-2 | Putative phospholipase B-like 2;Putative phospholipase B-li | *Plbd2* | 4.43948 |
| Q3TDD9;Q3TDD9-2 | Q3TDD9 | Protein phosphatase 1 regulatory subunit 21 | *Ppp1r21* | -0.250708 |
| Z4YKB8;Q3TEA8-2;Q3TEA8;Z4YKA3;Q3TEA8-3;A2AM65;A2AM | Z4YKB8;Q3TEA8-2;Q3TEA8;Z4YKA3;Q3TE | Heterochromatin protein 1-binding protein 3 | *Hp1bp3* | -0.176777 |
| Q3TGM7;Q69ZS7;Q69ZS7-2;J3KMK0;A0A1W2P833;L7N209;Q | Q3TGM7;Q69ZS7;Q69ZS7-2 | HBS1-like protein | *Hbs1l* | -0.314035 |
| Q6ZWQ5;Q3TGS7;Q3V2H3;O70493 | Q6ZWQ5;Q3TGS7;Q3V2H3;O70493 | Sorting nexin-12 | *Snx12* | 1.98209 |
| Q3TIR3;A0A1B0GSI8;A0A1B0GS30;Q80XE1;Q80XE1-1 | Q3TIR3;A0A1B0GSI8 | Synembryn-A | *Ric8a* | -0.176777 |
| Q3TIV5;Q3TIV5-2;Q3TIV5-3 | Q3TIV5;Q3TIV5-2 | Zinc finger CCCH domain-containing protein 15 | *Zc3h15* | -0.682902 |
| Q3TIX9 | Q3TIX9 | U4/U6.U5 tri-snRNP-associated protein 2 | *Usp39* | -0.176777 |
| Q3TJD7;Q8BVJ7;Q3TJD7-2;B8JJB3;Q3TJD7-3;B8JJB2;B8JJE | Q3TJD7;Q8BVJ7 | PDZ and LIM domain protein 7 | *Pdlim7* | -0.757138 |
| Q3TKX1;Q9R1Q9;B7FAU3;B7FAU7 | Q3TKX1;Q9R1Q9;B7FAU3 | V-type proton ATPase subunit S1 | *Atp6ap1* | -0.280544 |
| Q80ZP8;Q3TMX5;Q9CXI5;F6USD5;F6T4L3;F7C1S6 | Q80ZP8;Q3TMX5;Q9CXI5 | Mesencephalic astrocyte-derived neurotrophic factor | *Manf* | -0.365398 |
| Q3TVI8;D3YUE1;D3YY15;D3Z2F0 | Q3TVI8 | Pre-B-cell leukemia transcription factor-interacting protein 1 | *Pbxip1* | -0.246767 |
| Q3TVK3;Q9Z2W0;Q8BPW9;A0A087WS31;A0A087WSE6;A0A08 | Q3TVK3;Q9Z2W0;Q8BPW9;A0A087WS31; | Aspartyl aminopeptidase | *Dnpep* | -0.254129 |
| Q3TW96;Q3TW96-2 | Q3TW96;Q3TW96-2 | UDP-N-acetylhexosamine pyrophosphorylase-like protein | *Uap1l1* | -0.253839 |
| Q3TWI9;D3YY83;D3Z2D8;D3YXV7;D3YYJ8 | Q3TWI9;D3YY83;D3Z2D8 | CSC1-like protein 2 | *Tmem63b* | -0.176777 |
| Q3TXV4;Q921E2;Q3TM43;D6RFT6 | Q3TXV4;Q921E2 | Ras-related protein Rab-31 | *Rab31* | -0.419537 |
| Q3U1U4;E9Q604;G5E8F1;E9Q5K8;P05555;A0A0R4J1B4;P055 | Q3U1U4;E9Q604;G5E8F1;E9Q5K8;P05555 | Integrin alpha-M | *Itgam* | -0.176777 |
| Q3U2G2;Q61316 | Q3U2G2;Q61316 | Heat shock 70 kDa protein 4 | *Hspa4* | -0.67523 |
| Q3U367;Q9JLJ2 | Q3U367;Q9JLJ2 | 4-trimethylaminobutyraldehyde dehydrogenase | *Aldh9a1* | -0.56476 |
| Q3U3G8;Q9EPB5;F6TMN2 | Q3U3G8;Q9EPB5 | Serine hydrolase-like protein | *Serhl* | -0.176777 |
| Q3U3J1;P50136;A0A0U1RPK5 | Q3U3J1;P50136 | 2-oxoisovalerate dehydrogenase subunit alpha, mitochondr | *Bckdha* | 3.59357 |
| Q3U422;Q8BK30 | Q3U422 | | *Ndufv3* | -0.176777 |
| Q3U741;Q501J6;Q3TU25;Q501J6-2 | Q3U741;Q501J6;Q3TU25;Q501J6-2 | Probable ATP-dependent RNA helicase DDX17 | *Ddx17* | -0.482291 |
| Q8C0M2;Q3U7M5;Q8VD66;A0A0G2JEC8;A0A0G2JDM2;A0A0 | Q8C0M2;Q3U7M5;Q8VD66;A0A0G2JEC8; | Abhydrolase domain-containing protein 4 | *Abhd4* | -0.176777 |
| Q3U9G9;A0A0A6YY12 | Q3U9G9;A0A0A6YY12 | Lamin-B receptor | *Lbr* | -0.313906 |
| Q3U9H3;Q61337;F7ABX5;D3YZR8;F6S493 | Q3U9H3;Q61337;F7ABX5;D3YZR8 | Bcl2-associated agonist of cell death | *Bad* | -0.315268 |
| Q3UCV8;A0A2I3BRS5 | Q3UCV8;A0A2I3BRS5 | Ubiquitin thioesterase otulin | *Otulin* | -0.176777 |
| Q3UE92;Q6P1B1;S4R1I3;S4R228;E9Q286 | Q3UE92;Q6P1B1;S4R1I3 | Xaa-Pro aminopeptidase 1 | *Xpnpep1* | 0.799473 |
| Q3UEB3-3;Q3UEB3-2;Q3UEB3 | Q3UEB3-3;Q3UEB3-2;Q3UEB3 | Poly(U)-binding-splicing factor PUF60 | *Puf60* | -0.311602 |
| Q3UGB5;Q9JII5-2;Q9JII5;D3Z4J1 | Q3UGB5;Q9JII5-2;Q9JII5;D3Z4J1 | DAZ-associated protein 1 | *Dazap1* | 1.05235 |
| Q3UH60;B2RQC7;Q3UH60-2 | Q3UH60;B2RQC7;Q3UH60-2 | Disco-interacting protein 2 homolog B | *Dip2b* | 0.447894 |
| Q3UH68;D3YU22;D3Z589;Q3UH68-2;D3YU59;Q3UH68-3;F6X | Q3UH68;D3YU22;D3Z589;Q3UH68-2;D3YU | LIM and calponin homology domains-containing protein 1 | *Limch1* | -0.451858 |
| Q3UHA3 | Q3UHA3 | Spatacsin | *Spg11* | -0.319854 |
| Q3UHD6-2;Q3UHD6;A0A0G2JF85;A0A0G2JG07;A0A0G2JGD | Q3UHD6-2;Q3UHD6;A0A0G2JF85 | Sorting nexin-27 | *Snx27* | -0.207145 |
| Q3UHG5;Q62283 | Q3UHG5;Q62283 | Tetraspanin;Tetraspanin-7 | *Tspan7* | -0.236202 |
| Q3UHJ0-2;Q3UHJ0;A0A0N4SVA1 | Q3UHJ0-2;Q3UHJ0 | AP2-associated protein kinase 1 | *Aak1* | 0.385389 |
| Q3UID0;Q6PDG5-2;Q6PDG5;Q3UNN4;P97496-2;P97496;A0A1 | Q3UID0;Q6PDG5-2;Q6PDG5 | SWI/SNF complex subunit SMARCC2 | *Smarcc2* | -0.298694 |
| Q3UJB0 | Q3UJB0 | | *Sf3b2* | -0.176777 |
| Q3ULD5;Q6PD20 | Q3ULD5;Q6PD20 | Methylcrotonoyl-CoA carboxylase beta chain, mitochondria | *Mccc2* | -0.176777 |
| Q3ULJ0-2;Q3ULJ0;D3Z0L6 | Q3ULJ0-2;Q3ULJ0 | Glycerol-3-phosphate dehydrogenase 1-like protein | *Gpd1l* | -0.227971 |
| Q3UM45;A0A087WRA7;F6TGJ2 | Q3UM45 | Protein phosphatase 1 regulatory subunit 7 | *Ppp1r7* | 0.534744 |

| | | | | | |
|---|---|---|---|---|---|
| Q3UMB9;A0A1W2P6V9 | Q3UMB9 | WASH complex subunit 7 | *Kiaa1033* | -0.312547 | [illegible] |
| Q3UMF0-3;Q3UMF0;B1AZ15;B1AZ14;Q3UMF0-4;Q3UMF0-2;E | Q3UMF0-3;Q3UMF0;B1AZ15;B1AZ14;Q3U | Cordon-bleu protein-like 1 | *Cobll1* | -0.30941 | [illegible] |
| Q3UN04;A0A0G2JDF7 | Q3UN04;A0A0G2JDF7 | Ubiquitin carboxyl-terminal hydrolase 30 | *Usp30* | -0.253026 | [illegible] |
| S4R189;Q3UPH7-3;Q3UPH7-6;Q3UPH7-2;Q3UPH7;S4R2U9 | S4R189;Q3UPH7-3;Q3UPH7-6;Q3UPH7-2; | Rho guanine nucleotide exchange factor 40 | *Arhgef40* | -0.308321 | [illegible] |
| Q3USX5;Q8R4K2 | Q3USX5;Q8R4K2 | Interleukin-1 receptor-associated kinase 4 | *Irak4* | -0.250166 | [illegible] |
| Q3UVG3 | Q3UVG3 | Protein FAM91A1 | *Fam91a1* | -0.410234 | [illegible] |
| Q3UVK0;F6TQS9;Q3UVK0-2;G8JL46 | Q3UVK0 | Endoplasmic reticulum metallopeptidase 1 | *Ermp1* | -0.25244 | [illegible] |
| Q3UYV9 | Q3UYV9 | Nuclear cap-binding protein subunit 1 | *Ncbp1* | -0.176777 | [illegible] |
| Q3V0K9 | Q3V0K9 | Plastin-1 | *Pls1* | -0.252806 | [illegible] |
| Q3V117;Q91V92;Q3TS02 | Q3V117;Q91V92 | ATP-citrate synthase | *Acly* | 1.22856 | [illegible] |
| Q3V1H1 | Q3V1H1 | Cytoskeleton-associated protein 2 | *Ckap2* | -0.176777 | [illegible] |
| Q3V3R1 | Q3V3R1 | Monofunctional C1-tetrahydrofolate synthase, mitochondria | *Mthfd1l* | -0.176777 | [illegible] |
| Q3V3R4 | Q3V3R4 | Integrin alpha-1 | *Itga1* | -0.58003 | [illegible] |
| Q561M1;Q9D358-2 | Q561M1;Q9D358-2 | Low molecular weight phosphotyrosine protein phosphatas | *Acp1* | -0.215584 | [illegible] |
| Q569Z5;F8WHR6;Q569Z5-2 | Q569Z5;F8WHR6;Q569Z5-2 | Probable ATP-dependent RNA helicase DDX46 | *Ddx46* | -0.370355 | [illegible] |
| Q569Z6;Q8BZN7;F7AB07;F6YSQ2 | Q569Z6;Q8BZN7 | Thyroid hormone receptor-associated protein 3 | *Thrap3* | -0.254128 | [illegible] |
| Q58A65-2;Q58A65;Q58A65-6;Q58A65-5;A0A0R4J196;F8WH31 | Q58A65-2;Q58A65;Q58A65-6;Q58A65-5;A( | C-Jun-amino-terminal kinase-interacting protein 4 | *Spag9* | -0.31411 | [illegible] |
| Q5F2E8;Q6ZQ29-2;Q6ZQ29 | Q5F2E8 | Serine/threonine-protein kinase TAO1 | *Taok1* | -0.663338 | [illegible] |
| Q5M8N4;A0A2I3BQS7 | Q5M8N4;A0A2I3BQS7 | Epimerase family protein SDR39U1 | *Sdr39u1* | -0.472431 | [illegible] |
| Q5PRF0;A0A1W2P7I6 | Q5PRF0 | HEAT repeat-containing protein 5A | *Heatr5a* | -0.176777 | [illegible] |
| Q5SNZ0-2;Q5SNZ0;Z4YLP6;Q5SNZ0-3;F6R1B7 | Q5SNZ0-2;Q5SNZ0;Z4YLP6;Q5SNZ0-3 | Girdin | *Ccdc88a* | 3.50086 | [illegible] |
| Q5SRX1;Q5SRX1-2;Q5SXA5;Q5SRX1-3;Q5SRX1-4;Q5SXA4;Q | Q5SRX1;Q5SRX1-2;Q5SXA5;Q5SRX1-3;Q | TOM1-like protein 2 | *Tom1l2* | -0.253951 | [illegible] |
| Q5SUR0;F6RUL9;G3UY86 | Q5SUR0 | Phosphoribosylformylglycinamidine synthase | *Pfas* | -0.313062 | [illegible] |
| Q5SUT0;Q5SUS9;Q61545;Q5SUS8 | Q5SUT0;Q5SUS9;Q61545 | RNA-binding protein EWS | *Ewsr1* | -0.22889 | [illegible] |
| Q5SW88;P62821;Q5SW87;Q5SW86 | Q5SW88;P62821;Q5SW87 | Ras-related protein Rab-1A | *Rab1;Rab1A* | -0.601528 | [illegible] |
| Q5SWZ5;F6XZM9 | Q5SWZ5 | | *Mprip* | -0.52619 | [illegible] |
| Q5SXR6;Q68FD5 | Q5SXR6;Q68FD5 | Clathrin heavy chain;Clathrin heavy chain 1 | *Cltc* | -0.425336 | [illegible] |
| Q5SYL3 | Q5SYL3 | Protein KIAA0100 | *Kiaa0100* | -0.252804 | [illegible] |
| Q5U458;E9Q8B3 | Q5U458;E9Q8B3 | DnaJ homolog subfamily C member 11 | *Dnajc11* | -0.176777 | [illegible] |
| Q5XJY5 | Q5XJY5 | Coatomer subunit delta | *Arcn1* | -0.308823 | [illegible] |
| Q60597;Z4YJV4;Q60597-2;Q60597-4;Q60597-3 | Q60597;Z4YJV4;Q60597-2;Q60597-4;Q60 | 2-oxoglutarate dehydrogenase, mitochondrial | *Ogdh* | -0.176777 | [illegible] |
| Q60605;A0A1W2P6F6;A0A1W2P7Q9;Q60605-2;A0A1W2P6G5 | Q60605;A0A1W2P6F6;A0A1W2P7Q9;Q606 | Myosin light polypeptide 6 | *Myl6* | -0.313734 | [illegible] |
| Q60648 | Q60648 | Ganglioside GM2 activator | *Gm2a* | -0.446745 | [illegible] |
| Q60692 | Q60692 | Proteasome subunit beta type-6 | *Psmb6* | -0.386789 | [illegible] |
| Q60715-2;E9Q7B0 | Q60715-2;E9Q7B0 | Prolyl 4-hydroxylase subunit alpha-1 | *P4ha1* | -0.382583 | [illegible] |
| Q60749;Q9WU01-2;Q9WU01 | Q60749 | KH domain-containing, RNA-binding, signal transduction-as | *Khdrbs1* | -0.326556 | [illegible] |
| Q60770;Q60770-2;A0A0G2JEI5;A2AEC9;A0A0G2JEQ3 | Q60770;Q60770-2 | Syntaxin-binding protein 3 | *Stxbp3* | -0.357523 | [illegible] |
| Q60841-3;Q60841-2;Q60841 | Q60841-3;Q60841-2;Q60841 | Reelin | *Reln* | -0.649696 | [illegible] |
| Q60854;F8WIV2;K7E6F1;E9Q3Y1;E9Q6X2;E9Q108;E9Q0P9;E | Q60854;F8WIV2;K7E6F1 | Serpin B6 | *Serpinb6;Serpinb6a* | -0.347245 | [illegible] |
| Q60865;F6YP03;A2BH47;F6YLI0 | Q60865 | Caprin-1 | *Caprin1* | -0.176777 | [illegible] |
| Q60866;A2AUR3;A2AUR5;Q60866-2 | Q60866;A2AUR3;A2AUR5;Q60866-2 | Phosphotriesterase-related protein | *Pter* | -0.176777 | [illegible] |
| Q60902-3;Q60902;A0A1D5RLS1;Q60902-4;Q60902-2 | Q60902-3;Q60902;A0A1D5RLS1;Q60902-4 | Epidermal growth factor receptor substrate 15-like 1 | *Eps15l1* | -0.359102 | [illegible] |
| Q60931;J3QMG3 | Q60931;J3QMG3 | Voltage-dependent anion-selective channel protein 3 | *Vdac3* | -0.176777 | [illegible] |
| Q60932-2;Q60932;F2Z471 | Q60932-2;Q60932;F2Z471 | Voltage-dependent anion-selective channel protein 1 | *Vdac1* | -0.468802 | [illegible] |
| Q60953-2;Q60953;F7BTZ2;D3Z3A6;D3YXR5 | Q60953-2;Q60953;F7BTZ2;D3Z3A6;D3YXF | Protein PML | *Pml* | -0.176777 | [illegible] |
| Q6NZM8;Q60967;A0A0G2JFM1;A0A0G2JFK8 | Q6NZM8;Q60967 | Bifunctional 3-phosphoadenosine 5-phosphosulfate synthas | *Papss1* | -0.176777 | [illegible] |
| Q60972 | Q60972 | Histone-binding protein RBBP4 | *Rbbp4* | -0.420381 | [illegible] |
| Q61001 | Q61001 | Laminin subunit alpha-5 | *Lama5* | -0.176777 | [illegible] |
| Q61024;D3Z0C2;D3Z028;D3YYR0 | Q61024 | Asparagine synthetase [glutamine-hydrolyzing] | *Asns* | -0.445993 | [illegible] |
| Q61029;Q61029-2;Q61029-4;Q61033-2 | Q61029;Q61029-2;Q61029-4 | Lamina-associated polypeptide 2, isoforms beta/delta/epsilc | *Tmpo* | 0.171139 | [illegible] |
| Q61033 | Q61033 | Lamina-associated polypeptide 2, isoforms alpha/zeta | *Tmpo* | -0.253506 | [illegible] |
| Q61081;A0A1L1STC0 | Q61081;A0A1L1STC0 | Hsp90 co-chaperone Cdc37;Hsp90 co-chaperone Cdc37, | *Cdc37* | -0.289456 | [illegible] |
| Q61112;Q61112-2 | Q61112;Q61112-2 | 45 kDa calcium-binding protein | *Sdf4* | -0.473966 | [illegible] |
| Q61166 | Q61166 | Microtubule-associated protein RP/EB family member 1 | *Mapre1* | -0.357145 | [illegible] |
| Q61171;D3Z4A4 | Q61171;D3Z4A4 | Peroxiredoxin-2 | *Prdx2* | -0.746775 | [illegible] |
| Q61216;Q61216-2;F6RX99 | Q61216;Q61216-2;F6RX99 | Double-strand break repair protein MRE11A | *Mre11a* | -0.24763 | [illegible] |
| Q61233;D3YZ25;D3YVW8;D3Z7D9;D3Z311 | Q61233 | Plastin-2 | *Lcp1* | 0.827118 | [illegible] |
| Q61245-2;Q61245;E9PVT8;V9GWX8 | Q61245-2;Q61245;E9PVT8 | Collagen alpha-1(XI) chain | *Col11a1* | -0.241438 | [illegible] |
| Q61391;A0A0A6YY36;A0A0A6YWB9;A0A0A6YW93 | Q61391 | Neprilysin | *Mme* | -0.312893 | [illegible] |
| Q61425 | Q61425 | Hydroxyacyl-coenzyme A dehydrogenase, mitochondrial | *Hadh* | -0.315882 | [illegible] |
| Q61474-2;Q61474;A0A0J9YU67;F8WJA5;A0A0J9YTX9 | Q61474-2;Q61474;A0A0J9YU67;F8WJA5;A | RNA-binding protein Musashi homolog 1 | *Msi1* | -0.176777 | [illegible] |
| Q8BRR9;Q61481;F6X0V1;Q61481-2;Q9JLL9;Q5I7S8;A2ASF9;( | Q8BRR9;Q61481;F6X0V1;Q61481-2;Q9JLL | Calcium/calmodulin-dependent 3,5-cyclic nucleotide phospl | *Pde1a* | 4.69045 | [illegible] |
| Q61553;A0A0G2JDU7;F7BDR1 | Q61553;A0A0G2JDU7 | Fascin | *Fscn1* | 0.0424435 | [illegible] |
| Q61576;A2A4H9;F6W360 | Q61576;A2A4H9 | Peptidyl-prolyl cis-trans isomerase FKBP10 | *Fkbp10* | -0.176777 | [illegible] |
| Q61578 | Q61578 | NADPH:adrenodoxin oxidoreductase, mitochondrial | *Fdxr* | -0.253946 | [illegible] |
| Q61584-5;Q61584-6;Q61584-4;Q61584-7;Q61584-3;Q61584;A( | Q61584-5;Q61584-6;Q61584-4;Q61584-7;( | Fragile X mental retardation syndrome-related protein 1 | *Fxr1* | -0.361506 | [illegible] |
| Q61592 | Q61592 | Growth arrest-specific protein 6 | *Gas6* | -0.252491 | [illegible] |
| Q61598;Q61598-2;A0A1Y7VL99;A0A1Y7VLG4 | Q61598;Q61598-2;A0A1Y7VL99 | Rab GDP dissociation inhibitor beta | *Gdi2* | -0.584291 | [illegible] |
| Q61655 | Q61655 | ATP-dependent RNA helicase DDX19A | *Ddx19a* | -0.25865 | [illegible] |
| Q8BTS0;Q61656;S4R1I6;B1ARB9;B1ARC0;S4R1E3 | Q8BTS0;Q61656;S4R1I6 | Probable ATP-dependent RNA helicase DDX5 | *Ddx5* | 2.38232 | [illegible] |
| Q61699-2;Q61699;E9Q0U7;D3Z027;D3Z3I9;A0A0J9YTZ7 | Q61699-2;Q61699;E9Q0U7 | Heat shock protein 105 kDa | *Hsph1* | -0.338093 | [illegible] |
| Q61739-2;Q61739;F6VSK8 | Q61739-2;Q61739;F6VSK8 | Integrin alpha-6;Integrin alpha-6 heavy chain;Integrin alpha- | *Itga6* | -0.31392 | [illegible] |
| Q61753;F6ZSB7 | Q61753 | D-3-phosphoglycerate dehydrogenase | *Phgdh* | -0.646046 | [illegible] |
| Q61792;A2A6H0;A2A6G9;A2A6G7;A2A6G8;A2A6G6;A2A6H1;A | Q61792;A2A6H0;A2A6G9;A2A6G7;A2A6G8 | LIM and SH3 domain protein 1 | *Lasp1* | -0.406537 | [illegible] |
| Q61937;Q9DAY9;Q5SQB5;Q5SQB0 | Q61937;Q9DAY9;Q5SQB5;Q5SQB0 | Nucleophosmin | *Npm1* | -0.38137 | [illegible] |
| Q923A8;Q62073;A2AP93;A2AP92 | Q923A8;Q62073;A2AP93;A2AP92 | Mitogen-activated protein kinase kinase kinase 7 | *Map3k7* | -0.176777 | [illegible] |
| Q62095 | Q62095 | ATP-dependent RNA helicase DDX3Y | *Ddx3y* | -0.311269 | [illegible] |
| Q62101;D3Z6I0;Q5FWX6;Q8K1Y2;Q8BZ03;B1AU42;F2Z3Z6;B | Q62101;D3Z6I0;Q5FWX6;Q8K1Y2;Q8BZ03 | Serine/threonine-protein kinase D1;Protein kinase C;Serine | *Prkd1;Prkd3;Prkd2* | 0.917898 | [illegible] |
| Q62181;E9Q0Z0 | Q62181 | Semaphorin-3C | *Sema3c* | -0.176777 | [illegible] |
| Q9D8L3;Q62186 | Q9D8L3;Q62186 | Translocon-associated protein subunit delta | *Ssr4* | -0.350145 | [illegible] |
| Q62261;A0A0A0MQG2;Q62261-2;Q68FG2 | Q62261;A0A0A0MQG2;Q62261-2 | Spectrin beta chain, non-erythrocytic 1 | *Sptbn1* | 2.3498 | [illegible] |
| Q62318;Q62318-2 | Q62318;Q62318-2 | Transcription intermediary factor 1-beta | *Trim28* | -0.370417 | [illegible] |
| Q62348 | Q62348 | Translin | *Tsn* | 0.193944 | [illegible] |
| Q62376;A0A1B0GRR8;Q62376-2;A0A1B0GR44;A0A1B0GSX5;A | Q62376;A0A1B0GRR8;Q62376-2 | U1 small nuclear ribonucleoprotein 70 kDa | *Snrnp70* | -0.234608 | [illegible] |
| Q62417-4;E9Q6A3 | Q62417-4;E9Q6A3 | | *Sorbs1* | -0.251548 | [illegible] |
| Q62418-3;Q62418-2;Q62418 | Q62418-3;Q62418-2;Q62418 | Drebrin-like protein | *Dbnl* | -0.345505 | [illegible] |
| Q62419;A0A3B2W7K0;A2ALV1;Q8BXU5;A2ALV3;Q62420 | Q62419;A0A3B2W7K0 | Endophilin-A2 | *Sh3gl1* | -0.868385 | [illegible] |
| Q62422 | Q62422 | Osteoclast-stimulating factor 1 | *Ostf1* | -0.267696 | [illegible] |
| Q62433;E9Q514;E9Q5I8;E9Q3F9;E9PVF3;F6VLR8;E9Q0J8;E9 | Q62433 | Protein NDRG1 | *Ndrg1* | -0.662949 | [illegible] |
| Q62446;A0A1Y7VLK0;A0A1Y7VP01;A0A1Y7VJ86;A0A1Y7VMJ9 | Q62446;A0A1Y7VLK0;A0A1Y7VP01 | Peptidyl-prolyl cis-trans isomerase FKBP3 | *Fkbp3* | -0.435442 | [illegible] |
| Q62465 | Q62465 | Synaptic vesicle membrane protein VAT-1 homolog | *Vat1* | -0.293239 | [illegible] |
| Q62470;Q62470-2;Q62470-3;F6UD16 | Q62470;Q62470-2;Q62470-3 | Integrin alpha-3;Integrin alpha-3 heavy chain;Integrin alpha- | *Itga3* | -0.235083 | [illegible] |
| Q62523;Q7TQE2;A0A0N4SVP4;A0A0N4SVD2;A0A0N4SUX7 | Q62523;Q7TQE2 | Zyxin | *Zyx* | 0.147181 | [illegible] |
| Q63829 | Q63829 | COMM domain-containing protein 3 | *Commd3* | -0.176777 | [illegible] |
| Q63844;A0A0U1RPX4;D3Z3G6;A0A0U1RPZ0;D3Z6D8 | Q63844;A0A0U1RPX4;D3Z3G6 | Mitogen-activated protein kinase 3;Mitogen-activated protei | *Mapk3* | -0.253304 | [illegible] |
| Q64010;Q8JZR2;Q5ND50;Q64010-2;F7D232;Q3TQV3 | Q64010;Q8JZR2;Q5ND50;Q64010-2;F7D2 | Adapter molecule crk | *Crk* | -0.290476 | [illegible] |
| Q64133 | Q64133 | Amine oxidase [flavin-containing] A | *Maoa* | -0.401967 | [illegible] |
| Q64324;F8WGM5;A0A140LJH9;A0A140LHM1;A0A140LJ60 | Q64324;F8WGM5 | Syntaxin-binding protein 2 | *Stxbp2* | -0.176777 | [illegible] |
| Q64471;D3Z5W7;D3Z3X5 | Q64471 | Glutathione S-transferase theta-1 | *Gstt1* | -0.176777 | [illegible] |
| Q64511;Q01320 | Q64511;Q01320 | DNA topoisomerase 2-beta;DNA topoisomerase 2-alpha | *Top2b;Top2a* | -0.293534 | [illegible] |
| Q64514-2;Q64514;A0A087WRC0;Q05DJ9;A0A087WP19;A0A0 | Q64514-2;Q64514;A0A087WRC0 | Tripeptidyl-peptidase 2 | *Tpp2* | -0.415421 | [illegible] |
| Q64674 | Q64674 | Spermidine synthase | *Srm* | -0.258957 | [illegible] |
| Q68FE6 | Q68FE6 | Protein FAM65A | *Fam65a* | -0.176777 | [illegible] |
| Q69ZK0;I7HPV9;Q69ZK0-2 | Q69ZK0;I7HPV9 | Phosphatidylinositol 3,4,5-trisphosphate-dependent Rac ex | *Prex1* | -0.252381 | [illegible] |
| Q6A0A2-2;Q6A0A2-3;Q6A0A2;A0A087WPU9 | Q6A0A2-2;Q6A0A2-3;Q6A0A2 | La-related protein 4B | *Larp4b* | -0.253992 | [illegible] |
| Q6DFZ1;Q6A099;H3BLN1;H3BIX5 | Q6DFZ1;Q6A099 | | *Gbf1* | -0.176777 | [illegible] |
| Q6GU68;A0A1L1STF4 | Q6GU68;A0A1L1STF4 | Immunoglobulin superfamily containing leucine-rich repeat p | *Islr* | -0.176777 | [illegible] |
| Q6IRU5-2;Q6IRU5;F7BHJ0;Q6IRU5-3 | Q6IRU5-2;Q6IRU5 | Clathrin light chain B | *Cltb* | -0.355489 | [illegible] |
| Q6KAR6;A0A1Y7VKY4;A0A1Y7VP30 | Q6KAR6;A0A1Y7VKY4 | Exocyst complex component 3 | *Exoc3* | -0.176777 | [illegible] |
| Q6KAU4;A2ARF0 | Q6KAU4;A2ARF0 | Multivesicular body subunit 12B | *Mvb12b* | -0.366921 | [illegible] |
| Q6NV83-3;Q6NV83-2;Q6NV83;A0A087WRG2 | Q6NV83-3;Q6NV83-2;Q6NV83 | U2 snRNP-associated SURP motif-containing protein | *U2surp* | -0.489752 | [illegible] |
| Q6NZR5;G3UYT1 | Q6NZR5 | | *Skiv2l* | -0.176777 | [illegible] |
| Q6P069-2;Q6P069 | Q6P069-2;Q6P069 | Sorcin | *Sri* | 5.48008 | [illegible] |
| Q6P1F6;Q9CWU3;Q6ZWR4-3;Q6ZWR4;Q6ZWR4-2;A0A286YD | Q6P1F6;Q9CWU3 | Serine/threonine-protein phosphatase 2A 55 kDa regulatory | *Ppp2r2a* | -0.673413 | [illegible] |
| Q6P4T2 | Q6P4T2 | U5 small nuclear ribonucleoprotein 200 kDa helicase | *Snrnp200* | 3.05083 | [illegible] |
| Q6P542 | Q6P542 | ATP-binding cassette sub-family F member 1 | *Abcf1* | -0.176777 | [illegible] |
| Q6P5B5;Q9WVR4 | Q6P5B5;Q9WVR4 | Fragile X mental retardation syndrome-related protein 2 | *Fxr2* | 1.47599 | [illegible] |
| Q6P5F6;A0A087WRC8;A0A087WQF7 | Q6P5F6;A0A087WRC8;A0A087WQF7 | Zinc transporter ZIP10 | *Slc39a10* | 0.828117 | [illegible] |
| Q6P5F9;A2AKT6;F6YA11 | Q6P5F9 | Exportin-1 | *Xpo1* | -0.531911 | [illegible] |
| Q6P5H2-2;Q6P5H2 | Q6P5H2-2;Q6P5H2 | Nestin | *Nes* | -0.30524 | [illegible] |
| Q6P8I4;Q3TM89;F8WI88;D3YWA4;D6RGL8;D6RDQ3 | Q6P8I4;Q3TM89;F8WI88 | PEST proteolytic signal-containing nuclear protein | *Pcnp* | -0.450497 | [illegible] |
| Q6P8X1;A0A1W2P701;A0A1W2P6Y0;Q80ZJ7 | Q6P8X1;A0A1W2P701 | Sorting nexin-6;Sorting nexin-6, N-terminally processed | *Snx6* | -0.265113 | [illegible] |
| Q6P9R4-1;Q6P9R4;Q6P9R4-2;Q6P9R4-3;A0A286YDI4;G5E8P | Q6P9R4-1;Q6P9R4;Q6P9R4-2;Q6P9R4-3 | Rho guanine nucleotide exchange factor 18 | *Arhgef18* | -0.694961 | [illegible] |
| Q6PAR5-2;Q6PAR5-6;Q6PAR5;Q6PAR5-4;Q6PAR5-5;F7ADT6 | Q6PAR5-2;Q6PAR5-6;Q6PAR5;Q6PAR5-4; | GTPase-activating protein and VPS9 domain-containing pro | *Gapvd1* | -0.176777 | [illegible] |
| Q6PB66;F6V2A3 | Q6PB66 | Leucine-rich PPR motif-containing protein, mitochondrial | *Lrpprc* | 0.0450188 | [illegible] |
| Q6PB93;Q6PB93-2 | Q6PB93;Q6PB93-2 | Polypeptide N-acetylgalactosaminyltransferase 2;Polypepti | *Galnt2* | -0.340246 | [illegible] |
| Q6PD03;A0A0A6YVV9 | Q6PD03 | Serine/threonine-protein phosphatase 2A 56 kDa regulatory | *Ppp2r5a* | -0.176777 | [illegible] |
| Q6PDL0;A0A1D5RMC1;A0A1D5RM94 | Q6PDL0;A0A1D5RMC1;A0A1D5RM94 | Cytoplasmic dynein 1 light intermediate chain 2 | *Dync1li2* | -0.363343 | [illegible] |
| Q6PFB2;Q8VE37 | Q6PFB2;Q8VE37 | Regulator of chromosome condensation | *Rcc1* | -0.176777 | [illegible] |
| Q6PGC1;Q8BN72 | Q6PGC1 | ATP-dependent RNA helicase Dhx29 | *Dhx29* | -0.176777 | [illegible] |
| Q6PGF7 | Q6PGF7 | Exocyst complex component 8 | *Exoc8* | 0.373597 | [illegible] |
| Q6PHQ8;F7A8R0;F6UFC8;F6WYJ5;E9Q6K7 | Q6PHQ8;F7A8R0 | N-alpha-acetyltransferase 35, NatC auxiliary subunit | *Naa35* | -0.176777 | [illegible] |
| Q6PIU9 | Q6PIU9 | Uncharacterized protein FLJ45252 homolog | | -0.289613 | [illegible] |
| Q6PIX5;Q6PIX5-2;A2AVE4;F2Z3W3 | Q6PIX5;Q6PIX5-2 | Inactive rhomboid protein 1 | *Rhbdf1* | -0.502783 | [illegible] |
| Q6Q477;Q6Q477-2;F7AAP4;E9Q828 | Q6Q477;Q6Q477-2;F7AAP4;E9Q828 | Calcium-transporting ATPase | *Atp2b4* | -0.305866 | [illegible] |
| Q6Q899;A2AP29;Q6Q899-2;Q6Q899-3;Q6Q899-4 | Q6Q899;A2AP29;Q6Q899-2 | Probable ATP-dependent RNA helicase DDX58 | *Ddx58* | -0.313481 | [illegible] |
| Q6R0H7;P63094;Q6R0H7-4;Z4YKV1;P63094-3;P63094-2;Q66L | Q6R0H7;P63094;Q6R0H7-4;Z4YKV1;P630 | Guanine nucleotide-binding protein G(s) subunit alpha isofo | *Gnas* | -0.368421 | [illegible] |
| Z4YJC9;Q6T3U4 | Z4YJC9;Q6T3U4 | Niemann-Pick C1-like protein 1 | *Npc1l1* | -0.176777 | [illegible] |
| Q6VH22;REV__Q8C5X9;A0A0J9YUJ7 | Q6VH22 | Intraflagellar transport protein 172 homolog | *Ift172* | -0.253046 | [illegible] |
| Q6VVW5-2;Q6VVW5;F6YT88;B1AWI7;F6YN80;Q6VVW5-3 | Q6VVW5-2;Q6VVW5;F6YT88;B1AWI7;F6Y | Atrial natriuretic peptide receptor 2;Guanylate cyclase | *Npr2* | -0.320433 | [illegible] |
| Q6WKZ8-2;Q6WKZ8;A0A286YCB2;Q6WKZ8-3 | Q6WKZ8-2;Q6WKZ8;A0A286YCB2;Q6WKZ | E3 ubiquitin-protein ligase UBR2 | *Ubr2* | 2.39082 | [illegible] |
| Q6ZPU9;E0CXN7;H3BIY2;Q6ZPU9-2;Q6ZPU9-3;E0CXJ9 | Q6ZPU9;E0CXN7;H3BIY2;Q6ZPU9-2 | KIF1-binding protein | *Kbp;2510003E04Rik* | -0.253115 | [illegible] |
| Q6ZQI3 | Q6ZQI3 | Malectin | *Mlec* | -0.668421 | [illegible] |
| Q6ZWQ7 | Q6ZWQ7 | | *Spcs3* | -0.176777 | [illegible] |
| Q6ZWQ9;Q3THE2;D3YV37;D3Z249 | Q6ZWQ9;Q3THE2 | Myosin regulatory light chain 12B | *Myl12a;Myl12b* | -0.473657 | [illegible] |
| Q6ZWV7;G3UWD7;F6YXU0 | Q6ZWV7;G3UWD7 | 60S ribosomal protein L35 | *Rpl35;Gm10269* | -0.197738 | [illegible] |
| Q6ZWX6 | Q6ZWX6 | Eukaryotic translation initiation factor 2 subunit 1 | *Eif2s1* | -0.225774 | [illegible] |
| Q6ZWY8;A0A0N4SVF0 | Q6ZWY8 | Thymosin beta-10 | *Tmsb10* | 0.424144 | [illegible] |
| Q6ZWZ7;Q9CPR4;B2RY53 | Q6ZWZ7;Q9CPR4 | 60S ribosomal protein L17 | *Rpl17* | -0.176777 | [illegible] |
| Q76MZ3;G3UWL2;G3UXQ1;H3BJ83;H3BK50 | Q76MZ3 | Serine/threonine-protein phosphatase 2A 65 kDa regulatory | *Ppp2r1a* | -0.0208054 | [illegible] |
| Q78PY7;Q3TJ56;E9Q3E9 | Q78PY7;Q3TJ56 | Staphylococcal nuclease domain-containing protein 1 | *Snd1* | 0.551978 | [illegible] |
| Q78ZM0;O70492;D3Z6Z0;D3Z789 | Q78ZM0;O70492;D3Z6Z0;D3Z789 | Sorting nexin-3 | *Snx3* | 0.499021 | [illegible] |
| Q791T5-2;Q791T5;D3YXC2 | Q791T5-2;Q791T5 | Mitochondrial carrier homolog 1 | *Mtch1* | -0.394024 | [illegible] |
| Q7M6W1;A3QM89 | Q7M6W1;A3QM89 | Reticulon | *Rtn1* | -0.176777 | [illegible] |
| Q7M6Y3-6;Q7M6Y3-5;Q7M6Y3;Q7M6Y3-2;Q7M6Y3-3;Q7M6Y3 | Q7M6Y3-6;Q7M6Y3-5;Q7M6Y3;Q7M6Y3-2; | Phosphatidylinositol-binding clathrin assembly protein | *Picalm* | 1.19637 | [illegible] |
| Q7TMB8;A0A0R4J119;Q7TMB8-2;A0A0U1RQ05;Q5SQX6;F6Q | Q7TMB8;A0A0R4J119;Q7TMB8-2;A0A0U1 | Cytoplasmic FMR1-interacting protein 1 | *Cyfip1* | -0.176777 | [illegible] |
| Q7TMC8;Q7TMC8-2;A0A1D5RMI1 | Q7TMC8;Q7TMC8-2 | | *Fuk* | 5.48008 | [illegible] |
| Q7TMM9 | Q7TMM9 | Tubulin beta-2A chain | *Tubb2a* | -0.176777 | [illegible] |
| Q7TMQ7;S4R1X1;D3Z101 | Q7TMQ7;S4R1X1 | WD repeat-containing protein 91 | *Wdr91* | -0.289121 | [illegible] |
| Q7TNC4-2;Q7TNC4-3;Q7TNC4-4;Q7TNC4;E9Q715;Q05CX5;D | Q7TNC4-2;Q7TNC4-3;Q7TNC4-4;Q7TNC4 | Putative RNA-binding protein Luc7-like 2 | *Luc7l2* | 3.32951 | [illegible] |
| Q7TNG5;Q7TNG5-2;E9QK48;D6RGM3;A0A0U1RQ42;A0A0U1 | Q7TNG5;Q7TNG5-2;E9QK48;D6RGM3 | Echinoderm microtubule-associated protein-like 2 | *Eml2* | 2.94702 | [illegible] |
| Q91V89;Q7TNL5 | Q91V89;Q7TNL5 | | *Ppp2r5d* | 0.376968 | [illegible] |
| Q7TPM6-2;Q7TPM6 | Q7TPM6-2;Q7TPM6 | Fibronectin type III and SPRY domain-containing protein 1 | *Fsd1* | -0.22781 | [illegible] |
| Q7TPV4 | Q7TPV4 | Myb-binding protein 1A | *Mybbp1a* | -0.176777 | [illegible] |
| Q7TSQ8;D6RD07;D3Z443 | Q7TSQ8;D6RD07 | Pyruvate dehydrogenase phosphatase regulatory subunit, | *Pdpr* | -0.176777 | [illegible] |
| Q7TSV4;A0A0G2JF47 | Q7TSV4 | Phosphoglucomutase-2 | *Pgm2* | 2.22028 | [illegible] |
| Q7TT15-2;Q7TT15;E9Q714 | Q7TT15-2;Q7TT15;E9Q714 | Putative polypeptide N-acetylgalactosaminyltransferase-like | *Wbscr17* | -0.314018 | [illegible] |
| Q7TT50;A0A1Y7VKD5;A0A1Y7VM95;A0A1Y7VLI0;D6RI32;E9Q | Q7TT50 | Serine/threonine-protein kinase MRCK beta | *Cdc42bpb* | -0.254088 | [illegible] |
| Q80SW1 | Q80SW1 | Putative adenosylhomocysteinase 2 | *Ahcyl1* | -0.176777 | [illegible] |
| Q80SZ7;A0A3B2WD42;D3Z7Q3 | Q80SZ7 | Guanine nucleotide-binding protein G(I)/G(S)/G(O) subunit | *Gng5* | -0.176777 | [illegible] |
| Q80U35;A0A140LIQ0;Q80U35-2 | Q80U35 | Rho guanine nucleotide exchange factor 17 | *Arhgef17* | -0.176777 | [illegible] |
| Q80U63-2;Q80U63;A2A7Y8;A8Y5E5 | Q80U63-2;Q80U63 | Mitofusin-2 | *Mfn2* | -0.176777 | [illegible] |
| Q80UP8 | Q80UP8 | Sodium-dependent phosphate transporter 2 | *Slc20a2* | -0.468311 | [illegible] |
| Q80VB6;Q9JLM8-2;A0A0G2JGQ6;Q9JLM8;A0A0G2JGR9;A8IF | Q80VB6;Q9JLM8-2;A0A0G2JGQ6;Q9JLM8 | Serine/threonine-protein kinase DCLK1 | *Dclk1* | 2.18294 | [illegible] |
| Q80VD1 | Q80VD1 | Protein FAM98B | *Fam98b* | -0.303549 | [illegible] |
| Q80VP1;Q80VP1-2;D3Z4V3;D3Z550 | Q80VP1;Q80VP1-2;D3Z4V3 | Epsin-1 | *Epn1* | 4.09379 | [illegible] |
| Q80VQ0;J3QMK6;E9Q3E1 | Q80VQ0 | Aldehyde dehydrogenase family 3 member B1 | *Aldh3b1* | -0.455211 | [illegible] |
| Q80W21;D3YVP6;D3YVP5;E9QAC8;D3YZ29 | Q80W21;D3YVP6;D3YVP5;E9QAC8;D3YZ2 | Glutathione S-transferase Mu 7 | *Gstm7* | -0.176777 | [illegible] |
| Q80W65 | Q80W65 | Proprotein convertase subtilisin/kexin type 9 | *Pcsk9* | 0.209005 | [illegible] |
| Q80WG5 | Q80WG5 | Volume-regulated anion channel subunit LRRC8A | *Lrrc8a* | -0.359785 | [illegible] |
| Q80WJ7;F6QHD1;F6ZSG0;F6ZQL0;F6QFT1;E9PUX0 | Q80WJ7;F6QHD1;F6ZSG0 | Protein LYRIC | *Mtdh* | -0.655185 | [illegible] |
| Q80X95;Q6NTA4;G3UYP1 | Q80X95;Q6NTA4 | Ras-related GTP-binding protein A;Ras-related GTP-bindin | *Rraga;Rragb* | -0.176777 | [illegible] |
| Q80XI4 | Q80XI4 | Phosphatidylinositol 5-phosphate 4-kinase type-2 beta | *Pip4k2b* | -0.354599 | [illegible] |
| Q80YS6;E9Q8X9;A0A1D5RLL0 | Q80YS6;E9Q8X9 | Actin filament-associated protein 1 | *Afap1* | -0.176777 | [illegible] |
| Q810U5-3;Q810U5-2;Q810U5;F6UK66 | Q810U5-3;Q810U5-2;Q810U5;F6UK66 | Coiled-coil domain-containing protein 50 | *Ccdc50* | -0.176777 | [illegible] |
| Q810Y8 | Q810Y8 | | *Pramel7* | -0.84114 | [illegible] |
| Q811U4;F6XRI9;D3Z4L5;D3Z5P8 | Q811U4 | Mitofusin-1 | *Mfn1* | -0.662637 | [illegible] |
| Q8BFR5;Q8BFR5-2;A0A0U1RPC4;A0A0U1RNQ6 | Q8BFR5;Q8BFR5-2 | Elongation factor Tu, mitochondrial | *Tufm* | -0.480828 | [illegible] |
| Q8BFW7;Q8BFW7-4;Q8BFW7-5;Q8BFW7-3;Q8BFW7-2 | Q8BFW7;Q8BFW7-4;Q8BFW7-5 | Lipoma-preferred partner homolog | *Lpp* | -0.176777 | [illegible] |
| Q8BFY9-2;Q8BFY9;Q3TKD0;J3QMX2 | Q8BFY9-2;Q8BFY9;Q3TKD0 | Transportin-1 | *Tnpo1* | -0.192938 | [illegible] |
| Q8BG05-2;B2RXM2 | Q8BG05-2;B2RXM2 | Heterogeneous nuclear ribonucleoprotein A3 | *Hnrnpa3;Gm6793* | -0.176777 | [illegible] |

| | | | | |
|---|---|---|---|---|
| Q8BG32;G3UYH2;G3UWW7;G3UXL5;G3UYI4;G3UYL3;G3UYL | Q8BG32 | 26S proteasome non-ATPase regulatory subunit 11 | *Psmd11* | -0.176777 |
| Q8BGC4 | Q8BGC4 | Zinc-binding alcohol dehydrogenase domain-containing pro | *Zadh2* | -0.176777 |
| Q8BGD9;B2RWE8;REV__B2RWE8;REV__Q8BGD9 | Q8BGD9;B2RWE8 | Eukaryotic translation initiation factor 4B | *Eif4b* | -0.303168 |
| Q8BGH2 | Q8BGH2 | Sorting and assembly machinery component 50 homolog | *Samm50* | -0.312809 |
| Q8BGJ5;Q922I7;Q8CB58;P17225;E9QMW9;F7AXP1;F7DCW4; | Q8BGJ5;Q922I7;Q8CB58;P17225;E9QMW | Polypyrimidine tract-binding protein 1 | *Ptbp1* | 1.03572 |
| Q8BGQ7 | Q8BGQ7 | Alanine--tRNA ligase, cytoplasmic | *Aars* | -0.31563 |
| Q8BGS2;Q8BGS2-2 | Q8BGS2;Q8BGS2-2 | BolA-like protein 2 | *Bola2* | -0.176777 |
| Q8BGT5;D6RFQ8 | Q8BGT5 | Alanine aminotransferase 2 | *Gpt2* | -0.359417 |
| Q8BGT7 | Q8BGT7 | Survival of motor neuron-related-splicing factor 30 | *Smndc1* | 0.547093 |
| Q8BH58;A0A0A6YWU8;A0A0A6YY87 | Q8BH58 | TIP41-like protein | *Tiprl* | 2.17007 |
| Q8BH64;D3Z7U7 | Q8BH64 | EH domain-containing protein 2 | *Ehd2* | 0.265643 |
| Q8BH80;Q9QY76 | Q8BH80;Q9QY76 | Vesicle-associated membrane protein-associated protein B | *Vapb* | -0.339123 |
| Q8BHC4;B0QZX9 | Q8BHC4 | Dephospho-CoA kinase domain-containing protein | *Dcakd* | 0.908414 |
| Q8BHF7;F8WI04;F2Z427 | Q8BHF7;F8WI04 | CDP-diacylglycerol--glycerol-3-phosphate 3-phosphatidyltr | *Pgs1* | -0.176777 |
| Q8BHJ8;Q9CXY1 | Q8BHJ8;Q9CXY1 | Transmembrane protein 175 | *Tmem175* | -0.315449 |
| Q8BHL3 | Q8BHL3 | TBC1 domain family member 10B | *Tbc1d10b* | 0.207569 |
| Q8BHL8 | Q8BHL8 | Proteasome inhibitor PI31 subunit | *Psmf1* | -0.176777 |
| Q8BHN3;Q8BHN3-2;Q8BHN3-3 | Q8BHN3;Q8BHN3-2;Q8BHN3-3 | Neutral alpha-glucosidase AB | *Ganab* | -0.435879 |
| Q8BI72;A0A1I7Q4A1 | Q8BI72 | CDKN2A-interacting protein | *Cdkn2aip* | -0.305943 |
| Q8BIJ6;E9PWN2;E9PWN3 | Q8BIJ6;E9PWN2;E9PWN3 | Isoleucine--tRNA ligase, mitochondrial | *Iars2* | -0.253226 |
| Q8BJ71;A0A1D5RLQ0;Q8BJ71-2;A0A1D5RM86 | Q8BJ71;A0A1D5RLQ0;Q8BJ71-2 | Nuclear pore complex protein Nup93 | *Nup93* | -0.176777 |
| Q8BJS4-3;Q8BJS4-2;Q8BJS4 | Q8BJS4-3;Q8BJS4-2;Q8BJS4 | SUN domain-containing protein 2 | *Sun2* | -0.176777 |
| Q8BJU0-2;Q8BJU0;A0A1W2P6P1;A0A1W2P7I5 | Q8BJU0-2;Q8BJU0;A0A1W2P6P1;A0A1W2 | Small glutamine-rich tetratricopeptide repeat-containing pro | *Sgta* | -0.633136 |
| Q8BJY1;F7BA91;REV__A2AQP0 | Q8BJY1 | 26S proteasome non-ATPase regulatory subunit 5 | *Psmd5* | 1.01368 |
| Q8BK64;A0A1Y7VM19 | Q8BK64 | Activator of 90 kDa heat shock protein ATPase homolog 1 | *Ahsa1* | -0.211036 |
| Q8BKC5;Q8BKC5-2 | Q8BKC5;Q8BKC5-2 | Importin-5 | *Ipo5* | -0.251761 |
| Q8BKG3 | Q8BKG3 | Inactive tyrosine-protein kinase 7 | *Ptk7* | 0.926871 |
| Q8BLF1;Q8BYQ0;D6RGP7 | Q8BLF1 | Neutral cholesterol ester hydrolase 1 | *Nceh1* | -0.350666 |
| Q8BMD8 | Q8BMD8 | Calcium-binding mitochondrial carrier protein SCaMC-1 | *Slc25a24* | 1.16578 |
| Q8BMF3 | Q8BMF3 | NADP-dependent malic enzyme, mitochondrial | *Me3* | -0.561058 |
| Q8BMF4 | Q8BMF4 | Dihydrolipoyllysine-residue acetyltransferase component of | *Dlat* | -0.315216 |
| Q8BMJ2 | Q8BMJ2 | Leucine--tRNA ligase, cytoplasmic | *Lars* | -0.455089 |
| Q8BMK4 | Q8BMK4 | Cytoskeleton-associated protein 4 | *Ckap4* | -0.341493 |
| Q8BMS1 | Q8BMS1 | Trifunctional enzyme subunit alpha, mitochondrial;Long-cha | *Hadha* | -0.731967 |
| Q8BND5-2;Q8BND5;Q8BND5-3;Q8BND5-4 | Q8BND5-2;Q8BND5;Q8BND5-3 | Sulfhydryl oxidase 1 | *Qsox1* | -0.372617 |
| Q8BNE1-2;Q8BNE1-3;Q8BNE1 | Q8BNE1-2;Q8BNE1-3;Q8BNE1 | TRPM8 channel-associated factor 1 | *Tcaf1* | -0.176777 |
| Q8BNW9 | Q8BNW9 | Kelch repeat and BTB domain-containing protein 11 | *Kbtbd11* | -0.176777 |
| Q8BP40 | Q8BP40 | Lysophosphatidic acid phosphatase type 6 | *Acp6* | 4.05533 |
| Q8BP47 | Q8BP47 | Asparagine--tRNA ligase, cytoplasmic | *Nars* | -0.3652 |
| Q8BP48;A0A0G2JFL1;A0A0G2JF71 | Q8BP48 | Methionine aminopeptidase 1 | *Metap1* | -0.304006 |
| Q8BP67;F6RSK3 | Q8BP67 | 60S ribosomal protein L24 | *Rpl24* | -0.260627 |
| Q8BP92 | Q8BP92 | Reticulocalbin-2 | *Rcn2* | -0.396126 |
| Q8BPB5;F6ZFS0 | Q8BPB5;F6ZFS0 | EGF-containing fibulin-like extracellular matrix protein 1 | *Efemp1* | 0.631963 |
| Q8BPU7;Q8BPU7-4;Q8BPU7-2;Q8BPU7-3;A0A1Y7VIX9 | Q8BPU7;Q8BPU7-4;Q8BPU7-2 | Engulfment and cell motility protein 1 | *Elmo1* | -0.24997 |
| Q8BQ30;Q8BQ30-2;A0A087WRI9 | Q8BQ30;Q8BQ30-2 | Phostensin | *Ppp1r18* | 0.740997 |
| Q8BR90;Q8BR90-2 | Q8BR90;Q8BR90-2 | UPF0600 protein C5orf51 homolog | | -0.176777 |
| Q8BSK8-2;Q8BSK8;D6RH86;E9PUN3;D3YWB8;Q9Z1M4 | Q8BSK8-2;Q8BSK8;D6RH86;E9PUN3;D3Y | Ribosomal protein S6 kinase beta-1;Ribosomal protein S6 k | *Rps6kb1;Rps6kb2* | -0.176777 |
| Q8BT60;Q9D6C8;A0A0R4J0J1;Q3UYN2;Q1RLL3;Q9Z140;Q8E | Q8BT60 | Copine-3 | *Cpne3* | -0.541728 |
| Q8BTI8-3;Q8BTI8;Q8BTI8-2;A0A087WRX8;A0A3B2W3Z4;A0A0 | Q8BTI8-3;Q8BTI8;Q8BTI8-2 | Serine/arginine repetitive matrix protein 2 | *Srrm2* | -0.247861 |
| Q8BTS3 | Q8BTS3 | | *Gbp9* | -0.176777 |
| Q8BTS4 | Q8BTS4 | Nuclear pore complex protein Nup54 | *Nup54* | -0.176777 |
| Q8BTZ7 | Q8BTZ7 | Mannose-1-phosphate guanyltransferase beta | *Gmppb* | 5.48008 |
| Q8BU30;E9Q866;F6Q6R1 | Q8BU30 | Isoleucine--tRNA ligase, cytoplasmic | *Iars* | -0.231657 |
| Q8BUE4;Q8BUE4-2 | Q8BUE4;Q8BUE4-2 | Apoptosis-inducing factor 2 | *Aifm2* | -0.176777 |
| Q8BUK6;F6W1A2 | Q8BUK6 | Protein Hook homolog 3 | *Hook3* | -0.176777 |
| Q8BUR4;Q8BUR4-2;A0A1B0GSI1;A0A1B0GS53;B2RY04 | Q8BUR4;Q8BUR4-2 | Dedicator of cytokinesis protein 1 | *Dock1* | -0.42169 |
| Q8BVA0;Q99P91 | Q8BVA0;Q99P91 | Transmembrane glycoprotein NMB | *Gpnmb* | -0.840575 |
| Q8BVL3;D3Z3J1;H3BKG9;D3Z7S9;H3BKQ6;REV__Q9CY73-2;I | Q8BVL3 | Sorting nexin-17 | *Snx17* | -0.49135 |
| Q8BVQ5;A0A140LI84 | Q8BVQ5 | Protein phosphatase methylesterase 1 | *Ppme1* | -0.364121 |
| Q8BVU5;A2AH27;A2AH28;A2AH29;A2AH30 | Q8BVU5;A2AH27 | ADP-ribose pyrophosphatase, mitochondrial | *Nudt9* | -0.176777 |
| Q8BW96-3;Q8BW96;Q8BW96-2 | Q8BW96-3;Q8BW96;Q8BW96-2 | Calcium/calmodulin-dependent protein kinase type 1D | *Camk1d* | -0.176777 |
| Q8BWM0 | Q8BWM0 | Prostaglandin E synthase 2;Prostaglandin E synthase 2 tru | *Ptges2* | -0.176777 |
| Q8BWP8;Q8BWP8-2 | Q8BWP8;Q8BWP8-2 | Beta-1,4-glucuronyltransferase 1 | *B4gat1* | -0.253081 |
| Q8BWW3;Q9CYR6 | Q8BWW3;Q9CYR6 | Phosphoacetylglucosamine mutase | *Pgm3* | 0.61883 |
| Q8BWY3 | Q8BWY3 | Eukaryotic peptide chain release factor subunit 1 | *Etf1* | -0.176777 |
| Q8BWZ3-2;Q8BWZ3 | Q8BWZ3-2;Q8BWZ3 | N-alpha-acetyltransferase 25, NatB auxiliary subunit | *Naa25* | 2.48135 |
| Q8BXZ1 | Q8BXZ1 | Protein disulfide-isomerase TMX3 | *Tmx3* | -0.176777 |
| Q8BYA0;F6UI15;B1ATU0;Q9D2H8 | Q8BYA0 | Tubulin-specific chaperone D | *Tbcd* | -0.176777 |
| Q8BYH7;I6L974 | Q8BYH7;I6L974 | TBC1 domain family member 17 | *Tbc1d17* | -0.176777 |
| Q8BZ20 | Q8BZ20 | Poly [ADP-ribose] polymerase 12 | *Parp12* | -0.176777 |
| Q8BZA9;A0A0J9YV11 | Q8BZA9 | Fructose-2,6-bisphosphatase TIGAR | *Tigar* | -0.234988 |
| Q8BZQ7 | Q8BZQ7 | Anaphase-promoting complex subunit 2 | *Anapc2* | -0.733169 |
| Q8BZW8 | Q8BZW8 | NHL repeat-containing protein 2 | *Nhlrc2* | -0.176777 |
| Q8C052;A0A1D5RLY6 | Q8C052;A0A1D5RLY6 | Microtubule-associated protein 1S;MAP1S heavy chain;MA | *Map1s* | -0.306926 |
| Q8C0E2;Q8C0E2-2 | Q8C0E2 | Vacuolar protein sorting-associated protein 26B | *Vps26b* | -0.176777 |
| Q8C0L6;Q3TXR6;Q8C0L6-2 | Q8C0L6;Q3TXR6 | Peroxisomal N(1)-acetyl-spermine/spermidine oxidase | *Paox* | -0.415558 |
| Q8C0N2;Q8K2C8 | Q8C0N2;Q8K2C8 | Glycerol-3-phosphate acyltransferase 3;Glycerol-3-phosph | *Agpat9;Agpat6* | 0.0678728 |
| Q8C129 | Q8C129 | Leucyl-cystinyl aminopeptidase | *Lnpep* | 5.48008 |
| Q8C1Q4;Q8C1Q4-2 | Q8C1Q4 | Meteorin | *Metrn* | -0.74717 |
| Q8C2Q3;E9QL13;Q8C2Q3-2;J3QN51;J3QPT3;B0LM42 | Q8C2Q3;E9QL13;Q8C2Q3-2 | RNA-binding protein 14 | *Rbm14* | -0.414857 |
| Q8C3J5;Q5SRI3;Q3TMS1 | Q8C3J5;Q5SRI3 | Dedicator of cytokinesis protein 2 | *Dock2* | -0.176777 |
| Q8C3W1 | Q8C3W1 | Uncharacterized protein C1orf198 homolog | | -0.244131 |
| Q8C407;Q8C407-2 | Q8C407;Q8C407-2 | Protein YIPF4 | *Yipf4* | -0.176777 |
| Q8C4S8;D3Z1C6 | Q8C4S8 | DENN domain-containing protein 2A | *Dennd2a* | -0.412861 |
| Q8C570;A2ASC6 | Q8C570 | mRNA export factor | *Rae1* | -0.176777 |
| Q8C5G6;Q9QZ06;F7AT44;A9JEI5 | Q8C5G6;Q9QZ06;F7AT44;A9JEI5 | Toll-interacting protein | *Tollip* | -0.176777 |
| Q8C5H8-2;Q8C5H8-3;Q8C5H8;Z4YK42;Q8C5H8-4 | Q8C5H8-2;Q8C5H8-3;Q8C5H8;Z4YK42;Q8 | NAD kinase 2, mitochondrial | *Nadk2* | -0.176777 |
| Q8C5W3;Q8C5W3-2;D3YVP4 | Q8C5W3;Q8C5W3-2 | Tubulin-specific chaperone cofactor E-like protein | *Tbcel* | -0.893919 |
| Q8C6I2-3;Q8C6I2 | Q8C6I2-3;Q8C6I2 | Succinate dehydrogenase assembly factor 2, mitochondrial | *Sdhaf2* | -0.510171 |
| Q8C754;G3UY33;Q8C754-2 | Q8C754;G3UY33;Q8C754-2 | Vacuolar protein sorting-associated protein 52 homolog | *Vps52* | -0.24988 |
| Q8C7K6;A0A0A6YY26 | Q8C7K6 | Prenylcysteine oxidase-like | *Pcyox1l* | -0.176777 |
| Z4YJW0;Q8C7X2-2;Q8C7X2;Q8C7X2-3;A0A1Y7VP81 | Z4YJW0;Q8C7X2-2;Q8C7X2 | ER membrane protein complex subunit 1 | *Emc1* | -0.251717 |
| Q8C854-3;Q8C854;A2ATP5;A2ATP6;G8JL68;Q8C854-1;Q8C8 | Q8C854-3;Q8C854;A2ATP5;A2ATP6;G8JL | Myelin expression factor 2 | *Myef2* | -0.411361 |
| Q8C863;Q8C863-2 | Q8C863;Q8C863-2 | E3 ubiquitin-protein ligase Itchy | *Itch* | -0.245183 |
| Q8CAG6;Q9JHK5;Q5F270 | Q8CAG6;Q9JHK5 | Pleckstrin | *Plek* | -0.314288 |
| Q8CAQ8-2;Q8CAQ8;Q8CAQ8-5;E9Q800;E9QAY6;A0A0U1RQ | Q8CAQ8-2;Q8CAQ8;Q8CAQ8-5;E9Q800;E | MICOS complex subunit Mic60 | *Immt* | -0.445188 |
| Q8CBY8-2;Q8CBY8;Q3TRR2 | Q8CBY8-2;Q8CBY8 | Dynactin subunit 4 | *Dctn4* | -0.176777 |
| Q8CC88;Q8CC88-2 | Q8CC88;Q8CC88-2 | von Willebrand factor A domain-containing protein 8 | *Vwa8* | -0.176777 |
| Q8CCF0;Q8CCF0-2 | Q8CCF0;Q8CCF0-2 | U4/U6 small nuclear ribonucleoprotein Prp31 | *Prpf31* | -0.277558 |
| Q8CCT4;A2AJS2;A2AJS1 | Q8CCT4 | Transcription elongation factor A protein-like 5 | *Tceal5* | -0.639908 |
| Q8CDA1;Q8CDA1-3;Q8CDA1-2;Q8CDA1-4 | Q8CDA1;Q8CDA1-3;Q8CDA1-2 | Phosphatidylinositide phosphatase SAC2 | *Inpp5f* | 5.34317 |
| Q8CDN6 | Q8CDN6 | Thioredoxin-like protein 1 | *Txnl1* | -0.547379 |
| Q8CFA2 | Q8CFA2 | Aminomethyltransferase, mitochondrial | *Amt* | -0.252539 |
| Q8CFR5;Q9D2N4-2;A0A1Y7VL34;Q9D2N4;Q9D2N4-7;A0A1Y7 | Q8CFR5;Q9D2N4-2;A0A1Y7VL34;Q9D2N4 | Dystrobrevin;Dystrobrevin alpha | *Dtna* | -0.58323 |
| Q8CG16;Q8CFG9;Q3UZ09 | Q8CG16;Q8CFG9 | Complement C1r-A subcomponent;Complement C1r-A sub | *C1ra;C1rb* | -0.296097 |
| Q8CGA0;A0A338P7F2 | Q8CGA0 | Protein phosphatase 1F | *Ppm1f* | -0.176777 |
| Q8CGF7-2;Q8CGF7;Q8CGF7-3;Q3B807-3;Q3B807 | Q8CGF7-2;Q8CGF7;Q8CGF7-3 | Transcription elongation regulator 1 | *Tcerg1* | -0.481456 |
| Q8CGK3 | Q8CGK3 | Lon protease homolog, mitochondrial | *Lonp1* | -0.232887 |
| Q8CH18-3;Q8CH18;Q8CH18-2;A0A1W2P765 | Q8CH18-3;Q8CH18 | Cell division cycle and apoptosis regulator protein 1 | *Ccar1* | -0.176777 |
| Q8CHX7;Q8CHX7-2 | Q8CHX7;Q8CHX7-2 | Raftlin-2 | *Rftn2* | -0.176777 |
| Q8CI08;Q8CI08-2;A0A0J9YUJ0 | Q8CI08;Q8CI08-2;A0A0J9YUJ0 | SLAIN motif-containing protein 2 | *Slain2* | 0.715489 |
| Q8CI70 | Q8CI70 | Leucine-rich repeat-containing protein 20 | *Lrrc20* | -0.37815 |
| Q8CI94;Q9WUB3;E9PUM3 | Q8CI94 | Glycogen phosphorylase, brain form | *Pygb* | 0.402697 |
| Q8CIB5;A6X940;A6X941;A0A2I3BPQ7 | Q8CIB5 | Fermitin family homolog 2 | *Fermt2* | 0.251097 |
| Q8CIE6;F8WHL2;F6XJN3 | Q8CIE6;F8WHL2 | Coatomer subunit alpha;Xenin;Proxenin;Coatomer subunit a | *Copa* | 1.12794 |
| Q8CIG8;A0A0R4J049;F6QQQ6;A0A2I3BRG2;S4R295 | Q8CIG8;A0A0R4J049 | Protein arginine N-methyltransferase 5 | *Prmt5* | 1.74906 |
| Q8CII2;Q8CII2-2;A0A0A6YVX4 | Q8CII2;Q8CII2-2 | Cell division cycle protein 123 homolog | *Cdc123* | -0.251728 |
| Q8CIN4 | Q8CIN4 | Serine/threonine-protein kinase PAK 2;PAK-2p27;PAK-2p34 | *Pak2* | 1.43299 |
| Q8CJ53-4;Q8CJ53-3;Q8CJ53-2;Q8CJ53 | Q8CJ53-4;Q8CJ53-3;Q8CJ53-2;Q8CJ53 | Cdc42-interacting protein 4 | *Trip10* | -0.176777 |
| Q8CJG0 | Q8CJG0 | Protein argonaute-2 | *Aug-02* | -0.242948 |
| Q8CJH3;A0A0G2JDM9 | Q8CJH3 | Plexin-B1 | *Plxnb1* | -0.176777 |
| Q8JZK9;P54869 | Q8JZK9 | Hydroxymethylglutaryl-CoA synthase, cytoplasmic | *Hmgcs1* | -0.694737 |
| Q8JZQ2 | Q8JZQ2 | AFG3-like protein 2 | *Afg3l2* | -0.176777 |
| Q8JZU2;F6VVY4 | Q8JZU2;F6VVY4 | Tricarboxylate transport protein, mitochondrial | *Slc25a1* | -0.176777 |
| Q8JZX4;A2AP40 | Q8JZX4 | Splicing factor 45 | *Rbm17* | -0.508796 |
| Q8JZZ5 | Q8JZZ5 | | *Pitpnb* | -0.176777 |
| Q8K010;E9Q484;A0A1B0GSQ4;E9PZB8 | Q8K010;E9Q484 | 5-oxoprolinase | *Oplah* | -0.252115 |
| Q8K0L9-2;Q8K0L9;E0CXT9;E0CXK1;E0CYJ9;E0CYH3;Q52KG | Q8K0L9-2;Q8K0L9 | Zinc finger and BTB domain-containing protein 20 | *Zbtb20* | -0.226033 |
| Q8K0T0 | Q8K0T0 | Reticulon-1 | *Rtn1* | -0.278456 |
| Q8K124 | Q8K124 | Pleckstrin homology domain-containing family O member 2 | *Plekho2* | -0.176777 |
| Z4YK56;Q8K135;Q8K135-2;A2A786;A2A787;A2A788 | Z4YK56;Q8K135;Q8K135-2;A2A786;A2A78 | Dyslexia-associated protein KIAA0319-like protein | *AU040320;Kiaa0319l* | -0.253417 |
| Q8K1M6-3;E9PUD2;Q8K1M6-2;Q8K1M6;Q8K1M6-4;A0A2U3TZ | Q8K1M6-3;E9PUD2;Q8K1M6-2;Q8K1M6;Q | Dynamin-1-like protein | *Dnm1l* | -0.176777 |
| Q8K1R3;Q8K1R3-2 | Q8K1R3;Q8K1R3-2 | Polyribonucleotide nucleotidyltransferase 1, mitochondrial | *Pnpt1* | -0.176777 |
| Q8K1R7 | Q8K1R7 | Serine/threonine-protein kinase Nek9 | *Nek9* | -0.176777 |
| Q8K2F8;F6X2V9;A0A0U1RP30;A0A0U1RP90;A2BH69 | Q8K2F8;F6X2V9 | Protein LSM14 homolog A | *Lsm14a* | -0.176777 |
| Q8K2L8;Q8C0C3 | Q8K2L8 | Trafficking protein particle complex subunit 12 | *Trappc12* | -0.176777 |
| Q8K2Y7 | Q8K2Y7 | 39S ribosomal protein L47, mitochondrial | *Mrpl47* | -0.176777 |
| Q8K341-5;Q8K341-4;Q8K341-3;Q8K341-2;Q8K341;F6RPX5;F7 | Q8K341-5;Q8K341-4;Q8K341-3;Q8K341-2; | Alpha-tubulin N-acetyltransferase 1 | *Atat1* | -0.241953 |
| Q8K3G9;D3Z340;H3BL43;D6RFR7 | Q8K3G9 | DCC-interacting protein 13-beta | *Appl2* | -0.540228 |
| Q8K3H0 | Q8K3H0 | DCC-interacting protein 13-alpha | *Appl1* | -0.366611 |
| Q8K411-3;Q8K411-2;Q8K411 | Q8K411-3;Q8K411-2;Q8K411 | Presequence protease, mitochondrial | *Pitrm1* | -0.176777 |
| Q8K4K1-2;Q8K4K1;E9PVN5;A0A0A0MQN3;Q8K4K1-3 | Q8K4K1-2;Q8K4K1;E9PVN5;A0A0A0MQN3 | Centrin-4 | *Cetn4* | -0.253964 |
| Q8K4Q8 | Q8K4Q8 | Collectin-12 | *Colec12* | 0.896829 |
| Q8K4X7;E9Q2T9;E9Q6S5 | Q8K4X7 | 1-acyl-sn-glycerol-3-phosphate acyltransferase delta | *Agpat4* | -0.229862 |
| Q8N7N5;Q8N7N5-2;A0A0A6YWY2 | Q8N7N5;Q8N7N5-2 | DDB1- and CUL4-associated factor 8 | *Dcaf8* | -0.253164 |
| Q8QZT1 | Q8QZT1 | Acetyl-CoA acetyltransferase, mitochondrial | *Acat1* | -0.253609 |
| Q8R010;Q8R3V2 | Q8R010;Q8R3V2 | Aminoacyl tRNA synthase complex-interacting multifunctior | *Aimp2* | 0.166038 |
| Q8R016;E9PY26;E9QA53;E9Q6V3;E9PZH4 | Q8R016;E9PY26 | Bleomycin hydrolase | *Blmh* | 0.660993 |
| Q8R050-2;Q8R050;Q149F3;F7CE88;REV__S4R1J5;REV__S4F | Q8R050-2;Q8R050 | Eukaryotic peptide chain release factor GTP-binding subun | *Gspt1* | -0.176777 |
| Q8R086 | Q8R086 | Sulfite oxidase, mitochondrial | *Suox* | 1.1102 |
| Q8R0X7;D6REF7;D3YZT4;D3YY13;D3Z1Z3 | Q8R0X7;D6REF7 | Sphingosine-1-phosphate lyase 1 | *Sgpl1* | -0.334769 |
| Q8R180;A0A2I3BPM1 | Q8R180;A0A2I3BPM1 | ERO1-like protein alpha | *Ero1l* | -0.468248 |
| Q8R1F1;A2ARS6 | Q8R1F1 | Niban-like protein 1 | *Fam129b* | 0.489806 |
| Q8R1I1 | Q8R1I1 | Cytochrome b-c1 complex subunit 9 | *Uqcr10* | 0.113674 |
| Q8R1Q8 | Q8R1Q8 | Cytoplasmic dynein 1 light intermediate chain 1 | *Dync1li1* | -0.176777 |
| Q8R2G6 | Q8R2G6 | Coiled-coil domain-containing protein 80 | *Ccdc80* | -0.176777 |
| Q8R2U6 | Q8R2U6 | Diphosphoinositol polyphosphate phosphohydrolase 2 | *Nudt4* | -0.254114 |
| Q8R2Y2-2;Q8R2Y2;A0A1L1SQC5;F7ATH5 | Q8R2Y2-2;Q8R2Y2 | Cell surface glycoprotein MUC18 | *Mcam* | 2.2383 |
| Q8R307 | Q8R307 | Vacuolar protein sorting-associated protein 18 homolog | *Vps18* | -0.176777 |
| Q8R317-2;Q8R317;A0A286YE09 | Q8R317-2;Q8R317 | Ubiquilin-1 | *Ubqln1* | -0.176777 |
| Q8R366;G3UYZ1;A0A0R4J117;A0A0A6YWK7;A0A0A6YVX9;D3 | Q8R366;G3UYZ1;A0A0R4J117 | Immunoglobulin superfamily member 8 | *Igsf8* | -0.176777 |
| Q8R3C0;H9KV04 | Q8R3C0;H9KV04 | Mini-chromosome maintenance complex-binding protein | *Mcmbp* | 3.89962 |
| Q8R3V5-1;Q8R3V5;A2AWI9;A2AWI7;Q8R3V5-3 | Q8R3V5-1;Q8R3V5;A2AWI9;A2AWI7;Q8R3 | Endophilin-B2 | *Sh3glb2* | -0.176777 |
| Q8R422 | Q8R422 | CD109 antigen | *Cd109* | -0.0518236 |
| Q8R464 | Q8R464 | Cell adhesion molecule 4 | *Cadm4* | -0.22864 |
| Q8R4G6 | Q8R4G6 | Alpha-1,6-mannosylglycoprotein 6-beta-N-acetylglucosami | *Mgat5* | -0.176777 |
| Q8R4R6;A2ATJ2 | Q8R4R6 | Nucleoporin NUP53 | *Nup35* | -0.410187 |
| Q8R4U7;E9Q9L3 | Q8R4U7 | Leucine zipper protein 1 | *Luzp1* | -0.447888 |
| Q8R570;B2FDF6;Q8R570-2;I7HPB1;I7HFU4;J3JS29 | Q8R570;B2FDF6;Q8R570-2 | Synaptosomal-associated protein 47 | *Snap47* | -0.176777 |
| Q8R5C5;E0CZD4;E0CYB4 | Q8R5C5 | Beta-centractin | *Actr1b* | 0.361334 |
| Q8R5L3-2;Q8R5L3 | Q8R5L3-2;Q8R5L3 | Vam6/Vps39-like protein | *Vps39* | -0.252898 |
| Q8VCC9;Q3UYD7 | Q8VCC9 | Spondin-1 | *Spon1* | -0.247199 |
| Q8VCI5;Q8VCI5-2 | Q8VCI5;Q8VCI5-2 | Peroxisomal biogenesis factor 19 | *Pex19* | -0.176777 |
| Q8VCN5 | Q8VCN5 | Cystathionine gamma-lyase | *Cth* | -0.236421 |
| Q8VCT3;E9PYF1 | Q8VCT3;E9PYF1 | Aminopeptidase B | *Rnpep* | -0.323862 |
| Q8VCV2;Q9QYF9;Q8CBD0 | Q8VCV2;Q9QYF9;Q8CBD0 | Protein NDRG3 | *Ndrg3* | -0.176777 |
| Q8VCX5;Q8VCX5-3;Q8VCX5-2;F6Z1Z9;E9Q9E0 | Q8VCX5;Q8VCX5-3;Q8VCX5-2;F6Z1Z9;E9 | Calcium uptake protein 1, mitochondrial | *Micu1* | -0.176777 |
| Q8VD75;A0A0J9YUA3;A0A1D5RMM7;A0A0J9YV08;A0A0J9YU) | Q8VD75;A0A0J9YUA3 | Huntingtin-interacting protein 1 | *Hip1* | -0.176777 |
| Q8VDC0;A0A1L1SQK2;A0A1L1SSW1 | Q8VDC0 | Probable leucine--tRNA ligase, mitochondrial | *Lars2* | -0.302335 |
| Q8VDJ3;A0A087WP83;A0A087WS92;A0A087WQY9;A0A087WF | Q8VDJ3;A0A087WP83 | Vigilin | *Hdlbp* | -0.2709 |
| Q8VDM4;A0A338P6M5;E9Q2S8;A0A338P6S9;A0A338P7H4 | Q8VDM4 | 26S proteasome non-ATPase regulatory subunit 2 | *Psmd2* | 0.554212 |
| Q8VDM6-2;Q8VDM6;Q8VDM6-3 | Q8VDM6-2;Q8VDM6 | Heterogeneous nuclear ribonucleoprotein U-like protein 1 | *Hnrnpul1* | -0.262475 |
| Q8VDN2;Q91WH7;E9QNX7;Q64436 | Q8VDN2 | Sodium/potassium-transporting ATPase subunit alpha-1 | *Atp1a1* | 3.40543 |
| Q8VDP3-3;Q8VDP3;Q8VDP3-2;E9PUI4 | Q8VDP3-3;Q8VDP3;Q8VDP3-2;E9PUI4 | Protein-methionine sulfoxide oxidase MICAL1 | *Mical1* | -0.337855 |
| Q8VDQ1;Q3TXN1;Q8VDQ1-2;D6RGL6 | Q8VDQ1;Q3TXN1;Q8VDQ1-2 | Prostaglandin reductase 2 | *Ptgr2* | -0.543184 |
| Q8VEH5 | Q8VEH5 | EPM2A-interacting protein 1 | *Epm2aip1* | -0.176777 |
| Q8VEH8;F6Z458 | Q8VEH8;F6Z458 | Endoplasmic reticulum lectin 1 | *Erlec1* | -0.25226 |
| Q8VEK3-2;Q8VEK3 | Q8VEK3-2;Q8VEK3 | Heterogeneous nuclear ribonucleoprotein U | *Hnrnpu* | -0.186776 |
| Q8VHM5;F7B5B5;A2AW41;A2AW40;V9GWW3;S4R1E9 | Q8VHM5;F7B5B5;A2AW41 | | *Hnrnpr* | -0.176777 |
| Q8VHX6-2;Q8VHX6 | Q8VHX6-2;Q8VHX6 | Filamin-C | *Flnc* | -0.425882 |
| Q8VHY0;Q8VHY0-3;Q8VHY0-2 | Q8VHY0;Q8VHY0-3 | Chondroitin sulfate proteoglycan 4 | *Cspg4* | -0.176777 |

| | | | | | | | | | | | | | | | | | | | | | | | | | | | | | | | | | | | | | | | | | | | | | | |
|---|---|---|---|---|---|---|---|---|---|---|---|---|---|---|---|---|---|---|---|---|---|---|---|---|---|---|---|---|---|---|---|---|---|---|---|---|---|---|---|---|---|---|---|---|---|---|
| Q91V35;P18052-2;P18052;A0A2R8VHH6;A0A1B0GRH2;A0A1B | Q91V35;P18052-2;P18052 | Receptor-type tyrosine-protein phosphatase;Receptor-type | *Ptpra* | -0.357541 | -0.357541 | -0.357541 | 1.82172 | -0.357541 | -0.357541 | -0.357541 | -0.357541 | -0.357541 | -0.357541 | -0.357541 | -0.357541 | -0.357541 | -0.357541 | -0.357541 | -0.357541 | -0.357541 | -0.357541 | -0.357541 | -0.357541 | -0.357541 | -0.357541 | -0.357541 | -0.357541 | -0.357541 | -0.357541 | -0.357541 | 2.62078 | 1.85466 | -0.357541 | -0.357541 | 3.71399 | 14 | 23.6 | 23.6 | 89.841 | 135.05 | 1418700000 | 28 | 0 | 0.863095 | 0 | 0 |
| Q91V41;Q50HX3;A0A2R8VHW9 | Q91V41;Q50HX3 | Ras-related protein Rab-14 | *Rab14* | -0.19078 | -0.19078 | -0.19078 | -0.19078 | -0.19078 | -0.19078 | -0.19078 | -0.19078 | -0.19078 | -0.19078 | -0.19078 | -0.19078 | -0.19078 | -0.19078 | -0.19078 | -0.19078 | -0.19078 | -0.19078 | 5.46261 | -0.19078 | -0.19078 | -0.19078 | -0.19078 | 0.260802 | -0.19078 | -0.19078 | -0.19078 | -0.19078 | -0.19078 | -0.19078 | -0.19078 | -0.19078 | 16 | 73.5 | 73.5 | 23.897 | 323.31 | 11230000000 | 63 | 0.581812 | 0.560593 | 0.872138 | 1.17277 |
| Q91V61-2;Q91V61;Q3U4F0 | Q91V61-2;Q91V61;Q3U4F0 | Sideroflexin-3 | *Sfxn3* | -0.176777 | -0.176777 | -0.176777 | -0.176777 | -0.176777 | -0.176777 | -0.176777 | -0.176777 | -0.176777 | -0.176777 | 5.48008 | -0.176777 | -0.176777 | -0.176777 | -0.176777 | -0.176777 | -0.176777 | -0.176777 | -0.176777 | -0.176777 | -0.176777 | -0.176777 | -0.176777 | -0.176777 | -0.176777 | -0.176777 | -0.176777 | -0.176777 | -0.176777 | -0.176777 | -0.176777 | -0.176777 | 11 | 46.5 | 46.5 | 31.656 | 138.76 | 5903300000 | 33 | 0.433155 | 0.611797 | -0.707107 | -0.930949 |
| Q91V76;A0A1L1SSF8;A0A1L1ST86 | Q91V76;A0A1L1SSF8 | Ester hydrolase C11orf54 homolog | | -0.298174 | -0.298174 | -0.298174 | -0.298174 | -0.298174 | -0.298174 | -0.298174 | -0.298174 | 1.53271 | -0.298174 | -0.298174 | -0.298174 | -0.298174 | 2.93842 | 4.1759 | -0.298174 | -0.298174 | -0.298174 | -0.298174 | -0.298174 | -0.298174 | -0.298174 | -0.298174 | -0.298174 | -0.298174 | -0.298174 | -0.298174 | -0.298174 | -0.298174 | -0.298174 | -0.298174 | -0.298174 | 15 | 51.1 | 51.1 | 34.995 | 323.31 | 5385200000 | 47 | 0.731797 | 0.513714 | -0.963834 | -1.39824 |
| Q91V87-2;Q91V87 | Q91V87-2;Q91V87 | Fibroblast growth factor receptor-like 1 | *Fgfrl1* | -0.176777 | -0.176777 | -0.176777 | -0.176777 | -0.176777 | -0.176777 | -0.176777 | -0.176777 | -0.176777 | -0.176777 | -0.176777 | -0.176777 | -0.176777 | -0.176777 | 5.48008 | -0.176777 | -0.176777 | -0.176777 | -0.176777 | -0.176777 | -0.176777 | -0.176777 | -0.176777 | -0.176777 | -0.176777 | -0.176777 | -0.176777 | -0.176777 | -0.176777 | -0.176777 | -0.176777 | -0.176777 | 2 | 7.1 | 7.1 | 46.98 | 49.924 | 426190000 | 6 | 0.433155 | 0.611279 | -0.707107 | -0.930949 |
| Q91V98 | Q91V98 | Endosialin | *Cd248* | -0.176777 | -0.176777 | -0.176777 | -0.176777 | -0.176777 | -0.176777 | -0.176777 | -0.176777 | -0.176777 | 5.48008 | -0.176777 | -0.176777 | -0.176777 | -0.176777 | -0.176777 | -0.176777 | -0.176777 | -0.176777 | -0.176777 | -0.176777 | -0.176777 | -0.176777 | -0.176777 | -0.176777 | -0.176777 | -0.176777 | -0.176777 | -0.176777 | -0.176777 | -0.176777 | -0.176777 | -0.176777 | 2 | 2.4 | 2.4 | 81.812 | 3.0036 | 74329000 | 3 | 0.433155 | 0.610762 | -0.707107 | -0.930949 |
| Q91VA7;V9GXV0 | Q91VA7 | Isocitrate dehydrogenase [NAD] subunit, mitochondrial | *Idh3b* | -0.307357 | -0.307357 | -0.307357 | -0.307357 | -0.307357 | -0.307357 | 0.831002 | -0.307357 | -0.307357 | -0.307357 | 0.142829 | -0.307357 | -0.307357 | -0.307357 | -0.307357 | 1.33849 | 0.989544 | -0.307357 | -0.307357 | -0.307357 | -0.307357 | -0.307357 | -0.307357 | -0.307357 | -0.307357 | 4.99676 | -0.307357 | -0.307357 | -0.307357 | -0.307357 | -0.307357 | -0.307357 | 20 | 48.4 | 48.4 | 42.194 | 323.31 | 8456400000 | 71 | 0.920452 | 0.444441 | -0.424117 | -1.66358 |
| Q91VC9;A0A286YDK2 | Q91VC9;A0A286YDK2 | Growth hormone-inducible transmembrane protein | *Ghitm* | 0.667597 | 0.872172 | 0.122494 | -0.903231 | -0.903231 | 0.950442 | 1.14304 | -0.903231 | -0.903231 | -0.903231 | 1.09856 | 2.26464 | 1.87323 | -0.903231 | -0.903231 | -0.0300308 | -0.0765782 | -0.174593 | 0.489111 | 0.486198 | 0.722301 | -0.903231 | 0.329141 | 1.84915 | -0.903231 | -0.903231 | -0.903231 | -0.903231 | 0.961577 | -0.903231 | -0.903231 | -0.903231 | 2 | 6.1 | 6.1 | 37.275 | 6.4421 | 987030000 | 5 | 0.0617692 | 0.937999 | 0.097208 | 0.170265 |
| Q91VD9;A0A087WQR0;A0A087WP77;A0A087WSU3;A0A087W | Q91VD9 | NADH-ubiquinone oxidoreductase 75 kDa subunit, mitocho | *Ndufs1* | 3.54497 | -0.314619 | -0.314619 | -0.314619 | -0.314619 | -0.314619 | -0.314619 | -0.314619 | -0.314619 | -0.314619 | -0.314619 | -0.314619 | -0.314619 | -0.314619 | -0.314619 | -0.314619 | -0.314619 | -0.314619 | -0.314619 | -0.314619 | -0.314619 | -0.314619 | -0.314619 | -0.314619 | -0.314619 | -0.314619 | -0.314619 | -0.314619 | 2.78573 | 2.79325 | -0.314619 | -0.314619 | 30 | 41.1 | 41.1 | 79.776 | 305.03 | 12353000000 | 75 | 0 | 0.862667 | 0 | 0 |
| Q91VE0 | Q91VE0 | Long-chain fatty acid transport protein 4 | *Slc27a4* | -0.25351 | -0.25351 | -0.25351 | -0.25351 | -0.25351 | -0.25351 | -0.25351 | -0.25351 | -0.25351 | -0.25351 | -0.25351 | -0.25351 | -0.25351 | -0.25351 | -0.25351 | 3.52729 | 4.07802 | -0.25351 | -0.25351 | -0.25351 | -0.25351 | -0.25351 | -0.25351 | -0.25351 | -0.25351 | -0.25351 | -0.25351 | -0.25351 | -0.25351 | -0.25351 | -0.25351 | -0.25351 | 8 | 16 | 16 | 72.318 | 10.552 | 152280000 | 8 | 0.74521 | 0.511319 | -1.01404 | -1.4177 |
| Q91VI7;A0A1B0GSG5;A0A1B0GRG4;A0A1B0GRY7 | Q91VI7;A0A1B0GSG5;A0A1B0GRG4 | Ribonuclease inhibitor | *Rnh1* | -0.176777 | -0.176777 | -0.176777 | -0.176777 | -0.176777 | -0.176777 | -0.176777 | -0.176777 | -0.176777 | -0.176777 | -0.176777 | 5.48008 | -0.176777 | -0.176777 | -0.176777 | -0.176777 | -0.176777 | -0.176777 | -0.176777 | -0.176777 | -0.176777 | -0.176777 | -0.176777 | -0.176777 | -0.176777 | -0.176777 | -0.176777 | -0.176777 | -0.176777 | -0.176777 | -0.176777 | -0.176777 | 20 | 55.5 | 55.5 | 49.816 | 323.31 | 16021000000 | 82 | 0.433155 | 0.610247 | -0.707107 | -0.930949 |
| Q91VR5;A0A1Y7VM48 | Q91VR5 | ATP-dependent RNA helicase DDX1 | *Ddx1* | -0.579144 | -0.579144 | -0.374771 | -0.579144 | -0.579144 | 1.5929 | 2.7056 | -0.579144 | -0.579144 | -0.579144 | -0.579144 | 1.78049 | -0.579144 | 0.402342 | -0.579144 | -0.317832 | 2.40298 | -0.579144 | -0.579144 | -0.579144 | -0.579144 | -0.579144 | -0.579144 | 0.0125192 | -0.579144 | -0.379193 | 1.64714 | 1.08587 | -0.579144 | -0.579144 | 1.02483 | -0.579144 | 32 | 51.9 | 51.9 | 82.499 | 323.31 | 8684600000 | 85 | 0.872366 | 0.482523 | -0.738547 | -1.59751 |
| Q91VR7 | Q91VR7 | Microtubule-associated proteins 1A/1B light chain 3A | *Map1lc3a* | -0.176777 | -0.176777 | -0.176777 | -0.176777 | -0.176777 | -0.176777 | 5.48008 | -0.176777 | -0.176777 | -0.176777 | -0.176777 | -0.176777 | -0.176777 | -0.176777 | -0.176777 | -0.176777 | -0.176777 | -0.176777 | -0.176777 | -0.176777 | -0.176777 | -0.176777 | -0.176777 | -0.176777 | -0.176777 | -0.176777 | -0.176777 | -0.176777 | -0.176777 | -0.176777 | -0.176777 | -0.176777 | 4 | 29.8 | 22.3 | 14.272 | 10.807 | 455110000 | 7 | 0 | 0.86224 | 0 | 0 |
| Q91VU7;Q91VU7-2;F6XMS4;F7CG42 | Q91VU7;Q91VU7-2 | | *Pus7* | -0.176777 | -0.176777 | -0.176777 | -0.176777 | -0.176777 | -0.176777 | -0.176777 | -0.176777 | -0.176777 | -0.176777 | -0.176777 | -0.176777 | -0.176777 | -0.176777 | -0.176777 | -0.176777 | -0.176777 | -0.176777 | -0.176777 | -0.176777 | -0.176777 | -0.176777 | 5.48008 | -0.176777 | -0.176777 | -0.176777 | -0.176777 | -0.176777 | -0.176777 | -0.176777 | -0.176777 | -0.176777 | 4 | 8 | 8 | 74.792 | 4.6987 | 172920000 | 5 | 0.520085 | 0.56935 | 0.808122 | 1.07497 |
| Q91W43 | Q91W43 | Glycine dehydrogenase (decarboxylating), mitochondrial | *Gldc* | -0.224505 | -0.171517 | -0.454179 | -0.454179 | 0.976291 | -0.454179 | -0.454179 | -0.454179 | -0.454179 | -0.454179 | -0.454179 | -0.0687757 | -0.164656 | -0.454179 | 0.341239 | -0.402561 | -0.454179 | -0.454179 | -0.454179 | -0.454179 | -0.133899 | -0.114794 | -0.454179 | -0.454179 | -0.454179 | -0.454179 | 1.99673 | -0.454179 | 3.5811 | 2.98229 | -0.454179 | 0.0324562 | 22 | 29 | 29 | 113.27 | 323.31 | 6279300000 | 70 | 0.347853 | 0.805272 | -0.0960075 | -0.780813 |
| Q91WD5;D3YXT0;A0A0A6YW30;A0A0A6YXD3 | Q91WD5;D3YXT0 | NADH dehydrogenase [ubiquinone] iron-sulfur protein 2, mi | *Ndufs2* | 0.863138 | 0.40408 | 0.463549 | -0.792415 | 2.31917 | 1.81972 | -0.792415 | -0.792415 | -0.792415 | -0.792415 | -0.792415 | 1.55816 | 0.564778 | -0.792415 | -0.792415 | -0.792415 | -0.792415 | -0.792415 | -0.792415 | -0.792415 | -0.792415 | 0.210418 | 2.13633 | -0.792415 | -0.792415 | -0.792415 | 0.877078 | -0.285853 | 0.39044 | 0.254092 | 1.33003 | 0.565914 | 14 | 34.1 | 34.1 | 52.625 | 323.31 | 4179900000 | 51 | 0.0692546 | 0.934917 | 0.0981837 | 0.189536 |
| Q91WK2 | Q91WK2 | Eukaryotic translation initiation factor 3 subunit H | *Eif3h* | -0.21224 | -0.21224 | -0.21224 | -0.21224 | 5.34901 | -0.21224 | -0.21224 | -0.21224 | -0.21224 | -0.21224 | 1.01818 | -0.21224 | -0.21224 | -0.21224 | -0.21224 | -0.21224 | -0.21224 | -0.21224 | -0.21224 | -0.21224 | -0.21224 | -0.21224 | -0.21224 | -0.21224 | -0.21224 | -0.21224 | -0.21224 | -0.21224 | -0.21224 | -0.21224 | -0.21224 | -0.21224 | 10 | 36.9 | 36.9 | 39.832 | 323.31 | 4615600000 | 31 | 0.433155 | 0.609732 | -0.153802 | -0.930949 |
| Q91WM2 | Q91WM2 | Cat eye syndrome critical region protein 5 homolog | *Cecr5* | -0.254128 | -0.254128 | -0.254128 | -0.254128 | -0.254128 | -0.254128 | -0.254128 | -0.254128 | 3.83491 | -0.254128 | -0.254128 | -0.254128 | -0.254128 | -0.254128 | -0.254128 | 3.78893 | -0.254128 | -0.254128 | -0.254128 | -0.254128 | -0.254128 | -0.254128 | -0.254128 | -0.254128 | -0.254128 | -0.254128 | -0.254128 | -0.254128 | -0.254128 | -0.254128 | -0.254128 | -0.254128 | 5 | 12.9 | 12.9 | 46.306 | 56.288 | 684360000 | 11 | 0.433155 | 0.609218 | -0.505383 | -0.930949 |
| Q91WS0 | Q91WS0 | CDGSH iron-sulfur domain-containing protein 1 | *Cisd1* | -0.176777 | -0.176777 | -0.176777 | -0.176777 | -0.176777 | 5.48008 | -0.176777 | -0.176777 | -0.176777 | -0.176777 | -0.176777 | -0.176777 | -0.176777 | -0.176777 | -0.176777 | -0.176777 | -0.176777 | -0.176777 | -0.176777 | -0.176777 | -0.176777 | -0.176777 | -0.176777 | -0.176777 | -0.176777 | -0.176777 | -0.176777 | -0.176777 | -0.176777 | -0.176777 | -0.176777 | -0.176777 | 7 | 52.8 | 52.8 | 12.097 | 60.503 | 3127800000 | 25 | 0 | 0.861813 | 0 | 0 |
| Q91XD6 | Q91XD6 | Vacuolar protein-sorting-associated protein 36 | *Vps36* | -0.398707 | -0.398707 | -0.398707 | -0.398707 | -0.398707 | -0.398707 | -0.398707 | -0.398707 | -0.398707 | -0.398707 | -0.398707 | 1.41554 | -0.398707 | -0.398707 | -0.398707 | -0.398707 | 3.10654 | -0.398707 | -0.398707 | -0.398707 | -0.398707 | -0.398707 | -0.398707 | -0.398707 | 2.2294 | 3.01398 | -0.398707 | -0.398707 | -0.398707 | -0.398707 | -0.398707 | 0.999636 | 13 | 36.5 | 36.5 | 43.735 | 44.033 | 1223000000 | 21 | 0.688672 | 0.526856 | -0.664936 | -1.33496 |
| Q91XH5;G3UXX3;Q64105;G3UZ79 | Q91XH5;G3UXX3;Q64105 | Sepiapterin reductase | *Spr* | -0.252091 | -0.252091 | -0.252091 | -0.252091 | -0.252091 | -0.252091 | -0.252091 | -0.252091 | 0.424211 | -0.252091 | -0.252091 | -0.252091 | 1.82061 | -0.252091 | -0.252091 | -0.252091 | -0.252091 | -0.252091 | -0.252091 | -0.252091 | -0.252091 | -0.252091 | -0.252091 | -0.252091 | -0.252091 | -0.252091 | -0.252091 | -0.252091 | 5.06581 | -0.252091 | -0.252091 | -0.252091 | 14 | 66.4 | 66.4 | 27.928 | 323.31 | 6813900000 | 54 | 0.433155 | 0.608704 | -0.259088 | -0.930949 |
| Q91Y44 | Q91Y44 | Bromodomain testis-specific protein | *Brdt* | -1.18747 | -1.18747 | -1.18747 | 0.0576654 | 1.51457 | 2.01047 | -1.18747 | -1.18747 | -0.162389 | 0.844097 | 1.05075 | 0.76119 | 0.789936 | 0.514258 | 0.811291 | -1.18747 | -1.18747 | -1.18747 | -0.17392 | -0.0500071 | 1.51433 | 0.717585 | -0.263403 | -0.147854 | -1.18747 | -1.18747 | 0.721889 | 0.52855 | 0.472521 | 0.881855 | 0.668784 | -1.18747 | 2 | 2.6 | 1.3 | 107.25 | 0.13068 | 9156500000 | 7 | 0.214243 | 0.849688 | -0.241107 | -0.521777 |
| Q91YH5;E9PYT3;Q91YH5-2 | Q91YH5;E9PYT3;Q91YH5-2 | Atlastin-3 | *Atl3* | -0.453785 | -0.453785 | -0.453785 | -0.453785 | 2.13419 | 3.21697 | -0.453785 | -0.453785 | -0.453785 | -0.453785 | -0.453785 | -0.453785 | -0.453785 | -0.453785 | -0.453785 | -0.453785 | -0.453785 | -0.453785 | -0.453785 | -0.453785 | 1.86088 | 1.12606 | -0.453785 | -0.453785 | -0.453785 | -0.453785 | -0.453785 | -0.453785 | 1.61919 | 1.84111 | -0.453785 | -0.453785 | 13 | 34.2 | 34.2 | 60.574 | 323.31 | 4747600000 | 43 | 0.892521 | 0.46699 | 0.556359 | 1.62532 |
| Q91YJ2;A0A338P7D2 | Q91YJ2 | Sorting nexin-4 | *Snx4* | 0.510228 | 0.301091 | 0.909234 | 1.11692 | -0.847635 | -0.847635 | -0.847635 | -0.847635 | -0.564547 | -0.847635 | -0.847635 | 0.804791 | 0.753545 | -0.847635 | -0.847635 | -0.847635 | -0.847635 | -0.847635 | -0.847635 | -0.847635 | 0.907833 | 1.14374 | -0.847635 | -0.847635 | -0.847635 | -0.847635 | 1.84337 | 1.56252 | 1.6496 | 1.41854 | 1.10971 | 0.943215 | 9 | 22.4 | 22.4 | 51.777 | 17.667 | 992640000 | 16 | 0.113419 | 0.904305 | 0.128563 | 0.298337 |
| Q91YN9;A0A087WNX9 | Q91YN9 | BAG family molecular chaperone regulator 2 | *Bag2* | -0.243212 | -0.243212 | -0.243212 | -0.243212 | -0.243212 | -0.243212 | -0.243212 | 4.78988 | -0.243212 | -0.243212 | -0.243212 | -0.243212 | -0.243212 | -0.243212 | -0.243212 | 2.50648 | -0.243212 | -0.243212 | -0.243212 | -0.243212 | -0.243212 | -0.243212 | -0.243212 | -0.243212 | -0.243212 | -0.243212 | -0.243212 | -0.243212 | -0.243212 | -0.243212 | -0.243212 | -0.243212 | 3 | 20 | 20 | 23.473 | 14.369 | 33655000 | 3 | 0.433155 | 0.608192 | -0.343712 | -0.930949 |
| Q91YQ5;A0A0N4SUJ8 | Q91YQ5 | Dolichyl-diphosphooligosaccharide--protein glycosyltransfe | *Rpn1* | 0.69127 | 0.768433 | 1.32234 | 1.09963 | 0.755488 | 0.611897 | -0.5559 | -1.03959 | 2.55025 | -1.03959 | -1.03959 | 0.273947 | 0.680914 | -0.957368 | -1.03959 | -1.03959 | -1.03959 | -1.03959 | -1.03959 | -1.03959 | 0.866812 | -1.03959 | -0.0907766 | -0.0563627 | -1.03959 | -1.03959 | 0.720569 | 0.213012 | 0.986219 | 1.25376 | 0.772411 | 0.568517 | 28 | 53.1 | 53.1 | 68.527 | 323.31 | 18495000000 | 95 | 0.167528 | 0.875413 | 0.158816 | 0.421969 |
| Q91YR1;D3Z2H0 | Q91YR1 | Twinfilin-1 | *Twf1* | -0.342504 | -0.342504 | -0.342504 | -0.342504 | -0.342504 | -0.342504 | -0.342504 | -0.342504 | -0.342504 | -0.342504 | -0.342504 | -0.342504 | 2.53059 | 3.73289 | -0.342504 | -0.342504 | 0.703189 | -0.342504 | -0.342504 | -0.342504 | -0.342504 | -0.342504 | -0.342504 | -0.342504 | -0.342504 | -0.342504 | -0.342504 | -0.342504 | -0.342504 | -0.342504 | 2.62346 | -0.342504 | 15 | 44 | 38 | 40.079 | 323.31 | 6437000000 | 41 | 0.90487 | 0.457188 | -0.999272 | -1.64228 |
| Q91YR7;Q91YR7-2 | Q91YR7 | Pre-mRNA-processing factor 6 | *Prpf6* | -0.252311 | -0.252311 | -0.252311 | -0.252311 | 4.25522 | -0.252311 | -0.252311 | -0.252311 | -0.252311 | -0.252311 | -0.252311 | -0.252311 | -0.252311 | -0.252311 | -0.252311 | -0.252311 | -0.252311 | -0.252311 | -0.252311 | -0.252311 | -0.252311 | -0.252311 | -0.252311 | -0.252311 | -0.252311 | -0.252311 | 3.3141 | -0.252311 | -0.252311 | -0.252311 | -0.252311 | -0.252311 | 5 | 5.6 | 5.6 | 106.72 | 3.7842 | 179500000 | 9 | 0 | 0.861386 | 0 | 0 |

[illegible]

| | | | | | |
|---|---|---|---|---|---|
| Q9D3B1 | Q9D3B1 | Very-long-chain (3R)-3-hydroxyacyl-CoA dehydratase 2 | *Hacd2* | -0.252297 | [illegible] |
| Q9D3D9 | Q9D3D9 | ATP synthase subunit delta, mitochondrial | *Atp5d* | -0.176777 | [illegible] |
| Q9D4H1 | Q9D4H1 | Exocyst complex component 2 | *Exoc2* | -0.176777 | [illegible] |
| Q9D4H2;F6VDD7 | Q9D4H2;F6VDD7 | GRIP and coiled-coil domain-containing protein 1 | *Gcc1* | 2.50123 | [illegible] |
| Q9D554 | Q9D554 | Splicing factor 3A subunit 3 | *Sf3a3* | -0.639643 | [illegible] |
| Q9D5T0 | Q9D5T0 | ATPase family AAA domain-containing protein 1 | *Atad1* | -0.240741 | [illegible] |
| Q9D600 | Q9D600 | DNA replication complex GINS protein PSF2 | *Gins2* | -0.404558 | [illegible] |
| Q9D662;A2ANA0;A2AN97;A2AN98;A2AN99 | Q9D662 | Protein transport protein Sec23B | *Sec23b* | -0.176777 | [illegible] |
| Q9D6F9 | Q9D6F9 | Tubulin beta-4A chain | *Tubb4a* | -0.269253 | [illegible] |
| Q9D6J1;H3BL50 | Q9D6J1;H3BL50 | Ceramide synthase 4 | *Cers4* | -0.176777 | [illegible] |
| Q9D6J6;Q9D6J6-2;M0QWP9;D6RGN4 | Q9D6J6;Q9D6J6-2;M0QWP9 | NADH dehydrogenase [ubiquinone] flavoprotein 2, mitocho | *Ndufv2* | -0.176777 | [illegible] |
| Q9D6K5;E9PVN6;Q9D6K5-3;Q9D6K5-2;Q9D6K5-4 | Q9D6K5;E9PVN6;Q9D6K5-3;Q9D6K5-2;Q9 | Synaptojanin-2-binding protein | *Synj2bp;Gm20498* | -0.334396 | [illegible] |
| Q9D6M3;E9Q6M6;E9PV90;E9PY45;V9GWS2;Q80X52;E9Q579; | Q9D6M3;E9Q6M6;E9PV90;E9PY45 | Mitochondrial glutamate carrier 1 | *Slc25a22* | -0.176777 | [illegible] |
| Q9D6U8;A0A338P6X2 | Q9D6U8 | Protein FAM162A | *Fam162a* | -0.176777 | [illegible] |
| Q9D708;D3Z2Y6 | Q9D708;D3Z2Y6 | | *S100a16* | -0.30787 | [illegible] |
| Q9D7N3;A0A0J9YU02 | Q9D7N3 | 28S ribosomal protein S9, mitochondrial | *Mrps9* | -0.176777 | [illegible] |
| Q9D7N9 | Q9D7N9 | Adipocyte plasma membrane-associated protein | *Apmap* | -0.468057 | [illegible] |
| Q9D7P6;D3Z0K9;D3Z7W0 | Q9D7P6;D3Z0K9;D3Z7W0 | Iron-sulfur cluster assembly enzyme ISCU, mitochondrial | *Iscu* | -0.176777 | [illegible] |
| Q9D7S7-2;Q9D7S7 | Q9D7S7-2;Q9D7S7 | 60S ribosomal protein L22-like 1 | *Rpl22l1* | -0.176777 | [illegible] |
| Q9D7S9 | Q9D7S9 | Charged multivesicular body protein 5 | *Chmp5* | -0.331152 | [illegible] |
| Q9D883;G3UW94;E9PWA1;A0A140LII6;E9Q5Y2;E9PUZ3;E9Q1 | Q9D883;G3UW94 | Splicing factor U2AF 35 kDa subunit | *U2af1* | -0.369466 | [illegible] |
| Q9D8E6 | Q9D8E6 | 60S ribosomal protein L4 | *Rpl4* | -0.527302 | [illegible] |
| Q9D8S9 | Q9D8S9 | BolA-like protein 1 | *Bola1* | -0.176777 | [illegible] |
| Q9D8T7;F8WHU8;Q9D8T7-2;F7DE82 | Q9D8T7;F8WHU8;Q9D8T7-2 | SRA stem-loop-interacting RNA-binding protein, mitochondr | *Slirp* | -0.434473 | [illegible] |
| Q9D8W5;B1AT36;Q3TRH2 | Q9D8W5;B1AT36;Q3TRH2 | 26S proteasome non-ATPase regulatory subunit 12 | *Psmd12* | 1.18386 | [illegible] |
| Q9D8W7;A0A0J9YUI7;A0A0J9YU56;A0A0J9YU93 | Q9D8W7;A0A0J9YUI7;A0A0J9YU56 | OCIA domain-containing protein 2 | *Ociad2* | -0.23006 | [illegible] |
| Q9DAU1;Q9DAU1-2;B0V2V1 | Q9DAU1 | Protein canopy homolog 3 | *Cnpy3* | -0.639471 | [illegible] |
| Q9DAW9;A0A0G2JDV8;A0A0G2JFI6;A0A0G2JFC8;A0A0G2JF | Q9DAW9;A0A0G2JDV8 | Calponin-3 | *Cnn3* | -0.351545 | [illegible] |
| Q9DB05;A0A1B0GR35;P28663 | Q9DB05 | Alpha-soluble NSF attachment protein | *Napa* | -0.315567 | [illegible] |
| Q9DB10;A0A2R8VHK6 | Q9DB10;A0A2R8VHK6 | Essential MCU regulator, mitochondrial | *Smdt1* | 1.32138 | [illegible] |
| Q9DB41-2;Q9DB41 | Q9DB41-2;Q9DB41 | Mitochondrial glutamate carrier 2 | *Slc25a18* | -0.294424 | [illegible] |
| Q9DB77;A0A140LI98 | Q9DB77 | Cytochrome b-c1 complex subunit 2, mitochondrial | *Uqcrc2* | -0.413093 | [illegible] |
| Q9DBF1-2;Q9DBF1;G3UYR8;G3UY72;E9Q1H3;E9Q1G1 | Q9DBF1-2;Q9DBF1;G3UYR8;G3UY72 | Alpha-aminoadipic semialdehyde dehydrogenase | *Aldh7a1* | 3.06043 | [illegible] |
| Q9DBG3;Q9DBG3-2;H3BIY9;H3BKM0;H3BJ06 | Q9DBG3;Q9DBG3-2;H3BIY9;H3BKM0 | AP-2 complex subunit beta;AP complex subunit beta | *Ap2b1* | -0.195966 | [illegible] |
| Q9DBG7;F6VF36 | Q9DBG7;F6VF36 | Signal recognition particle receptor subunit alpha | *Srpr* | -0.176777 | [illegible] |
| Q9DBH5 | Q9DBH5 | Vesicular integral-membrane protein VIP36 | *Lman2* | 5.15971 | [illegible] |
| Q9DBL1;E9Q5L3;A0A140LHL4 | Q9DBL1;E9Q5L3 | Short/branched chain specific acyl-CoA dehydrogenase, m | *Acadsb* | -0.403542 | [illegible] |
| Q9DBP5;A0A0R4J093 | Q9DBP5;A0A0R4J093 | UMP-CMP kinase | *Cmpk1* | -0.192539 | [illegible] |
| Q9DBR7;Q9DBR7-2;A0A1W2P750 | Q9DBR7;Q9DBR7-2;A0A1W2P750 | Protein phosphatase 1 regulatory subunit 12A | *Ppp1r12a* | 0.740095 | [illegible] |
| Q9DBS1 | Q9DBS1 | Transmembrane protein 43 | *Tmem43* | -0.176777 | [illegible] |
| Q9DBS5 | Q9DBS5 | Kinesin light chain 4 | *Klc4* | -0.176777 | [illegible] |
| Q9DBV4;A2AD97 | Q9DBV4;A2AD97 | Matrix-remodeling-associated protein 8 | *Mxra8* | 1.1167 | [illegible] |
| Q9DBZ5;A0A140LJ59;Q3TY56;Q9DBZ5-2 | Q9DBZ5;A0A140LJ59;Q3TY56;Q9DBZ5-2 | Eukaryotic translation initiation factor 3 subunit K | *Eif3k* | -0.176777 | [illegible] |
| Q9DC07;F6RM14;B7ZCI2;F6X9B6;A2AQB2;A2AQA9;E9Q1W3 | Q9DC07;F6RM14 | LIM zinc-binding domain-containing Nebulette | *Nebl* | -0.20116 | [illegible] |
| Q9DC11;B1AY85;B1AY86 | Q9DC11;B1AY85;B1AY86 | Plexin domain-containing protein 2 | *Plxdc2* | 1.21651 | [illegible] |
| Q9DC50 | Q9DC50 | Peroxisomal carnitine O-octanoyltransferase | *Crot* | -0.25393 | [illegible] |
| Q9DC70 | Q9DC70 | NADH dehydrogenase [ubiquinone] iron-sulfur protein 7, mi | *Ndufs7* | -0.312572 | [illegible] |
| Q9DCC8;A0A1D5RLZ6 | Q9DCC8;A0A1D5RLZ6 | Mitochondrial import receptor subunit TOM20 homolog | *Tomm20* | -0.254085 | [illegible] |
| Q9DCH2 | Q9DCH2 | Ribonuclease P protein subunit p20 | *Pop7* | 0.797853 | [illegible] |
| Q9DCN2-2;Q9DCN2;F2Z456;A0A0A0MQM3 | Q9DCN2-2;Q9DCN2;F2Z456;A0A0A0MQM | NADH-cytochrome b5 reductase 3;NADH-cytochrome b5 r | *Cyb5r3* | -0.176777 | [illegible] |
| Q9DCP2;A0A0A6YWF5;A0A0A6YX60;A0A0A6YWH4 | Q9DCP2;A0A0A6YWF5;A0A0A6YX60;A0A0 | Sodium-coupled neutral amino acid transporter 3 | *Slc38a3* | -0.229419 | [illegible] |
| Q9DCS2;G5E8X1;E9Q5B2;E9Q7K5;D3YZM4 | Q9DCS2;G5E8X1;E9Q5B2;E9Q7K5 | UPF0585 protein C16orf13 homolog | *0610011F06Rik* | -0.222482 | [illegible] |
| Q9DCS3;A2A845 | Q9DCS3;A2A845 | Trans-2-enoyl-CoA reductase, mitochondrial | *Mecr* | -0.176777 | [illegible] |
| Q9DCT1;Q8CEB6 | Q9DCT1;Q8CEB6 | 1,5-anhydro-D-fructose reductase | *Akr1e2;Akr1e1* | -0.176777 | [illegible] |
| Q9DCT2 | Q9DCT2 | NADH dehydrogenase [ubiquinone] iron-sulfur protein 3, mi | *Ndufs3* | -0.176777 | [illegible] |
| Q9DCT8;A0A0G2JF37 | Q9DCT8 | Cysteine-rich protein 2 | *Crip2* | -0.176777 | [illegible] |
| Q9DCW4;A0A0U1RNP5;A0A0N4SVE0;A0A0U1RNR3;A0A0N4S | Q9DCW4;A0A0U1RNP5;A0A0N4SVE0;A0A | Electron transfer flavoprotein subunit beta | *Etfb* | -0.364176 | [illegible] |
| Q9DCX2;B1ASE2 | Q9DCX2;B1ASE2 | ATP synthase subunit d, mitochondrial | *Atp5h* | -0.263991 | [illegible] |
| Q9DCZ1;F6VY18 | Q9DCZ1;F6VY18 | GMP reductase 1 | *Gmpr* | -0.305183 | [illegible] |
| Q9DCZ4-2;Q9DCZ4;Q9DCZ4-3 | Q9DCZ4-2;Q9DCZ4;Q9DCZ4-3 | Apolipoprotein O | *Apoo* | -0.46173 | [illegible] |
| Q9EP69;A0A1L1SR59 | Q9EP69 | Phosphatidylinositide phosphatase SAC1 | *Sacm1l* | -0.223876 | [illegible] |
| Q9EP96;E9Q0W2;L7N264;Q9QXZ6;E9PWV5;L7N251;M0QWR | Q9EP96 | Solute carrier organic anion transporter family member 1A4 | *Slco1a4* | -0.570932 | [illegible] |
| Q9EPK2-3;Q9EPK2;Q9EPK2-2;Q9EPK2-4 | Q9EPK2-3;Q9EPK2;Q9EPK2-2;Q9EPK2-4 | Protein XRP2 | *Rp2* | -0.285969 | [illegible] |
| Q9EPL8;A0A140LHD0 | Q9EPL8 | Importin-7 | *Ipo7* | -0.492385 | [illegible] |
| Q9EPU0-2;Q9EPU0;Q6ZQJ5 | Q9EPU0-2;Q9EPU0 | Regulator of nonsense transcripts 1 | *Upf1* | -0.200972 | [illegible] |
| Q9EQ20;A0A1Y7VJ32 | Q9EQ20 | Methylmalonate-semialdehyde dehydrogenase [acylating], | *Aldh6a1* | -0.176777 | [illegible] |
| Q9EQ80;D3Z0G0;E9PWJ6 | Q9EQ80;D3Z0G0 | NIF3-like protein 1 | *Nif3l1* | -0.252021 | [illegible] |
| Q9EQF6;Q3SYJ1 | Q9EQF6 | Dihydropyrimidinase-related protein 5 | *Dpysl5* | -0.176777 | [illegible] |
| Q9EQG9-2;Q9EQG9;Q9EQG9-3 | Q9EQG9-2;Q9EQG9;Q9EQG9-3 | Collagen type IV alpha-3-binding protein | *Col4a3bp* | 1.96867 | [illegible] |
| Q9EQH3 | Q9EQH3 | Vacuolar protein sorting-associated protein 35 | *Vps35* | -0.186687 | [illegible] |
| Q9EQP2 | Q9EQP2 | EH domain-containing protein 4 | *Ehd4* | -0.319592 | [illegible] |
| Q9EQQ2 | Q9EQQ2 | Protein YIPF5 | *Yipf5* | -0.512741 | [illegible] |
| Q9EQU5-2;A2BE93;Q9EQU5;A2BE92;Q9EQU5-3 | Q9EQU5-2;A2BE93;Q9EQU5;A2BE92 | Protein SET | *Set* | -0.304481 | [illegible] |
| Q9ER00 | Q9ER00 | Syntaxin-12 | *Stx12* | -0.250667 | [illegible] |
| Q9ER65-2;Q9ER65 | Q9ER65-2;Q9ER65 | Calsyntenin-2 | *Clstn2* | -0.0880686 | [illegible] |
| Q9ERG0;Q9ERG0-2;A0A2R8VI67;Q4U4S6 | Q9ERG0;Q9ERG0-2 | LIM domain and actin-binding protein 1 | *Lima1* | -0.176777 | [illegible] |
| Q9ERK4;E9Q1T9;F6ZEW4;E9QAX7 | Q9ERK4;E9Q1T9;F6ZEW4;E9QAX7 | Exportin-2 | *Cse1l* | -0.447357 | [illegible] |
| Q9ERS2 | Q9ERS2 | NADH dehydrogenase [ubiquinone] 1 alpha subcomplex su | *Ndufa13* | -0.45568 | [illegible] |
| Q9ERU9 | Q9ERU9 | E3 SUMO-protein ligase RanBP2 | *Ranbp2* | -0.532312 | [illegible] |
| Q9ES00;G3UZD6;G3UZM6;H7BXC2 | Q9ES00;G3UZD6 | Ubiquitin conjugation factor E4 B | *Ube4b* | -0.315647 | [illegible] |
| Q9ES74;Q3TN15 | Q9ES74;Q3TN15 | Serine/threonine-protein kinase Nek7 | *Nek7* | 2.13719 | [illegible] |
| Q9ESQ8-2;Q9ESQ8 | Q9ESQ8-2;Q9ESQ8 | Pro-FMRFamide-related neuropeptide VF;Neuropeptide NF | *Npvf* | -0.315409 | [illegible] |
| Q9ESW8;A0A1B0GSC4 | Q9ESW8;A0A1B0GSC4 | Pyroglutamyl-peptidase 1 | *Pgpep1* | -0.176777 | [illegible] |
| Q9ET01;Q3UEJ6 | Q9ET01;Q3UEJ6 | Glycogen phosphorylase, liver form;Alpha-1,4 glucan phos | *Pygl* | -0.176777 | [illegible] |
| Q9ET54-4;Q9ET54-6;Q9ET54-3;Q9ET54-2;D3Z1J5 | Q9ET54-4;Q9ET54-6;Q9ET54-3;Q9ET54-2 | | | -0.43993 | [illegible] |
| Q9JHF7 | Q9JHF7 | Hematopoietic prostaglandin D synthase | *Hpgds* | -0.176777 | [illegible] |
| Q9JHH9;F2Z4A2 | Q9JHH9 | Coatomer subunit zeta-2 | *Copz2* | -0.322205 | [illegible] |
| Q9JHJ0;A0A1L1SQ12;A0A1L1SR42 | Q9JHJ0 | Tropomodulin-3 | *Tmod3* | -0.326775 | [illegible] |
| Q9JHQ5;A0A1L1SSX7;F6VG18;F6VTH5;A0A1L1ST13;E9Q9P1 | Q9JHQ5;A0A1L1SSX7;F6VG18;F6VTH5 | Leucine zipper transcription factor-like protein 1 | *Lztfl1* | -0.252864 | [illegible] |
| Q9JHS3;D3YTS4 | Q9JHS3;D3YTS4 | Ragulator complex protein LAMTOR2 | *Lamtor2* | -0.176777 | [illegible] |
| Q9JI91 | Q9JI91 | Alpha-actinin-2 | *Actn2* | -0.244958 | [illegible] |
| Q9JIF7 | Q9JIF7 | Coatomer subunit beta | *Copb1* | -0.393092 | [illegible] |
| Q9JIG7 | Q9JIG7 | Coiled-coil domain-containing protein 22 | *Ccdc22* | 0.38726 | [illegible] |
| Q9JIG8;A0A140LJ36 | Q9JIG8;A0A140LJ36 | PRA1 family protein 2 | *Praf2* | -0.382409 | [illegible] |
| Q9JII6;B1AXW3 | Q9JII6;B1AXW3 | Alcohol dehydrogenase [NADP(+)] | *Akr1a1* | -0.176777 | [illegible] |
| Q9JIK9 | Q9JIK9 | 28S ribosomal protein S34, mitochondrial | *Mrps34* | -0.410679 | [illegible] |
| Q9JJ28 | Q9JJ28 | Protein flightless-1 homolog | *Flii* | -0.528112 | [illegible] |
| Q9JJ78;D3YVF2 | Q9JJ78;D3YVF2 | Lymphokine-activated killer T-cell-originated protein kinase | *Pbk* | -0.486904 | [illegible] |
| Q9JJU8 | Q9JJU8 | SH3 domain-binding glutamic acid-rich-like protein | *Sh3bgrl* | -0.796378 | [illegible] |
| Q9JJV2-3;Q9JJV2;D3YWS3;Q9JJV2-2 | Q9JJV2-3;Q9JJV2;D3YWS3;Q9JJV2-2 | Profilin-2;Profilin | *Pfn2* | -0.419078 | [illegible] |
| Q9JK48-3;Q9JK48;A0A0G2JEC4;Q9JK48-2;A0A0G2JE45;A0A | Q9JK48-3;Q9JK48;A0A0G2JEC4;Q9JK48-2 | Endophilin-B1 | *Sh3glb1* | -0.176777 | [illegible] |
| Q9JK81;F8WGG3;F7A3N3 | Q9JK81;F8WGG3 | UPF0160 protein MYG1, mitochondrial | *Myg1* | -0.389575 | [illegible] |
| Q9JK92 | Q9JK92 | Heat shock protein beta-8 | *Hspb8* | -0.176777 | [illegible] |
| Q9JKB1;A0A2I3BQ39;P58321 | Q9JKB1;A0A2I3BQ39;P58321 | Ubiquitin carboxyl-terminal hydrolase isozyme L3;Ubiquitin | *Uchl3;Uchl4* | 0.972257 | [illegible] |
| Q9JKB3-2;Q9JKB3 | Q9JKB3-2;Q9JKB3 | Y-box-binding protein 3 | *Ybx3* | -0.23092 | [illegible] |
| Q9JKF1;A0A0U1RNG5;A0A0U1RPI2;A0A0U1RPU3 | Q9JKF1 | Ras GTPase-activating-like protein IQGAP1 | *Iqgap1* | -0.203814 | [illegible] |
| Q9JKR6;A0A1L1SQ34;E0CYZ2;F6TRP3 | Q9JKR6 | Hypoxia up-regulated protein 1 | *Hyou1* | 1.08616 | [illegible] |
| Q9JKY5;A0A0G2JFU3;E9Q3C5;F7DCR9 | Q9JKY5 | Huntingtin-interacting protein 1-related protein | *Hip1r* | 5.48008 | [illegible] |
| Q9JKZ2 | Q9JKZ2 | Sodium/myo-inositol cotransporter | *Slc5a3* | 1.20318 | [illegible] |
| Q9JLI8;Q9JLI8-2 | Q9JLI8 | Squamous cell carcinoma antigen recognized by T-cells 3 | *Sart3* | 0.0646927 | [illegible] |
| Q9JLN9;Q9JLN9-2 | Q9JLN9 | Serine/threonine-protein kinase mTOR | *Mtor* | 0.755367 | [illegible] |
| Q9JLQ0;A0A3B2W812 | Q9JLQ0;A0A3B2W812 | CD2-associated protein | *Cd2ap* | -0.365243 | [illegible] |
| Q9JLV1 | Q9JLV1 | BAG family molecular chaperone regulator 3 | *Bag3* | 3.4234 | [illegible] |
| Q9JLV5;E9Q4T8;F6ZZK0;F6UY44;F6R0N8 | Q9JLV5;E9Q4T8 | Cullin-3 | *Cul3* | -0.684347 | [illegible] |
| Q9JM76;H7BWZ3;D3Z2F7;D3Z2F8;A0A0G2JFK7 | Q9JM76;H7BWZ3;D3Z2F7;D3Z2F8;A0A0G | Actin-related protein 2/3 complex subunit 3 | *Arpc3* | -0.358968 | [illegible] |
| Q9JMC3;D3Z1U5 | Q9JMC3;D3Z1U5 | DnaJ homolog subfamily A member 4 | *Dnaja4* | -0.176777 | [illegible] |
| Q9JMD0-4;Q9JMD0-3;Q9JMD0-2;Q9JMD0;M0QWF0;Q8CCB2 | Q9JMD0-4;Q9JMD0-3;Q9JMD0-2;Q9JMD | BUB3-interacting and GLEBS motif-containing protein ZNF2 | *Znf207;Zfp207* | -0.176777 | [illegible] |
| Q9JMD3;G3UYM0;G3UY59;G3UW37;G3UY87;E9PVP0 | Q9JMD3;G3UYM0;G3UY59;G3UW37;G3U | PCTP-like protein | *Stard10* | -0.247574 | [illegible] |
| Q9JMG7;Q9JMG7-2 | Q9JMG7;Q9JMG7-2 | Hepatoma-derived growth factor-related protein 3 | *Hdgfrp3* | -0.497477 | [illegible] |
| Q9JMH6-2;Q9JMH6;A0A1W2P6U1 | Q9JMH6-2;Q9JMH6 | Thioredoxin reductase 1, cytoplasmic | *Txnrd1* | -0.252665 | [illegible] |
| Q9QUI0;A0A0A6YXF6 | Q9QUI0;A0A0A6YXF6 | Transforming protein RhoA | *Rhoa* | -0.249007 | [illegible] |
| Q9QUN9 | Q9QUN9 | Dickkopf-related protein 3 | *Dkk3* | -0.457769 | [illegible] |
| Q9QUR7;Q3ULQ2 | Q9QUR7 | Peptidyl-prolyl cis-trans isomerase NIMA-interacting 1 | *Pin1* | 0.0715441 | [illegible] |
| Q9QXB9 | Q9QXB9 | Developmentally-regulated GTP-binding protein 2 | *Drg2* | -0.176777 | [illegible] |
| Q9QXK3-4;Q9QXK3;Q9QXK3-3;Q9QXK3-2 | Q9QXK3-4;Q9QXK3;Q9QXK3-3 | Coatomer subunit gamma-2 | *Copg2* | -0.176777 | [illegible] |
| Q9QXT0;A0A1W2P729 | Q9QXT0;A0A1W2P729 | Protein canopy homolog 2 | *Cnpy2* | 1.22645 | [illegible] |
| Q9QXV0 | Q9QXV0 | ProSAAS;KEP;Big SAAS;Little SAAS;Big PEN-LEN;PEN;PE | *Pcsk1n* | 1.34163 | [illegible] |
| Q9QXX4 | Q9QXX4 | Calcium-binding mitochondrial carrier protein Aralar2 | *Slc25a13* | 5.48008 | [illegible] |
| Q9QXY6 | Q9QXY6 | EH domain-containing protein 3 | *Ehd3* | -0.316373 | [illegible] |
| Q9R1R8;Q9QYF1 | Q9R1R8;Q9QYF1 | Retinol dehydrogenase 11 | *Rdh11* | 1.3426 | [illegible] |
| Q9QYG0-2;Q9QYG0;A0A2I3BPW0;A0A2I3BPL1;A0A2I3BQR5; | Q9QYG0-2;Q9QYG0;A0A2I3BPW0 | Protein NDRG2 | *Ndrg2* | 3.69551 | [illegible] |
| Q9QZB7;A0A1Y7VM21;A0A1Y7VL71 | Q9QZB7 | Actin-related protein 10 | *Actr10* | -0.176777 | [illegible] |
| Q9QZD8 | Q9QZD8 | Mitochondrial dicarboxylate carrier | *Slc25a10* | -0.176777 | [illegible] |
| Q9QZE5;Q7TNQ1 | Q9QZE5 | Coatomer subunit gamma-1 | *Copg1* | -0.176777 | [illegible] |
| Q9QZE7;A0A1D5RLR0 | Q9QZE7;A0A1D5RLR0 | Translin-associated protein X | *Tsnax* | 3.41022 | [illegible] |
| Q9QZF2 | Q9QZF2 | Glypican-1;Secreted glypican-1 | *Gpc1* | -0.44848 | [illegible] |
| Q9QZM0 | Q9QZM0 | Ubiquilin-2 | *Ubqln2* | -0.233142 | [illegible] |
| Q9R002 | Q9R002 | Interferon-activable protein 202 | *Ifi202* | -0.27696 | [illegible] |
| Q9R059;F2Z455 | Q9R059;F2Z455 | Four and a half LIM domains protein 3 | *Fhl3* | -0.234758 | [illegible] |
| Q9R060;A0A2R8W6X5 | Q9R060;A0A2R8W6X5 | Cytosolic Fe-S cluster assembly factor NUBP1 | *Nubp1* | -0.176777 | [illegible] |
| Q9R0E2;A8Y5E6;F6WNR1 | Q9R0E2 | Procollagen-lysine,2-oxoglutarate 5-dioxygenase 1 | *Plod1* | -0.176777 | [illegible] |
| Q9R0H0-2;Q9R0H0;A2A848 | Q9R0H0-2;Q9R0H0;A2A848 | Peroxisomal acyl-coenzyme A oxidase 1 | *Acox1* | -0.250365 | [illegible] |
| Q9R0P5 | Q9R0P5 | Destrin | *Dstn* | 2.71663 | [illegible] |
| Q9R0P9 | Q9R0P9 | Ubiquitin carboxyl-terminal hydrolase isozyme L1 | *Uchl1* | -0.630391 | [illegible] |
| Q9R0Q6;A0A0G2JF52;D3YVI5 | Q9R0Q6;A0A0G2JF52;D3YVI5 | Actin-related protein 2/3 complex subunit 1A | *Arpc1a* | -0.358076 | [illegible] |
| Q9R0X4;Q32MW3 | Q9R0X4;Q32MW3 | Acyl-coenzyme A thioesterase 9, mitochondrial;Acyl-coenz | *Acot9;Acot10* | -0.176777 | [illegible] |
| Q9R0Y5;Q9R0Y5-2;Z4YN97;A0A0A6YXW8 | Q9R0Y5;Q9R0Y5-2 | Adenylate kinase isoenzyme 1 | *Ak1* | -0.176777 | [illegible] |
| Q9R112;F6ZKZ3;H3BLH2 | Q9R112;F6ZKZ3 | Sulfide:quinone oxidoreductase, mitochondrial | *Sqrdl* | -0.366623 | [illegible] |
| Q9R118 | Q9R118 | Serine protease HTRA1 | *Htra1* | -0.208659 | [illegible] |
| Q9R1J0 | Q9R1J0 | Sterol-4-alpha-carboxylate 3-dehydrogenase, decarboxylat | *Nsdhl* | -0.395202 | [illegible] |
| Q9R1P0;E9PW69;E9Q0X0 | Q9R1P0;E9PW69 | Proteasome subunit alpha type-4;Proteasome subunit alpha | *Psma4* | -0.368089 | [illegible] |
| Q9R1Q6;A0A0N4SUM7;A0A0N4SUY1;A0A0N4SVT4;A0A0N4S | Q9R1Q6 | Transmembrane protein 176B | *Tmem176b* | -0.380378 | [illegible] |
| Q9WTI7-3;Q9WTI7;Q9WTI7-4;Q5ND45;O88329 | Q9WTI7-3;Q9WTI7;Q9WTI7-4 | Unconventional myosin-Ic | *Myo1c* | -0.226375 | [illegible] |
| Q9WTX5;E9PUV4 | Q9WTX5 | S-phase kinase-associated protein 1 | *Skp1* | -0.250932 | [illegible] |
| Q9WTX6;Q3TPM3 | Q9WTX6;Q3TPM3 | Cullin-1 | *Cul1* | -0.239776 | [illegible] |
| Q9WU28;E9PVG3;E9PZ62;E9Q093;E9Q5Q8;H7BWX1 | Q9WU28;E9PVG3;E9PZ62;E9Q093 | Prefoldin subunit 5 | *Pfdn5* | -0.514995 | [illegible] |
| Q9WU78;Q9WU78-3;Q9WU78-2 | Q9WU78;Q9WU78-3 | Programmed cell death 6-interacting protein | *Pdcd6ip* | -0.562517 | [illegible] |
| Q9WUA2;A0A087WS80;A0A087WPV4;A0A087WQ15 | Q9WUA2 | Phenylalanine--tRNA ligase beta subunit | *Farsb* | -0.176777 | [illegible] |
| Q9WUD1 | Q9WUD1 | STIP1 homology and U box-containing protein 1 | *Stub1* | -0.302237 | [illegible] |
| Q9WUK2-2;Q9WUK2 | Q9WUK2-2;Q9WUK2 | Eukaryotic translation initiation factor 4H | *Eif4h* | -0.228604 | [illegible] |
| Q9WUL7 | Q9WUL7 | ADP-ribosylation factor-like protein 3 | *Arl3* | -0.176777 | [illegible] |
| Q9WUM3;D3YUG6 | Q9WUM3 | Coronin-1B | *Coro1b* | -0.311864 | [illegible] |
| Q9WUM4;E9PX03;E9PZJ0;E9PVJ1;B9EIZ7;Q920M5-3;Q920M5 | Q9WUM4 | Coronin-1C | *Coro1c* | -0.285515 | [illegible] |
| Q9WUR9;A2ARF6;F6TEU8;A2ARF5 | Q9WUR9;A2ARF6 | Adenylate kinase 4, mitochondrial | *Ak4* | -0.176777 | [illegible] |
| Q9WUU7 | Q9WUU7 | Cathepsin Z | *Ctsz* | 0.184595 | [illegible] |
| Q9WV54;D3Z505 | Q9WV54 | Acid ceramidase;Acid ceramidase subunit alpha;Acid ceram | *Asah1* | -0.513394 | [illegible] |
| Q9WV55;A0A3B2W837 | Q9WV55;A0A3B2W837 | Vesicle-associated membrane protein-associated protein A | *Vapa* | -0.375138 | [illegible] |
| Q9WVA2;Q4FZG7 | Q9WVA2;Q4FZG7 | Mitochondrial import inner membrane translocase subunit T | *Timm8a1;Timm8a2* | 1.13014 | [illegible] |
| Q9WVA4;A0A0A6YXG6 | Q9WVA4;A0A0A6YXG6 | Transgelin-2 | *Tagln2* | 0.450993 | [illegible] |
| Q9WVJ2;E9Q5I9;F6ZQQ3;E9Q0U1;F6PXS6;E9PY93 | Q9WVJ2;E9Q5I9 | 26S proteasome non-ATPase regulatory subunit 13 | *Psmd13* | 0.458703 | [illegible] |
| Q9WVK4 | Q9WVK4 | EH domain-containing protein 1 | *Ehd1* | -0.252226 | [illegible] |
| Q9Z0G0;A0A1D5RML2 | Q9Z0G0 | PDZ domain-containing protein GIPC1 | *Gipc1* | -0.398718 | [illegible] |
| Q9Z0H8;Q9Z0H8-2 | Q9Z0H8;Q9Z0H8-2 | CAP-Gly domain-containing linker protein 2 | *Clip2* | -0.22377 | [illegible] |
| Q9Z0L0 | Q9Z0L0 | Trophoblast glycoprotein | *Tpbg* | -0.176777 | [illegible] |
| Q9Z0N1;A2AAW9;Q9Z0N2 | Q9Z0N1;A2AAW9;Q9Z0N2 | Eukaryotic translation initiation factor 2 subunit 3, X-linked;E | *Eif2s3x;Eif2s3y* | -0.176777 | [illegible] |
| Q9Z0V1-3;Q9Z0V2;Q9Z0V1-2;Q9Z0V1;Q03719 | Q9Z0V1-3;Q9Z0V2;Q9Z0V1-2;Q9Z0V1;Q0 | Potassium voltage-gated channel subfamily D member 3;P | *Kcnd3;Kcnd2;Kcnd1* | -0.332707 | [illegible] |
| Q9Z110-2;Q9Z110;D3Z0B4;H3BKJ8;H3BLE8 | Q9Z110-2;Q9Z110 | Delta-1-pyrroline-5-carboxylate synthase;Glutamate 5-kina | *Aldh18a1* | -0.334027 | [illegible] |
| Q9Z175;E9Q0X7 | Q9Z175;E9Q0X7 | Lysyl oxidase homolog 3 | *Loxl3* | 1.50564 | [illegible] |
| Q9Z1A1;B8JJG7;B8JJG8;F6QJV5;B8JJG9 | Q9Z1A1;B8JJG7;B8JJG8;F6QJV5 | | *Tfg* | -0.253837 | [illegible] |
| Q9Z1F9;H3BLR3;H3BLM2;A2BH29 | Q9Z1F9 | SUMO-activating enzyme subunit 2 | *Uba2* | -0.321869 | [illegible] |
| Q9Z1G3;A0A2I3BPD0;D3Z5W3;Q99L60-3;Q99L60;Q99L60-2 | Q9Z1G3 | V-type proton ATPase subunit C 1 | *Atp6v1c1* | -0.176777 | [illegible] |
| Q9Z1J3;V9GXS4;A6QRH3;F7CZD1;F6TXD3 | Q9Z1J3 | Cysteine desulfurase, mitochondrial | *Nfs1* | -0.176777 | [illegible] |
| Q9Z1K5;H9KV21;H7BWY7 | Q9Z1K5;H9KV21;H7BWY7 | E3 ubiquitin-protein ligase ARIH1 | *Arih1* | 1.30462 | [illegible] |
| Q9Z1Q2 | Q9Z1Q2 | Abhydrolase domain-containing protein 16A | *Abhd16a* | -0.644103 | [illegible] |
| Q9Z1Q5 | Q9Z1Q5 | Chloride intracellular channel protein 1 | *Clic1* | -0.183759 | [illegible] |
| Q9Z1Q9;G3UY93;G3UZ22;G3UYW2;G3UZX1 | Q9Z1Q9;G3UY93 | Valine--tRNA ligase | *Vars* | 1.33401 | [illegible] |
| Q9Z1T2 | Q9Z1T2 | Thrombospondin-4 | *Thbs4* | 3.79254 | [illegible] |
| Q9Z1W8 | Q9Z1W8 | Potassium-transporting ATPase alpha chain 2 | *Atp12a* | -0.303496 | [illegible] |
| Q9Z204-2;Q9Z204;Q9Z204-4;Q9Z204-3;Q9Z204-5;A0A2I3BR | Q9Z204-2;Q9Z204;Q9Z204-4;Q9Z204-3;Q | Heterogeneous nuclear ribonucleoproteins C1/C2 | *Hnrnpc* | -0.769507 | [illegible] |
| Q9Z247 | Q9Z247 | Peptidyl-prolyl cis-trans isomerase FKBP9 | *Fkbp9* | 0.467241 | [illegible] |
| Q9Z2D6;Q9Z2D6-2 | Q9Z2D6;Q9Z2D6-2 | Methyl-CpG-binding protein 2 | *Mecp2* | -0.306187 | [illegible] |

| | | | | | | | | | | | | | | | | | | | |
|---|---|---|---|---|---|---|---|---|---|---|---|---|---|---|---|---|---|---|---|
| Q9Z2I8;Q9Z2I8-2;A0A0N4SWD1 | Q9Z2I8;Q9Z2I8-2 | Succinyl-CoA ligase [GDP-forming] subunit beta, mitochonc | *Suclg2* | -0.176777 | -0.176777 | -0.176777 | -0.176777 | -0.176777 | -0.176777 | -0.176777 | 5.48008 | -0.176777 | -0.176777 | -0.176777 | -0.176777 | -0.176777 | -0.176777 | -0.176777 | -0.176777 |
| Q9Z2I9 | Q9Z2I9 | Succinyl-CoA ligase [ADP-forming] subunit beta, mitochond | *Sucla2* | -0.176777 | -0.176777 | -0.176777 | 5.48008 | -0.176777 | -0.176777 | -0.176777 | -0.176777 | -0.176777 | -0.176777 | -0.176777 | -0.176777 | -0.176777 | -0.176777 | -0.176777 | -0.176777 |
| Q9Z2U0;A0A338P7D7;Q9CWH6 | Q9Z2U0;A0A338P7D7 | Proteasome subunit alpha type-7 | *Psma7* | -0.25366 | -0.25366 | -0.25366 | -0.25366 | -0.25366 | -0.25366 | -0.25366 | -0.25366 | -0.25366 | 4.04492 | -0.25366 | -0.25366 | -0.25366 | -0.25366 | -0.25366 | -0.25366 |
| Q9Z2U1;A0A0G2JF97 | Q9Z2U1 | Proteasome subunit alpha type-5 | *Psma5* | -0.252291 | -0.252291 | -0.252291 | -0.252291 | 3.31129 | -0.252291 | -0.252291 | -0.252291 | -0.252291 | 4.25745 | -0.252291 | -0.252291 | -0.252291 | -0.252291 | -0.252291 | -0.252291 |
| Q9Z2X1;Q9Z2X1-2;J3QM80;J3QMT0;J3QNH2;J3QMV8;J3QP | Q9Z2X1;Q9Z2X1-2;J3QM80;J3QMT0 | Heterogeneous nuclear ribonucleoprotein F;Heterogeneous | *Hnrnpf* | -0.345705 | -0.345705 | -0.345705 | -0.345705 | -0.345705 | -0.345705 | 0.350202 | -0.345705 | -0.345705 | -0.345705 | -0.345705 | -0.345705 | -0.345705 | -0.345705 | -0.345705 | 2.07915 |
| S4R1P5;Q91ZU6;Q91ZU6-4;E9Q9X1;Q91ZU6-2;A0A087WSP0; | S4R1P5;Q91ZU6;Q91ZU6-4;E9Q9X1;Q91Z | Dystonin | *Dst* | -0.427912 | -0.427912 | -0.427912 | -0.427912 | -0.427912 | -0.427912 | 2.70813 | -0.427912 | -0.427912 | -0.427912 | -0.427912 | -0.427912 | 0.42078 | 0.289254 | -0.427912 | -0.427912 |

| | | | | | | | | | | | | | | | | | | | | | | | | | | | |
|---|---|---|---|---|---|---|---|---|---|---|---|---|---|---|---|---|---|---|---|---|---|---|---|---|---|---|---|
| *Suclg2* | -0.176777 | -0.176777 | -0.176777 | -0.176777 | -0.176777 | -0.176777 | -0.176777 | -0.176777 | -0.176777 | -0.176777 | -0.176777 | -0.176777 | -0.176777 | -0.176777 | -0.176777 | -0.176777 | 18 | 44.1 | 44.1 | 46.839 | 132.23 | 5594100000 | 52 | 0 | 0.830946 | 0 | 0 |
| *Sucla2* | -0.176777 | -0.176777 | -0.176777 | -0.176777 | -0.176777 | -0.176777 | -0.176777 | -0.176777 | -0.176777 | -0.176777 | -0.176777 | -0.176777 | -0.176777 | -0.176777 | -0.176777 | -0.176777 | 18 | 37.1 | 37.1 | 50.113 | 50.733 | 5598100000 | 35 | 0 | 0.830549 | 0 | 0 |
| *Psma7* | -0.25366 | -0.25366 | -0.25366 | 3.56488 | -0.25366 | -0.25366 | -0.25366 | -0.25366 | -0.25366 | -0.25366 | -0.25366 | -0.25366 | -0.25366 | -0.25366 | -0.25366 | -0.25366 | 15 | 52 | 52 | 27.855 | 97.181 | 8683200000 | 56 | 0.00363454 | 0.995851 | 0.00818328 | 0.0106479 |
| *Psma5* | -0.252291 | -0.252291 | -0.252291 | -0.252291 | -0.252291 | -0.252291 | -0.252291 | -0.252291 | -0.252291 | -0.252291 | -0.252291 | -0.252291 | -0.252291 | -0.252291 | -0.252291 | -0.252291 | 11 | 54.8 | 54.8 | 26.411 | 323.31 | 12490000000 | 31 | 0.433155 | 0.780185 | -0.563717 | -0.930949 |
| *Hnrnpf* | 1.43207 | -0.345705 | -0.345705 | -0.345705 | -0.345705 | 0.961188 | 4.51142 | -0.345705 | -0.345705 | -0.345705 | -0.345705 | -0.345705 | -0.345705 | -0.345705 | -0.345705 | -0.345705 | 10 | 34.5 | 28 | 45.729 | 265.16 | 6988700000 | 35 | 0.19385 | 0.857812 | 0.355245 | 0.478913 |
| *Dst* | -0.427912 | 0.899356 | -0.427912 | -0.427912 | -0.427912 | -0.105367 | -0.427912 | -0.427912 | 1.25099 | 0.810671 | -0.427912 | -0.427912 | 3.99607 | -0.427912 | -0.427912 | -0.427912 | 90 | 13.8 | 0.3 | 870.52 | 323.31 | 7813500000 | 162 | 0.0653401 | 0.937407 | 0.0399554 | 0.179491 |

**Supplementary Table 2.** Tools used to predict extracellular or secreted proteins

| Software | Features Identified | Methods |
|---|---|---|
| **LAGO** | Previously reported subcellular localisation | Gene Ontology Cell Component Annotation |
| **SignalP** | Classical signal peptide prediction | Artificial Neural Networks |
| **TargetP** | Classical signal peptide or mitochondrial transit peptide prediction | Artificial Neural Networks |
| **SecretomeP** | Non-classical protein secretion prediction | Artificial Neural Networks |
| **DeepLoc** | Subcellular localisation prediction | Artificial Neural Networks |
| **ProtComp** | Subcellular localisation prediction | Artificial Neural Networks, protein homology, prediction of functional peptide sequences, comparison with a database of multi-located proteins |
| **DeepTMHMM** | Prediction of the topology of both alpha-helical and beta-barrel transmembrane proteins | Artificial Neural Networks |

**Supplementary Table 3**. Table showing the individual and combined results from the protein secretion/localisation prediction analysis of all proteins differentially secreted by astrocytes treated with oligomeric Aβ. Where appropriate the significance cut-off was set to 0.05.

| Gene Name | Up- or Downregulated ? | SignalP Prediction | TargetP Prediction | Non-classically Secreted with SecretomeP | DeepLoc Localization Prediction | GO Annotation to 'Extracellular Space' or 'Extracellular Region' | ProtComp_Integral Prediction of Protein Location (possible additional locations) | Contains TM Helix? | Found in Vesiclepedia? | Predicted as Secreted or Extracellular by At Least One Tool |
|---|---|---|---|---|---|---|---|---|---|---|
| Actb | Up | NO_SP | OTHER | No | Cytoplasm\|Nucleus | No | Cytoplasmic (cytoplasm and nucleus) | No | Yes | Yes |
| Actc1 | Up | NO_SP | OTHER | No | Cytoplasm\|Nucleus | No | Nuclear (cytoplasm and nucleus) | No | Yes | Yes |
| Actg1 | Up | NO_SP | OTHER | No | Cytoplasm\|Nucleus | Yes | Plasma memberane (cytoplasm and nucleus) | No | Yes | Yes |
| Aldh1l1 | Up | NO_SP | OTHER | No | Cytoplasm | No | Cytoplasmic (cytoplasm and nucleus) | No | Yes | Yes |
| Apoe | Up | SP | SP | Yes | Extracellular | Yes | Extracellular (Secreted) | No | Yes | Yes |
| Bcan | Down | SP | SP | No | Extracellular | Yes | Membrane bound Extracellular (Secreted) | No | No | Yes |
| Ccar2 | Down | NO_SP | OTHER | No | Cytoplasm\|Nucleus | No | Cytoplasmic | No | No | Yes |
| Chchd3 | Down | NO_SP | OTHER | Yes | Mitochondrion | No | Mitochondrial | No | Yes | Yes |
| Chga | Down | SP | SP | No | Extracellular | Yes | Extracellular (Secreted) (cytoplasm and secreted) | No | No | Yes |
| Col1a1 | Up | SP | SP | No | Extracellular | Yes | Extracellular (Secreted) | No | Yes | Yes |
| Col1a2 | Up | SP | SP | No | Extracellular | Yes | Membrane bound Extracellular (Secreted) | No | Yes | Yes |
| Cttn | Down | NO_SP | OTHER | No | Cytoplasm | No | Membrane bound Mitochondrial | No | Yes | Yes |
| Dysf | Down | NO_SP | OTHER | No | Cell membrane\|Endoplasmic reticulum | No | Plasma membrane (cytoplasm and nucleus and membrane) | Yes | Yes | Yes |
| H6pd | Down | SP | SP | Yes | Endoplasmic reticulum | No | Membrane bound Mitochondrial | Yes | Yes | Yes |
| Hnrnph2 | Down | NO_SP | OTHER | No | Cytoplasm\|Nucleus | No | Nuclear | No | Yes | Yes |
| Irgq | Down | NO_SP | OTHER | No | Cytoplasm\|Mitochondrion | No | Membrane bound Extracellular (Secreted) | No | No | Yes |
| Krt76 | Down | NO_SP | OTHER | No | Cytoplasm | No | Cytoplasmic (cytoplasm and nucleus) | No | Yes | Yes |
| Mpp6 | Down | NO_SP | OTHER | No | Cytoplasm\|Cell membrane | No | Plasma membrane (cytoplasm and nucleus and membrane) | No | Yes | Yes |
| Ndufa2 | Down | NO_SP | mTP | Yes | Mitochondrion | No | Mitochondrial | No | Yes | Yes |
| Parva | Up | NO_SP | OTHER | Yes | Cytoplasm\|Cell membrane | No | Membrane bound Mitochondrial | No | Yes | Yes |
| Pcolce | Up | SP | SP | No | Extracellular | Yes | Membrane bound Extracellular (Secreted) | No | Yes | Yes |
| Pdia3 | Up | SP | SP | No | Endoplasmic reticulum | Yes | Membrane bound Endoplasmic reticulum (endoplasmic reticulum and vacuole) | No | Yes | Yes |
| Pebp1 | Up | NO_SP | OTHER | No | Cytoplasm\|Nucleus | Yes | Cytoplasmic | No | Yes | Yes |
| Pfkp | Down | NO_SP | OTHER | No | Cytoplasm | No | Cytoplasmic | No | Yes | Yes |
| Ppia | Up | NO_SP | OTHER | No | Cytoplasm | Yes | Mitochondrial (cytoplasm and secreted) | No | Yes | Yes |
| Ppib | Up | SP | SP | Yes | Endoplasmic reticulum | No | ›rane bound Endoplasmic reticulum (cytoplasm and secreted and endoplasmic retic | No | Yes | Yes |
| Prdx1 | Up | NO_SP | OTHER | No | Cytoplasm | No | Cytoplasmic (cytoplasm and secreted) | No | Yes | Yes |
| Prep | Up | NO_SP | OTHER | No | Cytoplasm | No | Cytoplasmic | No | Yes | Yes |
| Prkaa1 | Down | NO_SP | OTHER | No | Cytoplasm | No | Membrane bound Mitochondrial (cytoplasm and nucleus) | No | Yes | Yes |
| Ptgds | Up | SP | SP | Yes | Extracellular | Yes | Extracellular (Secreted) | No | Yes | Yes |
| Rab9a | Down | NO_SP | OTHER | No | Lysosome/Vacuole | No | Vacuolar (cytoplasm and Golgi and endoplasmic reticulum) | No | No | No |
| Rell1 | Down | SP | SP | Yes | Cell membrane\|Lysosome/Vacuole | No | Plasma membrane | Yes | Yes | Yes |
| Rpl32 | Up | NO_SP | OTHER | No | Cytoplasm | No | Cytoplasmic | No | Yes | Yes |
| Rps7 | Down | NO_SP | OTHER | Yes | Cytoplasm | No | Mitochondrial (cytoplasm and nucleus) | No | Yes | Yes |
| Sarnp | Down | NO_SP | OTHER | No | Nucleus | No | Nuclear | No | No | No |
| Scg5 | Down | SP | SP | Yes | Extracellular | Yes | Extracellular (Secreted) | No | No | Yes |
| Sept7 | Up | NO_SP | OTHER | No | Cytoplasm | No | Cytoplasmic | No | Yes | Yes |
| Serpina3n | Up | SP | SP | Yes | Extracellular | Yes | Membrane bound Extracellular (Secreted) (cytoplasm and nucleus) | No | No | Yes |
| Snx5 | Down | NO_SP | OTHER | No | Cytoplasm\|Lysosome/Vacuole\|Golgi apparatus | No | Cytoplasmic | No | Yes | Yes |
| Sod3 | Up | SP | SP | No | Extracellular | Yes | Membrane bound Extracellular (Secreted) | No | No | Yes |
| Spock2 | Down | SP | SP | No | Extracellular | Yes | Extracellular (Secreted) | No | Yes | Yes |
| Tnr | Down | SP | SP | No | Extracellular | No | Membrane bound Extracellular (Secreted) | No | No | Yes |
| Tpm4 | Up | NO_SP | OTHER | No | Cytoplasm | Yes | Cytoplasmic | No | Yes | Yes |
| Trp53bp1 | Up | NO_SP | OTHER | No | Nucleus | No | Membrane bound Extracellular (Secreted) (cytoplasm and nucleus) | No | No | No |
| Vim | Up | NO_SP | OTHER | Yes | Cytoplasm | No | Cytoplasmic (cytoplasm and nucleus) | No | Yes | Yes |

**Supplementary Table 4.** Functional Annotation Clustering of proteins differentially up-regulated in the media of astrocytes exposed to oligomeric Aβ. The enrichment significance threshold was set at the p-value < 0.05. The analysis was performed using Database for Annotation, Visualisation and Integration Discovery (DAVID) software.

| **Annotation Cluster 1** | **Enrichment Score: 3.5738258846577113** | | | | | | |
|---|---|---|---|---|---|---|---|
| **Category** | **Term** | **Count** | **PValue** | **Genes** | **Fold Enrichment** | **FDR** | **-Log10(FDR)** |
| REACTOME_PATHWAY | R-MMU-1650814~Collagen biosynthesis and modifying enzymes | 4 | 4.76E-05 | Pcolce, Ppib, Col1a2, Col1a1 | 53.24277908 | 0.007086098 | 2.149592842 |
| REACTOME_PATHWAY | R-MMU-1474290~Collagen formation | 4 | 1.24E-04 | Pcolce, Ppib, Col1a2, Col1a1 | 38.66439909 | 0.008226881 | 2.0847648 |
| REACTOME_PATHWAY | R-MMU-1474244~Extracellular matrix organization | 4 | 0.003226 | Pcolce, Ppib, Col1a2, Col1a1 | 12.58840901 | 0.043702621 | 1.359492514 |

| **Annotation Cluster 2** | **Enrichment Score: 3.3762769282248994** | | | | | | |
|---|---|---|---|---|---|---|---|
| **Category** | **Term** | **Count** | **PValue** | **Genes** | **Fold Enrichment** | **FDR** | **-Log10(FDR)** |
| GOTERM_CC_DIRECT | GO:0005884~actin filament | 4 | 1.67E-05 | Actb, Tpm4, Actg1, Actc1 | 76.88436633 | 3.68E-04 | 3.434080973 |
| KEGG_PATHWAY | mmu05410:Hypertrophic cardiomyopathy | 4 | 6.82E-05 | Actb, Tpm4, Actg1, Actc1 | 46.20555915 | 0.00136949 | 2.863441148 |
| KEGG_PATHWAY | mmu05414:Dilated cardiomyopathy | 4 | 7.75E-05 | Actb, Tpm4, Actg1, Actc1 | 44.26006192 | 0.00136949 | 2.863441148 |
| REACTOME_PATHWAY | R-MMU-9012999~RHO GTPase cycle | 5 | 0.001263 | Actb, Tpm4, 22352, Actg1, Actc1 | 9.712349054 | 0.026882927 | 1.570523447 |
| REACTOME_PATHWAY | R-MMU-194315~Signaling by Rho GTPases | 5 | 0.006818 | Actb, Tpm4, 22352, Actg1, Actc1 | 6.086599557 | 0.067575743 | 1.170209174 |
| REACTOME_PATHWAY | R-MMU-9716542~Signaling by Rho GTPases, Miro GTPases and RHOBTB3 | 5 | 0.007256 | Actb, Tpm4, 22352, Actg1, Actc1 | 5.979030788 | 0.067575743 | 1.170209174 |

| **Annotation Cluster 3** | **Enrichment Score: 3.177062857115894** | | | | | | |
|---|---|---|---|---|---|---|---|
| **Category** | **Term** | **Count** | **PValue** | **Genes** | **Fold Enrichment** | **FDR** | **-Log10(FDR)** |
| KEGG_PATHWAY | mmu04510:Focal adhesion | 5 | 2.70E-05 | Actb, Actg1, Parva, Col1a2, Col1a1 | 25.76412918 | 0.00136949 | 2.863441148 |
| KEGG_PATHWAY | mmu04611:Platelet activation | 4 | 1.88E-04 | Actb, Actg1, Col1a2, Col1a1 | 32.84926471 | 0.002487942 | 2.604159678 |
| KEGG_PATHWAY | mmu05205:Proteoglycans in cancer | 4 | 8.00E-04 | Actb, Actg1, Col1a2, Col1a1 | 20.02240896 | 0.00848226 | 2.071488443 |
| WIKIPATHWAYS | WP85~Focal adhesion | 4 | 0.002663 | Actb, Actg1, Col1a2, Col1a1 | 12.91992945 | 0.071911894 | 1.143199275 |
| REACTOME_PATHWAY | R-MMU-9006934~Signaling by Receptor Tyrosine Kinases | 4 | 0.012073 | Actb, Actg1, Col1a2, Col1a1 | 7.807234432 | 0.085662199 | 1.067210782 |

| **Annotation Cluster 4** | **Enrichment Score: 2.715644829769168** | | | | | | |
|---|---|---|---|---|---|---|---|
| **Category** | **Term** | **Count** | **PValue** | **Genes** | **Fold Enrichment** | **FDR** | **-Log10(FDR)** |
| REACTOME_PATHWAY | R-MMU-446353~Cell-extracellular matrix interactions | 3 | 2.21E-04 | Actb, Actg1, Parva | 128.2030075 | 0.008226881 | 2.0847648 |
| REACTOME_PATHWAY | R-MMU-446728~Cell junction organization | 3 | 0.002761 | Actb, Actg1, Parva | 36.35607676 | 0.041137376 | 1.385763412 |
| REACTOME_PATHWAY | R-MMU-1500931~Cell-Cell communication | 3 | 0.004504 | Actb, Actg1, Parva | 28.32392027 | 0.052336636 | 1.281194193 |
| GOTERM_CC_DIRECT | GO:0005925~focal adhesion | 3 | 0.004997 | Actb, Actg1, Parva | 27.10173913 | 0.03053678 | 1.515176758 |

| **Annotation Cluster 5** | **Enrichment Score: 1.9487251339780585** | | | | | | |
|---|---|---|---|---|---|---|---|
| **Category** | **Term** | **Count** | **PValue** | **Genes** | **Fold Enrichment** | **FDR** | **-Log10(FDR)** |
| REACTOME_PATHWAY | R-MMU-5674135~MAP2K and MAPK activation | 3 | 9.93E-04 | Actb, Pebp1, Actg1 | 60.89642857 | 0.024650663 | 1.608171395 |
| REACTOME_PATHWAY | R-MMU-5673001~RAF/MAP kinase cascade | 3 | 0.036482 | Actb, Pebp1, Actg1 | 9.404853833 | 0.159090556 | 0.7983556 |
| REACTOME_PATHWAY | R-MMU-5684996~MAPK1/MAPK3 signaling | 3 | 0.039351 | Actb, Pebp1, Actg1 | 9.021693122 | 0.159090556 | 0.7983556 |

**Supplementary Table 5**. Tables showing the ontology (GO) terms for molecular function (MF), biological process (BP), and cell component (CC) significantly (p-value < 0.05) over-represented within the proteins differentially up-regulated in the media of astrocytes exposed to oligomeric Aβ. The analysis was performed using LAGO software.

**LAGO Molecular Function**

| GO ID | Term | P-value | -Log10(P-value) | Uncorrected P-value | Number Annotated | Annotated Genes |
|---|---|---|---|---|---|---|
| GO:0098973 | structural constituent of postsynaptic actin cytoskeleton | 6.00E-04 | 3.2221 | 8.21E-06 | 2 | Actb, Actg1 |
| GO:0042802 | identical protein binding | 7.47E-04 | 3.1265 | 1.02E-05 | 10 | Actb, Actg1, Apoe, Col1a1, Col1a2, Pdia3, Prdx1, Sept7, Tpm4, Vim |
| GO:0016853 | isomerase activity | 8.22E-04 | 3.0852 | 1.13E-05 | 4 | Pdia3, Ppia, Ppib, Ptgds |
| GO:0005198 | structural molecule activity | 1.14E-03 | 2.9449 | 1.56E-05 | 6 | Actb, Actg1, Col1a1, Col1a2, Rpl32, Vim |
| GO:0005200 | structural constituent of cytoskeleton | 2.14E-03 | 2.6696 | 2.93E-05 | 3 | Actb, Actg1, Vim |
| GO:0048407 | platelet-derived growth factor binding | 3.94E-03 | 2.4043 | 5.40E-05 | 2 | Col1a1, Col1a2 |
| GO:0016209 | antioxidant activity | 4.62E-03 | 2.3358 | 6.32E-05 | 3 | Apoe, Prdx1, Sod3 |
| GO:0016018 | cyclosporin A binding | 4.66E-03 | 2.3320 | 6.38E-05 | 2 | Ppia, Ppib |
| GO:0005515 | protein binding | 7.15E-03 | 2.1457 | 9.79E-05 | 18 | Actb, Actc1, Actg1, Aldh1l1, Apoe, Col1a1, Col1a2, Parva, Pdia3, Pebp1, Ppia, Ppib, Prdx1, Sept7, Sod3, Tpm4, Trp53bp1, Vim |
| GO:0099186 | structural constituent of postsynapse | 1.13E-02 | 1.9470 | 1.55E-04 | 2 | Actb, Actg1 |
| GO:0042277 | peptide binding | 1.43E-02 | 1.8441 | 1.96E-04 | 4 | Apoe, Ppia, Ppib, Prep |
| GO:0005488 | binding | 2.07E-02 | 1.6842 | 2.83E-04 | 21 | Actb, Actc1, Actg1, Aldh1l1, Apoe, Col1a1, Col1a2, Parva, Pcolce, Pdia3, Pebp1, Ppia, Ppib, Prdx1, Prep, Ptgds, Sept7, Sod3, Tpm4, Trp53bp1, Vim |
| GO:0098918 | structural constituent of synapse | 2.57E-02 | 1.5898 | 3.52E-04 | 2 | Actb, Actg1 |
| GO:0033218 | amide binding | 2.96E-02 | 1.5294 | 4.05E-04 | 4 | Apoe, Ppia, Ppib, Prep |
| GO:0003755 | peptidyl-prolyl cis-trans isomerase activity | 3.51E-02 | 1.4549 | 4.81E-04 | 2 | Ppia, Ppib |
| GO:0030020 | extracellular matrix structural constituent conferring tensile strength | 4.14E-02 | 1.3832 | 5.67E-04 | 2 | Col1a1, Col1a2 |
| GO:0016859 | cis-trans isomerase activity | 4.14E-02 | 1.3832 | 5.67E-04 | 2 | Ppia, Ppib |

**LAGO Biological Process**

| GO ID | Term | P-value | -Log10(P-value) | Uncorrected P-value | Number Annotated | Annotated Genes |
|---|---|---|---|---|---|---|
| GO:0097435 | supramolecular fiber organization | 1.68E-03 | 2.7747 | 3.13E-06 | 7 | Actc1, Actg1, Apoe, Col1a1, Col1a2, Tpm4, Vim |
| GO:0042221 | response to chemical | 1.15E-02 | 1.9394 | 2.14E-05 | 13 | p |
| GO:0070887 | cellular response to chemical stimulus | 1.89E-02 | 1.7225 | 3.53E-05 | 11 | Actb, Actg1, Col1a1, Col1a2, Parva, Pcolce, Pdia3, Ppib, Prdx1, Sod3, Vim |
| GO:0043589 | skin morphogenesis | 3.42E-02 | 1.4661 | 6.38E-05 | 2 | Col1a1, Col1a2 |
| GO:0048524 | positive regulation of viral process | 3.51E-02 | 1.4546 | 6.55E-05 | 3 | Apoe, Ppia, Ppib |
| GO:1901700 | response to oxygen-containing compound | 3.65E-02 | 1.4373 | 6.82E-05 | 8 | Actb, Col1a1, Col1a2, Pdia3, Prdx1, Serpina3n, Sod3, Vim |
| GO:0032989 | cellular component morphogenesis | 4.56E-02 | 1.3407 | 8.51E-05 | 7 | Actb, Actc1, Actg1, Apoe, Parva, Sept7, Vim |
| GO:1901701 | cellular response to oxygen-containing compound | 4.88E-02 | 1.3118 | 9.10E-05 | 7 | Actb, Col1a1, Col1a2, Pdia3, Prdx1, Sod3, Vim |

**LAGO Cellular Component**

| GO ID | Term | P-value | -Log10(P-value) | Uncorrected P-value | Number Annotated | Annotated Genes |
|---|---|---|---|---|---|---|
| GO:0043209 | myelin sheath | 8.59E-08 | 7.0661 | 6.82E-10 | 7 | Actb, Actg1, Pdia3, Pebp1, Ppia, Prdx1, Sept7 |
| GO:0005615 | extracellular space | 9.73E-05 | 4.0118 | 7.72E-07 | 11 | Actg1, Apoe, Col1a1, Col1a2, Pcolce, Pdia3, Pebp1, Ppia, Ptgds, Serpina3n, Sod3 |
| GO:0005584 | collagen type I trimer | 1.04E-04 | 3.9843 | 8.23E-07 | 2 | Col1a1, Col1a2 |
| GO:0099512 | supramolecular fiber | 1.18E-04 | 3.9296 | 9.33E-07 | 8 | Actb, Actc1, Actg1, Apoe, Col1a1, Col1a2, Tpm4, Vim |
| GO:0099081 | supramolecular polymer | 1.25E-04 | 3.9044 | 9.89E-07 | 8 | Actb, Actc1, Actg1, Apoe, Col1a1, Col1a2, Tpm4, Vim |
| GO:0099080 | supramolecular complex | 1.26E-04 | 3.9009 | 9.97E-07 | 8 | Actb, Actc1, Actg1, Apoe, Col1a1, Col1a2, Tpm4, Vim |
| GO:0032991 | protein-containing complex | 3.31E-04 | 3.4796 | 2.63E-06 | 16 | Actb, Actc1, Actg1, Aldh1l1, Apoe, Col1a1, Col1a2, Parva, Pdia3, Ppia, Ppib, Rpl32, Sept7, Tpm4, Trp53bp1, Vim |
| GO:0044421 | extracellular region part | 3.43E-04 | 3.4646 | 2.72E-06 | 11 | Actg1, Apoe, Col1a1, Col1a2, Pcolce, Pdia3, Pebp1, Ppia, Ptgds, Serpina3n, Sod3 |
| GO:0005737 | cytoplasm | 6.30E-04 | 3.2008 | 5.00E-06 | 21 | Actb, Actc1, Actg1, Aldh1l1, Apoe, Col1a1, Col1a2, Parva, Pdia3, Pebp1, Ppia, Ppib, Prdx1, Prep, Ptgds, Rpl32, Sept7, Sod3, Tpm4, Trp53bp1, Vim |
| GO:0015629 | actin cytoskeleton | 6.41E-04 | 3.1931 | 5.09E-06 | 6 | Actb, Actc1, Actg1, Parva, Sept7, Tpm4 |
| GO:0005884 | actin filament | 8.90E-04 | 3.0504 | 7.07E-06 | 4 | Actb, Actc1, Actg1, Tpm4 |
| GO:0005576 | extracellular region | 2.27E-03 | 2.6431 | 1.81E-05 | 11 | Actg1, Apoe, Col1a1, Col1a2, Pcolce, Pdia3, Pebp1, Ppia, Ptgds, Serpina3n, Sod3 |
| GO:0043679 | axon terminus | 3.46E-03 | 2.4606 | 2.75E-05 | 4 | Actb, Actg1, Pebp1, Sept7 |
| GO:0099513 | polymeric cytoskeletal fiber | 3.88E-03 | 2.4107 | 3.08E-05 | 6 | Actb, Actc1, Actg1, Apoe, Tpm4, Vim |
| GO:0044444 | cytoplasmic part | 4.05E-03 | 2.3925 | 3.21E-05 | 18 | Actb, Actc1, Actg1, Aldh1l1, Apoe, Col1a1, Col1a2, Pdia3, Pebp1, Ppib, Prdx1, Prep, Ptgds, Rpl32, Sept7, Sod3, Tpm4, Vim |
| GO:0043229 | intracellular organelle | 4.76E-03 | 2.3219 | 3.78E-05 | 21 | Actb, Actc1, Actg1, Aldh1l1, Apoe, Col1a1, Col1a2, Parva, Pdia3, Pebp1, Ppia, Ppib, Prdx1, Prep, Ptgds, Rpl32, Sept7, Sod3, Tpm4, Trp53bp1, Vim |
| GO:0044306 | neuron projection terminus | 5.19E-03 | 2.2846 | 4.12E-05 | 4 | Actb, Actg1, Pebp1, Sept7 |
| GO:0005583 | fibrillar collagen trimer | 5.67E-03 | 2.2462 | 4.50E-05 | 2 | Col1a1, Col1a2 |
| GO:0098643 | banded collagen fibril | 5.67E-03 | 2.2462 | 4.50E-05 | 2 | Col1a1, Col1a2 |
| GO:0043226 | organelle | 7.78E-03 | 2.1093 | 6.17E-05 | 21 | Actb, Actc1, Actg1, Aldh1l1, Apoe, Col1a1, Col1a2, Parva, Pdia3, Pebp1, Ppia, Ppib, Prdx1, Prep, Ptgds, Rpl32, Sept7, Sod3, Tpm4, Trp53bp1, Vim |
| GO:0001725 | stress fiber | 8.55E-03 | 2.0683 | 6.78E-05 | 3 | Actb, Sept7, Tpm4 |
| GO:0097517 | contractile actin filament bundle | 8.55E-03 | 2.0683 | 6.78E-05 | 3 | Actb, Sept7, Tpm4 |
| GO:0098871 | postsynaptic actin cytoskeleton | 9.37E-03 | 2.0282 | 7.44E-05 | 2 | Actb, Actg1 |
| GO:0032432 | actin filament bundle | 1.15E-02 | 1.9395 | 9.12E-05 | 3 | Actb, Sept7, Tpm4 |
| GO:0042641 | actomyosin | 1.15E-02 | 1.9395 | 9.12E-05 | 3 | Actb, Sept7, Tpm4 |
| GO:0098644 | complex of collagen trimers | 1.57E-02 | 1.8036 | 1.25E-04 | 2 | Col1a1, Col1a2 |
| GO:0099571 | postsynaptic cytoskeleton | 1.76E-02 | 1.7555 | 1.39E-04 | 2 | Actb, Actg1 |
| GO:0002102 | podosome | 3.59E-02 | 1.4452 | 2.85E-04 | 2 | Actb, Tpm4 |
| GO:0044305 | calyx of Held | 4.15E-02 | 1.3824 | 3.29E-04 | 2 | Actb, Actg1 |

**Supplemental Table 6**. Tables showing the significant (FDR p-value < 0.05) Reactome pathways identified in the proteins differentially up-regulated in the media of astrocytes exposed to oligomeric Aβ. The Reactome pathway analysis was performed using PANTHER v16.0 software.

| Reactome Pathways | Fold Enrichment | Raw P-value | FDR | -Log10(FDR) | Gene Count | Genes Matched to the Pathway |
|---|---|---|---|---|---|---|
| Collagen biosynthesis and modifying enzymes | 57.26 | 8.45E-07 | 6.96E-04 | 3.1574 | 4 | Col1a1, Col1a2, Pcolce, Ppib |
| Crosslinking of collagen fibrils | > 100 | 1.73E-06 | 7.13E-04 | 3.1469 | 3 | Col1a1, Col1a2, Pcolce |
| Cell-extracellular matrix interactions | > 100 | 1.50E-06 | 8.22E-04 | 3.0851 | 3 | Actb, Actg1, Parva |
| Collagen formation | 41.64 | 2.85E-06 | 9.39E-04 | 3.0273 | 4 | Col1a1, Col1a2, Pcolce, Ppib |
| Striated Muscle Contraction | 76.35 | 1.01E-05 | 2.79E-03 | 2.5544 | 3 | Actc1, Tmp4, Vim |
| MAP2K and MAPK activation | 68.71 | 1.37E-05 | 3.22E-03 | 2.4921 | 3 | Actb, Actg1, Pebp1 |
| Assembly of collagen fibrils and other multimei | 44.33 | 4.76E-05 | 9.81E-03 | 2.0083 | 3 | Col1a1, Col1a2, Pcolce |
| Formation of annular gap junctions | > 100 | 7.47E-05 | 1.12E-02 | 1.9508 | 2 | Actb, Actg1 |
| Gap junction degradation | > 100 | 8.82E-05 | 1.12E-02 | 1.9508 | 2 | Actb, Actg1 |
| Cell junction organization | 40.42 | 6.20E-05 | 1.14E-02 | 1.9431 | 3 | Actb, Actg1, Parva |
| Platelet Adhesion to exposed collagen | > 100 | 8.82E-05 | 1.21E-02 | 1.9172 | 2 | Col1a1, Col1a2 |
| GP1b-IX-V activation signalling | > 100 | 1.03E-04 | 1.21E-02 | 1.9172 | 2 | Col1a1, Col1a2 |
| Interaction between L1 and Ankyrins | > 100 | 7.47E-05 | 1.23E-02 | 1.9101 | 2 | Actb, Actg1 |
| Cell-Cell communication | 30.54 | 1.39E-04 | 1.52E-02 | 1.8182 | 3 | Actb, Actg1, Parva |
| Anchoring fibril formation | > 100 | 1.72E-04 | 1.78E-02 | 1.7496 | 2 | Col1a1, Col1a2 |
| MAPK family signalling cascades | 13.62 | 2.02E-04 | 1.96E-02 | 1.7077 | 4 | Actb, Actg1, Pebp1, Sept7 |
| Extracellular matrix organization | 12.42 | 2.86E-04 | 2.62E-02 | 1.5817 | 4 | Col1a1, Col1a2, Pcolce, Ppib |
| RHO GTPases activate IQGAPs | 67.86 | 4.54E-04 | 3.94E-02 | 1.4045 | 2 | Actb, Actg1 |
| MET activates PTK2 signalling | 63.18 | 5.20E-04 | 4.28E-02 | 1.3686 | 2 | Col1a1, Col1a2 |
| Detoxification of Reactive Oxygen Species | 59.11 | 5.89E-04 | 4.63E-02 | 1.3344 | 2 | Prdx1, Sod3 |
| Signalling by Receptor Tyrosine Kinases | 8.85 | 1.01E-03 | 4.89E-02 | 1.3107 | 4 | Actb, Actg1, Col1a1, Col1a2 |
| RHO GTPases Activate WASPs and WAVEs | 50.9 | 7.82E-04 | 4.96E-02 | 1.3045 | 2 | Actb, Actg1 |
| Adherens junctions interactions | 55.53 | 6.63E-04 | 4.97E-02 | 1.3036 | 2 | Actb, Actg1 |

**Supplementary Table 7**. Table showing the significant (FDR p-value < 0.05) STRING local network clusters identified in the proteins differentially up-regulated in the media of astrocytes treated with oligomeric Aβ. The analysis was performed using STRING software.

| Term Description | Observed Gene Count | Strength | False Discovery Rate P-value | -Log10(FDR) | Matching Proteins in the Network |
|---|---|---|---|---|---|
| Banded collagen fibril, and peptidase m12b, adam-ts2 | 3 | 2.61 | 0.00057 | 3.244125144 | Col1a1,Col1a2,Pcolce |
| Collagen biosynthesis and modifying enzymes | 4 | 1.85 | 0.00089 | 3.050609993 | Col1a1,Col1a2,Pcolce,Ppib |
| Striated Muscle Contraction, and Thick filament | 3 | 1.84 | 0.0133 | 1.876148359 | Tpm4,Vim,Actc1 |
| Detoxification of Reactive Oxygen Species | 2 | 2.17 | 0.0478 | 1.320572103 | Sod3,Prdx1 |

**Supplementary Table 8**. Tables showing the ontology (GO) terms for biological process (BP) and cell component (CC) significantly (p-value < 0.05) over-represented in the proteins differentially down-regulated in the media of astrocytes exposed to oligomeric Aβ. The analysis was performed using LAGO software.

**LAGO Biological Process**

| GO ID | Term | P-value | -Log10(P-value) | Uncorrected P-value | Number Annotated | Annotated genes |
|---|---|---|---|---|---|---|
| GO:00903 | regulation of protein deacetylation | 0.0068004 | 2.167464903 | 1.2275E-05 | 3 | Ccar2, Dysf, Prkaa1 |
| GO:19044 | negative regulation of tubulin deacetylation | 0.0087167 | 2.059647403 | 1.5734E-05 | 2 | Dysf, Prkaa1 |
| GO:00900 | regulation of tubulin deacetylation | 0.0273218 | 1.563490692 | 4.9317E-05 | 2 | Dysf, Prkaa1 |
| GO:00070 | mitochondrion organization | 0.0329163 | 1.482588988 | 5.9416E-05 | 5 | Ccar2, Chchd3, Cttn, Ndufa2, Prkaa1 |
| GO:00150 | protein transport | 0.0336959 | 1.472422939 | 6.0823E-05 | 8 | Chga, Cttn, Dysf, Prkaa1, Rab9, Sarnp, Scg5, Snx5 |
| GO:00158 | peptide transport | 0.04233 | 1.373351732 | 7.6408E-05 | 8 | Chga, Cttn, Dysf, Prkaa1, Rab9, Sarnp, Scg5, Snx5 |
| GO:00900 | tubulin deacetylation | 0.0433965 | 1.362545296 | 7.8333E-05 | 2 | Dysf, Prkaa1 |
| GO:00428 | amide transport | 0.0470165 | 1.327749704 | 8.4867E-05 | 8 | Chga, Cttn, Dysf, Prkaa1, Rab9, Sarnp, Scg5, Snx5 |

**LAGO Cellular Component**

| GOID | Term | P-value | -Log10(P-value) | Uncorrected P-value | Number Annotated | Annotated genes |
|---|---|---|---|---|---|---|
| GO:00725 | perineuronal net | 1.21E-03 | 2.9156 | 1.12E-05 | 2 | Bcan, Tnr |
| GO:00989 | perisynaptic extracellular matrix | 1.70E-03 | 2.7697 | 1.57E-05 | 2 | Bcan, Tnr |
| GO:00995 | synapse-associated extracellular matrix | 2.26E-03 | 2.6450 | 2.10E-05 | 2 | Bcan, Tnr |
| GO:00995 | perisynaptic space | 4.44E-03 | 2.3525 | 4.11E-05 | 2 | Bcan, Tnr |

**Supplementary Table 10**. Table showing the gene-disease association (GDA) scores of 21 proteins differentially secreted by Aβ-treated astrocytes with Alzheimer's disease. The analysis was performed using DisGeNET software.

| Gene | Up- or Downregulated? | Disease | Gene-Disease Association Score |
|---|---|---|---|
| APOE | ↑ | Alzheimer's Disease | 0.7 |
| SERPINA3 | ↑ | Alzheimer's Disease | 0.1 |
| ACTB | ↑ | Alzheimer's Disease | 0.1 |
| ACTG1 | ↑ | Alzheimer's Disease | 0.1 |
| PRKAA1 | ↓ | Alzheimer's Disease | 0.1 |
| PREP | ↑ | Alzheimer's Disease | 0.06 |
| VIM | ↑ | Alzheimer's Disease | 0.05 |
| PRDX1 | ↑ | Alzheimer's Disease | 0.04 |
| PTGDS | ↑ | Alzheimer's Disease | 0.03 |
| CHGA | ↓ | Alzheimer's Disease | 0.03 |
| PEBP1 | ↑ | Alzheimer's Disease | 0.02 |
| BCAN | ↓ | Alzheimer's Disease | 0.02 |
| PPIA | ↑ | Alzheimer's Disease | 0.02 |
| CTTN | ↓ | Alzheimer's Disease | 0.01 |
| PPIB | ↑ | Alzheimer's Disease | 0.01 |
| TNR | ↓ | Alzheimer's Disease | 0.01 |
| TP53BP1 | ↑ | Alzheimer's Disease | 0.01 |
| DYSF | ↓ | Alzheimer's Disease | 0.01 |
| ALDH1L1 | ↑ | Alzheimer's Disease | 0.01 |
| SOD3 | ↑ | Alzheimer's Disease | 0.01 |
| RAB9A | ↓ | Familial Alzheimer Disease (FAD) | 0.01 |

**Supplementary Table 9**. Summary of RNAseq studies used in this study to compare to our astrocyte protein secretome

| **Publication** | **Method** | **Brain Region** | **Cohort** |
|---|---|---|---|
| Mathys et al., 2019 | snRNA-seq | Prefrontal cortex | 24 patients with a spectrum of mild to severe Aβ and other pathologies (AD-pathology) and 24 controls (no-pathology) |
| Grubman et al., 201[illegible] | snRNA-seq | Entorhinal cortex | 6 late-stage (Braak stage V-VI) AD patients and 6 controls |
| Lau et al., 2020 | snRNA-seq | Prefrontal cortex | 12 AD patients and 9 controls |
| Smith et al., 2022 | snRNA-seq | Entorhinal and somatosensory cortex | 6 non-diseased control cases (Braak stage 0–II) and 6 AD cases (Braak stage III–VI) |
| Galea et al., 2022 | Transcriptome data from the MtSINAI, MAYO, and ROSMAP databases | Whole brain | Samples from prefrontal cortex of 101 Controls and 129 AD patients (MtSINAI), temporal cortex of 78 Controls and 82 AD patients (MAYO), and dorsolateral prefrontal cortex from 200 Control cases, 157 MCI cases, and 219 AD cases (ROSMAP) |
| Sadick et al., 2022 | snRNA-seq | Prefrontal cortex | 9 AD patients and 5 controls |
